# FLIM Playground: An interactive, end-to-end graphical user interface for analyzing single cells with fluorescence lifetime imaging microscopy

**DOI:** 10.1101/2025.09.30.679625

**Authors:** Wenxuan Zhao, Kayvan Samimi, Melissa C. Skala, Rupsa Datta

## Abstract

Fluorescence lifetime imaging microscopy (FLIM) is a cellular-resolution molecular imaging technique. Yet, the journey from raw photon decays to biological insight remains fragmented by multi-step data extraction and siloed analyses. This work presents FLIM Playground, the first interactive graphical platform that unifies single-cell FLIM workflows, embeds user checks at each stage, and offers diverse user options. Built in Python and available open-source, FLIM Playground runs on major operating systems as a ready-to-run application and is web deployable. Its Data Extraction section collects and checks field-of-view metadata, calibrates via instrument response function shift or fluorescence lifetime standard, and extracts single-cell fluorescence lifetime features, along with morphology and texture features across channels. Multiple datasets can be merged through an interface that assigns categorical labels. The Data Analysis section provides real-time visual analytic modules for outputs from Data Extraction or user-provided datasets. Lifetime extraction by fitting and phasor were validated by comparison with a commercial software and published results, respectively, and both sections were demonstrated on a FLIM dataset of cancer cell lines to obtain biological insights. By adopting best practices and offering interactivity, FLIM Playground accelerates hypothesis-driven discovery and promotes reproducibility, and its modular design can incorporate new imaging modalities, extraction methods, and analysis modules.

## 1. INTRODUCTION

Fluorescence lifetime imaging microscopy (FLIM) is a popular molecular imaging tool across diverse fields such as cell biology, biophysics, and biomedical research^1–5^. FLIM is sensitive to changes in fluorophore microenvironment including temperature, pH, conformational changes with protein binding, and the presence of quenchers^2,4^. Advancements in instrumentation, detection technologies, acquisition speed, and automated cell-segmentation algorithms^6–9^ have greatly expanded its capabilities, making FLIM more powerful and widely adopted. Together, these developments make single-cell analyses possible that reveal biological heterogeneity^10^ and underscore the need for commensurate progress in analysis tools^11^.

A typical FLIM experiment spans multiple imaging sessions, each containing multiple multi-channel fields of view (FOVs). Each channel—spectral, acquisition, or color—contains multiple cell-level regions of interest (ROIs), defined by ROI masks and composed of pixel sets. At the pixel level, fluorescence decays can be fitted^12–14^ or transformed into phasor features^15–17^. ROI masks or pre-aggregated cell-level decays^18^ enable single-cell mapping of heterogeneity through fit^19^, phasor^20^, morphology, and texture^21^ extraction. The channel level introduces fluorophore-specific inputs, enabling per-channel extraction of single-cell numerical features. The FOV level aggregates information across channels, allowing unique single-cell identifiers to be linked with their imaging context. Finally, the experiment level takes qualitative category labels of FOVs and incorporates them into downstream analysis methods to gain biological insights. Together, pixels, cell ROIs, channels, FOVs, and experiments constitute the hierarchical data levels in FLIM analysis, each representing a distinct unit of information with progressively richer biological semantics.

While a diverse set of tools—both open-source and commercial, ranging from libraries to graphical user interfaces—offers alternative methods and flexibility for FLIM analysis, they typically address only subsets of the data levels. As a result, users often write custom code to reformat inputs, process outputs, and explore alternative methods, leading to a fragmented landscape of FLIM analysis.

FLIM Playground provides a unified framework for integrating diverse methods and input types across data levels, accessible through a cross-platform, interactive, and code-free graphical user interface. It spans the full pipeline: from FOV metadata organization, through calibration with instrument response functions (IRF) or fluorescence lifetime standards, to extraction of single-cell identifiers, numerical features (lifetime fits and phasor, morphology, and texture) and categorical features, followed by visualization and statistical modeling. Embedded validation checks guide users at every step, while interactive widgets and a built-in repertoire of visual-analytic modules encourage hypothesis-driven and iterative exploration of large datasets, including user-provided datasets.

FLIM Playground’s modular architecture can incorporate new imaging modalities, feature extraction methods, and analysis modules as they evolve, and, being open-source and Python-based, other labs can easily adopt and extend the platform and leverage the rich Python package ecosystem.

To ensure reliability, we compared FLIM Playground’s lifetime fits against the widely used commercial SPCImage package^14^ using two-photon autofluorescence FLIM of cells. We further illustrate the capabilities of FLIM Playground on autofluorescence FLIM of cancer cell lines (PANC-1 and MCF7) treated with metabolic inhibitors and on a fluorescence lifetime flow cytometry^18^ dataset of primary human T cells.

## 2. METHODS

FLIM Playground is organized into two main sections: Data Extraction and Data Analysis. Data Extraction derives single-cell features from raw FLIM images, while Data Analysis can process these outputs or directly operate on user-provided datasets for data visualization and statistical analysis.

Both Data Extraction and Data Analysis are designed to focus on three classes of single-cell features in the imaging data: categorical features (discrete cell groups such as treatment, cell line), identifiers (FOV identifiers and unique cell identifiers), and numerical features (quantitative measures such as lifetime and morphology features that capture differences or similarities between groups) (Table 1). At each Data Extraction stage, FLIM Playground targets one feature class and provides interactive interfaces with validation checks and status feedback. Data Analysis builds dedicated interactive components for each feature class, which are shared across the analysis modules it provides, while module-specific components further support a variety of statistical modeling approaches.

**Table 1.**
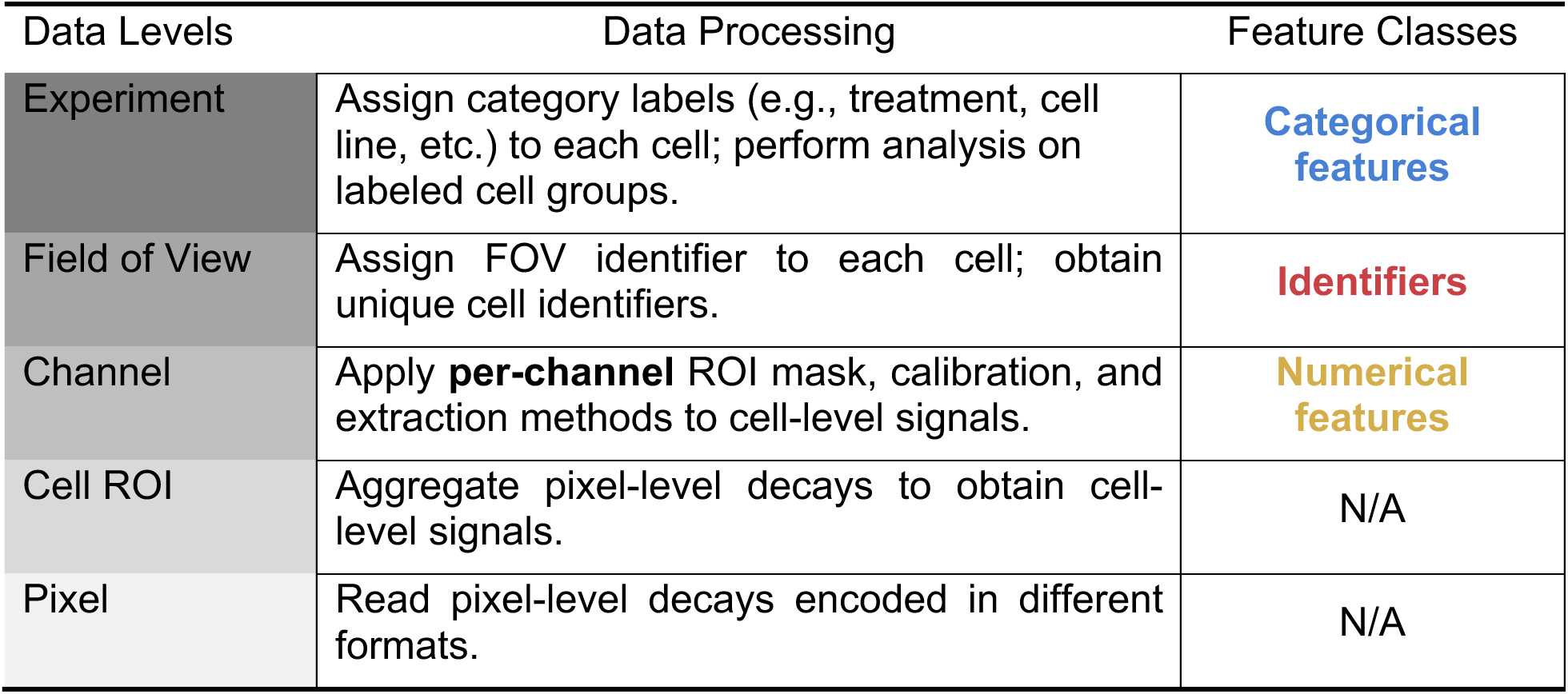
FLIM Playground processes at each data level to extract single-cell features from three feature classes. Each data level in the hierarchy is composed of a set of instances from the level below (e.g., a cell is composed of a set of pixels). The table shows each level in progressively darker gray from bottom (pixel) to top (experiment) to illustrate this hierarchical structure. FLIM Playground manages this complexity by reading pixel-level decays encoded in vendor-specific formats (e.g., Becker & Hickl, PicoQuant), and aggregates the decays that belong to the same cell ROI to obtain the cell-level signals (i.e., cell-level decay and cell intensity image). In this way, we can achieve single-cell resolution at all feature classes (in bold colors). For each acquisition channel, FLIM Playground flexibly handles the per-channel masks, calibration procedures, and methods (e.g., fit, phasor, texture, and morphology) to extract numerical features (yellow). FLIM Playground organizes metadata for FOVs and assigns to cells the FOV they come from and their unique identifiers (red). Category labels (blue) such as treatment and time points are biologically meaningful and FLIM Playground can assist the user in assigning those category labels to each cell and using them to perform data analysis.

FLIM Playground employs the interactive graphical user interface library Streamlit^22^, which supports both local and web deployment. PyInstaller^23^ bundles the software into executables that run locally on major operating systems (ready-to-use executables for Windows and Mac are available in the FLIM Playground GitHub^24^ repository). Figure 1 provides an overview of the software.

**Figure 1.**
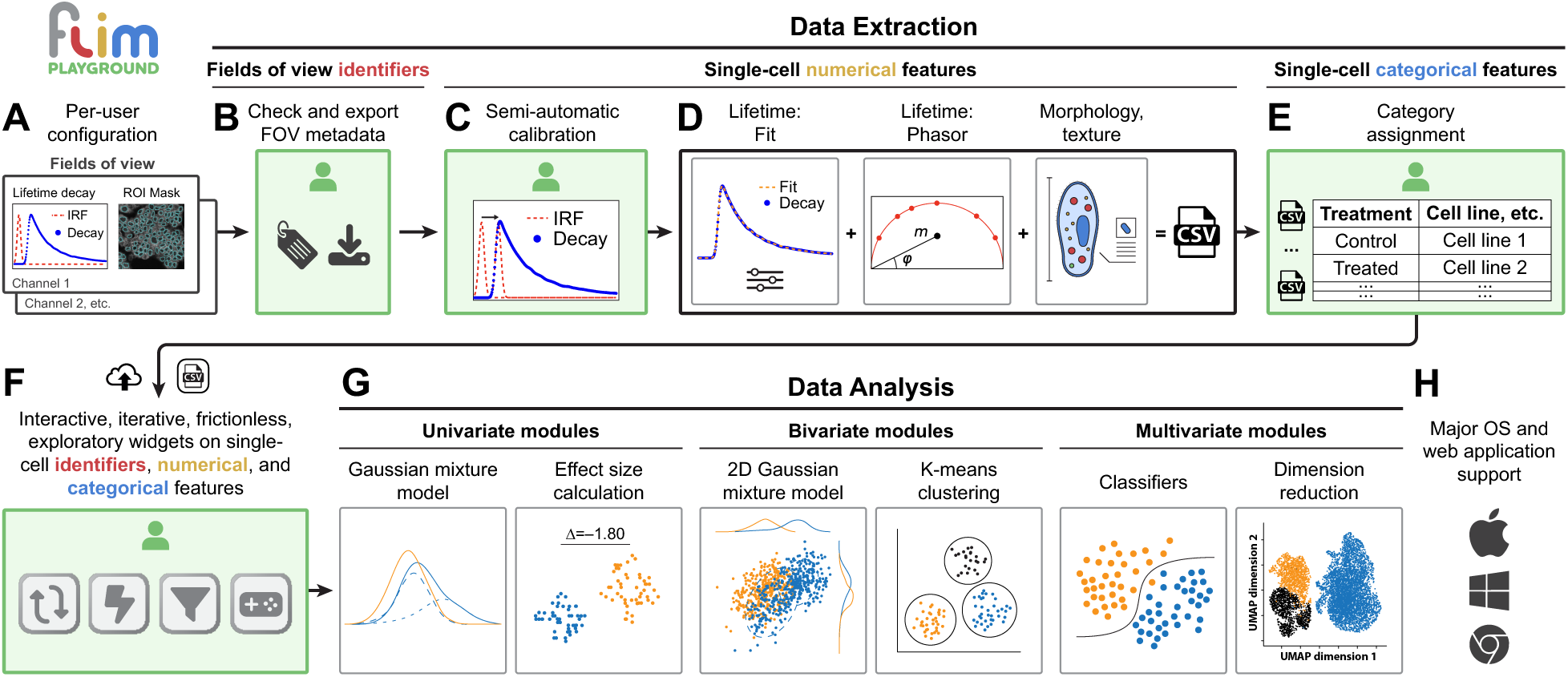
System overview of FLIM Playground. After **(A)** loading a saved configuration, **(B) Data Extraction** collects and validates channel inputs across all fields of views, **(C)** calibrates for IRF shifts or uses fluorescence lifetime standards, and **(D)** extracts single-cell lifetime (fit and phasor), morphology, and texture features in one step. **(E)** To bridge Data Extraction with downstream analysis, FLIM Playground combines multiple single-cell datasets and assists users with assigning category labels to each cell. **(F)** The three feature classes—identifiers (red), numerical features (yellow), and categorical features (blue)—whether extracted through Data Extraction or from user-provided datasets, are transformed into interactive widgets shared by all analysis methods. Icons denote interface attributes: iterative, frictionless, exploratory, and intuitive. **(G) Data Analysis** includes univariate, bivariate, and multivariate modules. **(H)** FLIM Playground is available as a ready-to-run executable for major operating systems and as a web application. Interactive interfaces are indicated by a green background.

### 2.1 Data Extraction produces single-cell features using a modularized framework

The modularized Data Extraction framework in FLIM Playground is *channel-centric*, which enables per-channel customization of masks that might focus on different cell ROIs, calibration procedures, and feature extraction methods (Table 1). These assignments are managed through a configuration interface, so the configuration can be set up once and reused across future analyses or altered as needed. After users specify the number of channels, the interface renders a column for configuring settings within each channel. Then users can assign a name (e.g., fluorophore label), an imaging modality, and a list of numerical feature extractors (e.g., *Lifetime Fit extractor* or *Morphology extractor*) to each channel.

Two imaging modalities are supported. In *FLIM mode*, the signal is time-resolved and corresponds to one of the decay types described below. In *intensity-only mode*, the signal is a spatial intensity map without a lifetime axis (e.g., TIFF/TIF format), enabling mixed-modality analyses within the same field of view and supporting analysis of non-FLIM images.

Each channel can be associated with a subset of feature extractors rather than a fixed choice across all channels. Lifetime-based extractors, fit or phasor, can be applied to FLIM channels, while morphology- and texture-based extractors are available for all imaging modalities. These extractors will be introduced in detail in later sections. This modular design also allows future integration of additional imaging modalities and corresponding feature extractors.

Based on the selected imaging modality and feature extractors, the interface prompts the user to specify file suffixes for each required input file type (e.g., raw decay, ROI mask, calibration file). These are later used during the FOV identification stage. Channels can either share a common ROI mask or use channel-specific masks to target distinct compartments within a cell.

Beyond channel-specific settings, several parameters apply across all channels. Three decay types are supported: (1) 3D/4D decay, consisting of spatial dimensions and the fluorescence decay axis, with an optional spectral channel dimension capturing multiple fluorophores (time-lapse acquisitions are not yet supported). These image data are stored in vendor-specific formats, and FLIM Playground supports those from Becker & Hickl and PicoQuant. (2) 2D decay, where each row of a tabular datasheet corresponds to the fluorescence decay of a single cell, and each column represents a time bin along the decay curve—an example of such data is obtained from fluorescence lifetime flow cytometry^18^; and (3) 3D/4D pixel-prefit decay, representing pixel-level lifetime features pre-fit by open-source or commercial software such as SPCImage. Two calibration methods are available for phasor analysis: either by shifting the IRF or by using a fluorescence lifetime standard with known lifetime^25^. Users may also specify a list of categorical features for extraction, which will later be recognized in the Data Analysis section. A few additional settings are described in the Supplemental Manual, Chapter 3; the most up-to-date version of the FLIM Playground manual is also available online^26^.

The next steps after specifying the reusable configuration are the three stages of Data Extraction, each focusing on one feature class (Figure 2). Supplemental Video 1 shows an example run of the Data Extraction pipeline.

**Figure 2.**
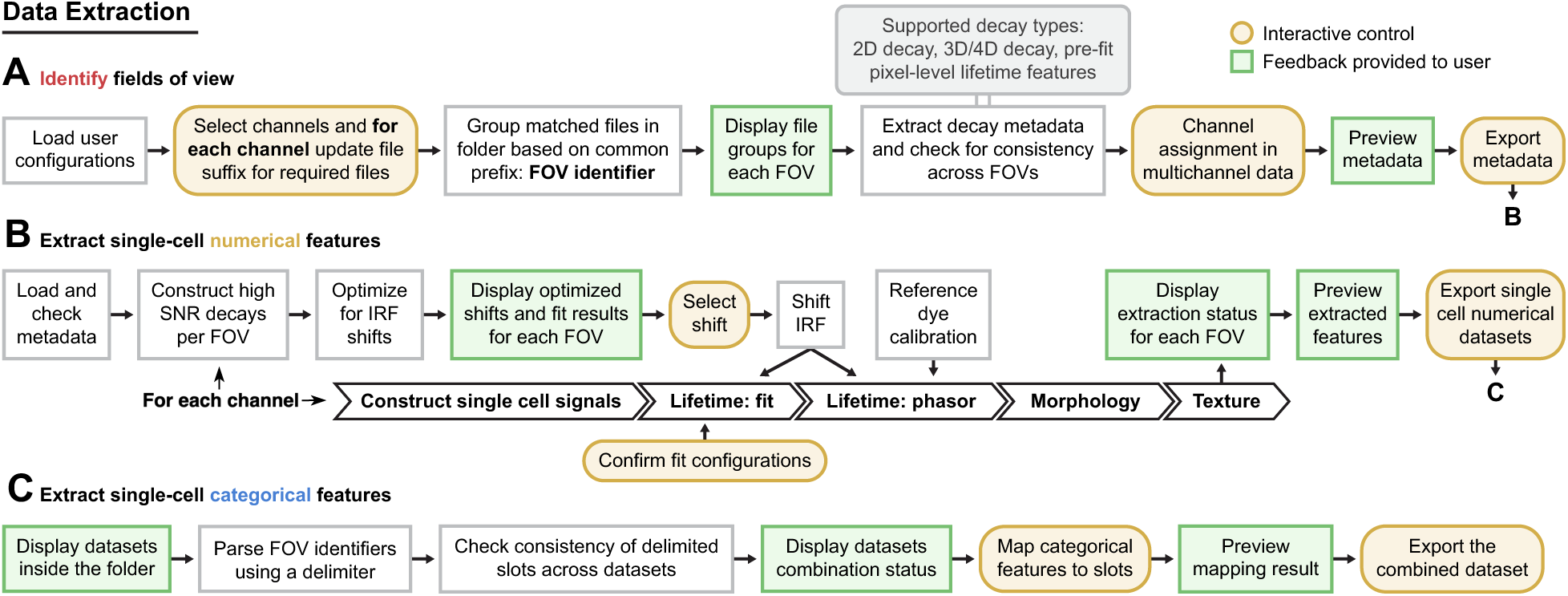
A high-level overview of the Data Extraction workflow. This section is divided into three stages **(A, B, C)**. A short description of steps involved in each stage is shown here, with detailed descriptions provided in following sections. The white boxes are internal processing steps not shown to the user. The yellow oval-shaped boxes are interactive controls user can change. Green boxes are interfaces that provide feedback to the user.

### 2.2 Identify Fields of View

Imaging sessions consist of multiple fields of view, each containing input files associated with potentially multiple channels. Users are prompted to enter the file path to the folder that contains all files necessary for extracting features (Figure 2A, Supp Man. Chapter 4). Per-channel and cross-channel settings are loaded from the configuration, and most can be updated at this stage, including the file suffixes for each input file type of each channel. FLIM Playground recursively searches the folder path for files that end with the first file suffix of the first channel. The prefixes of the matched files are defined as FOV identifiers (i.e., file name minus the first suffix), under the assumption that *all files associated with a FOV share the same prefix.* All other files across channels are then located by appending each suffix to the FOV identifier. The only exception is calibration files, which are not FOV-specific and are therefore searched based solely on their file suffixes. For each FOV, the search status is reported. A status of *Success* indicates that all required files were located, and the FOV is then recorded into a metadata file together with the paths to its files. If a required file type is missing, its type and expected filename are displayed. A status of *Duplicate* is shown when multiple files with the same name are detected.

FLIM Playground then reads the metadata of all decay files. The temporal laser pulse interval and the number of time bins per pulse interval are cross-checked across all channels for consistency. The time bins are also validated against the lifetime axis of the calibration files. In addition, the spatial dimensions of the decay files are cross-checked with the ROI masks to ensure consistency. For each acquisition channel, if the corresponding decay file contains more than one non-empty channel, the user is prompted to select the appropriate channel number. The sdtfile^27^ and ptufile^28^ Python libraries are used to read decay data from Becker & Hickl and PicoQuant files. Channel numbers, FOV spatial dimensions, pulse interval, and time bins are recorded into the metadata file. The imaging modality and feature extractors for each channel are also recorded. A preview of the metadata file is displayed, which can be exported in comma-separated values (CSV) format. The file is simultaneously cached in the system to enable seamless progression to the next stage.

### 2.3 Extract Single-Cell Numerical Features

Using either the cached metadata file from the previous step or a newly uploaded file, FLIM Playground initiates numerical feature extraction as a two-step process: calibration followed by feature extraction (Figure 2B). Calibration is required whenever lifetime fit or phasor analysis is selected, since the measured fluorescence decay is the convolution product of the underlying fluorescence decay with the IRF. Calibration is carried out separately for each channel.

#### 2.3.1 Fit Calibration

In *fit calibration*, IRF time shifts are estimated through reconvolution fitting, where the shift is treated as a free parameter optimized jointly with lifetimes and amplitudes using high signal-to-noise (SNR) decay curves from each FOV (Supp Man. Chapter 5). For each FOV, high SNR decays are constructed by summing the decay curves of all non-background pixels defined by the ROI mask. The brightest cells are selected as high SNR decays if the decay type is 2D decay. Reconvolution fitting models the measured fluorescence decay as a sum of exponential functions—reflecting the characteristic lifetimes of fluorophores—and fits this model after convolution with the IRF to account for temporal broadening introduced during acquisition.

The time axis (**t**) of the exponential function is defined by time bin width (Δ*t*), which is the ratio between the laser pulse period (*T*) and the number of time bins (*N*) (Eq. 1).

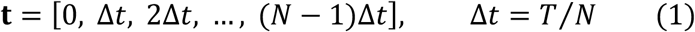

A linear convolution (∗) is performed between the multi-exponential model (*A_i_* are amplitudes, *τ_i_* are lifetimes of each component) and the shifted IRF (**IRF**_*s*_) with truncation (0: *N*) (Eq. 2). The constant offset parameter (*Z*) that represents a time-independent background is also optimized^12^. Together, this yields the fit curve (**ŷ**). FLIM Playground supports fit models with up to three *components*.

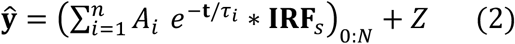

Parameter optimization is performed by minimizing the difference between the fitted curve (**ŷ**) and the measured curve (**y**), quantified by either Maximum Likelihood Estimation (MLE) or Least Square (LS) as a *cost metric*^14^. The fitting procedure also allows users to specify *time gates*, restricting the optimization to a selected temporal window of the decay. The cost metric chosen by the user (MLE or LS) is fed into lmfit^29^, a Python generic curve fitting library, to be minimized (maximizing MLE is mathematically equivalent to minimizing the negative log-likelihood function). FLIM Playground provides three *optimization modes* that balance speed against robustness to local minima. Global optimization employs the differential evolution algorithm^30^, a derivative-free, population-based but relatively slow global optimizer. Local optimization applies the Levenberg–Marquardt^31^ algorithm when the chosen metric is least square, or the Nelder–Mead^32^ algorithm otherwise. Hybrid optimization, the most time-consuming, combines both approaches by first using differential evolution to generate an initial guess, followed by local refinement.

For each high-SNR decay curve, a shift value is optimized and displayed as a point in an interactive scatter plot. By clicking on a point, the user can inspect the measured decay alongside the fitted curve, together with the fit metric and estimated parameters. This interface allows users to evaluate the fits, select a fixed shift to apply across all FOVs, or adjust the time gates to trigger re-optimization. Once the user confirms the *number of components* of the exponential model, *cost metric*, *fitting mode*, *time gates*, and shifts for all channels, these settings are stored in the metadata file and applied to extracting fit features for the entire dataset.

#### 2.3.2 Phasor Calibration

When calibration is performed using the IRF, the temporal shift for each high-SNR decay is estimated by maximizing the cross-correlation between the IRF and the corresponding decay. Similar to fit-based calibration, all estimated shifts are plotted, and the user can select a fixed shift to apply across all decays. In addition to shifting the IRF by the chosen value, each decay curve is corrected by subtracting an offset—defined as the mean of the last tenth percentile of the decay tail—and clipped to zero if negative values occur after subtraction. The raw phasor of the decay is divided by the phasor of the shifted IRF using complex division to obtain the calibrated phasor.

When using a fluorescence lifetime standard for calibration, the offset is subtracted from both the standard fluorophore decay and the data decay curve, and their raw phasors are calculated. Then, phasor calibration is performed using the *lifetime.phasor_calibrate* function from the PhasorPy^17^ library. This function applies the same polar rotation and scaling transform that moves the raw standard fluorophore phasor to its correct phasor location to the raw data decay phasor, yielding its calibrated phasor.

#### 2.3.3 Feature Extraction

For each channel, the selected feature extractors specified in the previous stage are applied to each cell ROI. A single-cell decay curve summed from all the decays of its pixels^19^ can be fit with an exponential model, transformed into phasor coordinates, or both. Additionally, morphological and texture features can be computed from single-cell masks and intensity images.

The *Lifetime Fit* extractor (Supp Man. Chapter 6) applies reconvolution fitting to the decay curve of each cell using the shifted IRF, with fit settings specified by the user in the calibration step. In addition to saving the absolute amplitudes (*A_i_*), it normalizes these amplitudes into fractions (*α_i_*) (Eq. 3) and computes the mean lifetime (*τ*_m_) (Eq. 4).

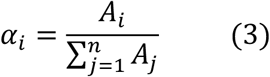

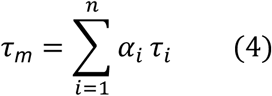

It can also compute cell-level lifetime fit features when pre-fit pixel-level results are available, by averaging the pixel-level features within each cell ROI.

The *Lifetime Phasor* extractor (Supp Man. Chapter 7) provides a fast, model-free view of lifetimes by transforming fluorescence decays into phasor coordinates through a discrete Fourier transform. For each cell, raw phasors are computed from the fluorescence decay for the first and second harmonics of the laser pulse repetition rate, and they are calibrated using the procedures described in *Phasor Calibration* to obtain calibrated phasor coordinates for the first harmonic (*g*, *s*) and second harmonic (*g*_2_, *s*_2_). Phasor-derived lifetime parameters are also computed, including phase (*ϕ*), modulation (*m*), tau phase (*τ_ϕ_*), and tau modulation (*τ_M_*)^15,33,4^.

The *Morphology* extractor (Supp Man. Chapter 8) computes single-cell morphological features of the cell ROI mask, including area, perimeter, solidity, eccentricity, major axis length, minor axis length, and circularity. The list can be readily extended.

The *Texture* extractor (Supp Man. Chapter 9) uses the single-cell ROI intensity image to extract texture features that can be readily extended. For example, *granularity_n_* is the percentage of intensity contributed by bright objects of *n* pixels in radius. It is computed by performing a morphological opening with a disk of radius *n* on the cell intensity image and measuring the intensity lost as a percentage of the total intensity of the original cell ROI. Five granularity-based features are calculated: *n* = 1,3,5,7,9. *Mass displacement* calculates the displacement, as Euclidean distance in pixels, of the intensity-weighted cell ROI centroid from the geometric centroid of the cell ROI. To calculate *radial distribution*, each cell intensity image is partitioned into four concentric rings, and the intensity fraction over the total intensity for each ring is calculated.

Each cell is assigned a unique identifier by concatenating its field-of-view (FOV) identifier with the corresponding label from the ROI mask, and this identifier is used as the row index. Each row then aggregates features extracted from every channel, produced by their respective feature extractors. The extraction status for each FOV is tracked with a progress bar, and the results are displayed for preview before being exported.

### 2.4 Extract Single-Cell Categorical Features

In this stage, FLIM Playground merges cell-level datasets produced by the numerical feature extraction stage into a unified table and assigns categorical features to each cell (Figure 2C, Supp Man. Chapter 10). FLIM Playground searches the user-designated folder for CSV files and verifies the uniqueness of cell identifiers. Datasets are then merged by aligning shared columns and concatenating rows. FOV identifiers are assumed to consist of segments delimited in a consistent manner, where each segment or combination of segments encodes a categorical feature (e.g., treatment, time point). Under this assumption, FLIM Playground parses the segments and presents them to the user, who interactively assigns them to categorical features with a live preview of the results. The resulting merged dataset, enriched with categorical features, can be exported for downstream analysis.

### 2.5 Data Analysis provides a shared interactive interface for diverse analysis methods

After single-cell datasets are generated—either through Data Extraction or by the user’s own methods—the Data Analysis section provides a suite of analysis methods (Table 2) to help the user gain insights. To ensure users can seamlessly apply alternative analysis methods across datasets with varying features, Data Analysis offers a unified interface built on a library of interactive widgets shared across its analysis modules, allowing new modules to be added easily by reusing existing components. Supplemental Video 2 shows a user iteratively exploring a dataset by switching among modules within the shared interface.

**Table 2.** Modules offered in FLIM Playground Data Analysis. Modules are grouped into three analysis classes—Univariate, Bivariate, and Multivariate—based on the number of numerical features required, with custom feature-selection widgets implemented for each class (yellow). Each module pairs an interactive visualization with a statistical modeling method. *Filter by* and *Color by* are interactive widgets derived from experiment-level categories (blue) and are shared by all modules (*The Classification module renames *Color by* to *Classify by* to form classification groups). For modules that come with point-based (red, blue) visualizations (e.g., scatter plot), hover-based interactions that show cell identifiers (red) and visual channel widgets derived from categorical features (blue) are implemented. Each module also comes with built-in modeling methods, supported by custom interactive widgets.

| Analysis class &<br>Numerical feature<br>selection | Modules | Filter<br>by | Color<br>by | Point-based | Built-in modeling methods |
| --- | --- | --- | --- | --- | --- |
| **Univariate**—Select *one* feature from a feature group. | Feature Comparison | ✓ | ✓ | ✓ | Effect size calculation & Statistical tests |
|  | Feature Histogram | ✓ | ✓ | ✗ | Gaussian Mixture Model & Heterogeneity-index |
| **Bivariate**—Two univariate selectors; or choose a channel and a harmonic to select phasor coordinates. | Feature Distribution | ✓ | ✓ | ✓ | 2D Gaussian Mixture Model & Linear Regression |
|  | Phasor Analysis | ✓ | ✓ | ✓ | K-means Clustering |
| **Multivariate**—Select multiple features from each feature group. | Dimension Reduction | ✓ | ✓ | ✓ | PCA, t-SNE, UMAP with user-adjustable hyperparameters |
|  | Classification | ✓ | ✓* | ✗ | Random Forest, SVM, Logistic Regression, Gradient Boosting |

#### 2.5.1 The Shared Interface

The shared interface supports common analysis tasks and is composed of interactive widgets built to operate the three feature classes—identifiers, numerical features, and categorical features (Supp Man. Chapter 11). The user has the option to map features in external datasets, i.e., datasets not output by Data Extraction, to these classes in the analysis configuration (Supplemental Video 3, Supp Man. Chapter 12). This option allows users to perform data extraction outside of FLIM Playground but still use the analysis methods provided by FLIM Playground.

Numerical features are grouped either by channel–extractor pairs or by custom groupings specified in the analysis configuration, making it faster to locate and select features. Univariate methods present one single-select widget per group; choosing a feature in any group clears the selections in the others. Bivariate methods present two such panels, with the first chosen feature excluded from the second panel. Multivariate methods present one multi-select widget per group, with an *All* option to quickly include every feature in that group.

Filter widgets, each linked to a categorical feature, allow users to select one or more labels to find data of interest, with an *All* option available to include every label in that feature.

Visual channel widgets map categorical features to colors, shapes, and opacities in the visualizations. Users can select multiple categorical features in a *Color by* widget; each unique label combination partitions the filtered dataset, with a distinct color assigned to each partition. For example, if the filtered dataset contains three treatments and two cell lines, and both features are selected in the *Color by* widget, six non-overlapping treatment–cell line partitions will be created. If no feature is selected, all data are grouped into a single partition and displayed in the same color. In point-based visualizations, *Opacity by* and *Shape by* widgets allow a single categorical feature to be mapped so that each label is represented with a distinct opacity or shape. Interactive visualizations are implemented in Plotly^34^, enabling users to hover over individual points (cells) to view their cell and field-of-view identifiers.

Widgets are also provided to allow interactive adjustment of key visualization parameters, including point size in point-based visualizations, axis label size, legend size, and the colormap applied to distinguish groups.

#### 2.5.2 Module-Specific Statistical Methods

Custom widgets are implemented to support module-specific modeling, and adjustments to them trigger real-time reanalysis. All models are applied separately to each partition created by the Cartesian product of the categorical features selected in *Color by*. These partitions will hereafter be referred to as color groups.

In *Feature Comparison* (Supp Man. Chapter 13), users can choose to compute effect sizes and/or perform statistical tests for each comparison pair and annotate the results, quantifying the difference and testing its significance. FLIM Playground offers two effect size calculation methods: Glass’ Delta^35^ and Cohen’s d^36^. The two supported statistical test options are implemented using the Scipy^37^ function *scipy.stats.ttest_ind*: Student’s t-test (the two groups are assumed to have equal variances) and Welch’s t-test^38^ (it generalizes the former test to account for unequal variances by specifying *equal_var=False*).

Available comparison pairs include all pairwise combinations of color groups. Because the number of pairs can grow combinatorially and many may be uninformative, users can restrict the annotations displayed in the visualization either by deselecting them in a selection widget or by setting a numerical threshold to show only annotations above a chosen value. The accompanying visualization comprises Sina plots for each color group, revealing single-cell heterogeneity and group-level distributions of the selected numerical feature, with filtered effect size annotations overlaid. Users can designate a categorical feature in the *Separate by* widget to generate sub-panels corresponding to the labels of the selected feature. The *Color by* widget then creates color groups and comparisons are performed between pairs in each panel.

In *Feature Histogram* (Supp Man. Chapter 14), the selected numerical feature is visualized on the x-axis, and the y-axis represents either the counts or densities estimated by the Gaussian Mixture Model^39^ (GMM) of each color group. The counts are determined by the user-adjustable bin width of the histograms. The default bin width is determined automatically by NumPy^40^ as the minimum of the Sturges’^41^ and Freedman–Diaconis^42^ estimators.

As an alternative to histograms, Gaussian Mixture Models (GMMs) can capture multimodal distributions by representing a dataset as a weighted sum of Gaussian components, each with its component weights, means, and variances. FLIM Playground uses scikit-learn^43^ to fit all GMMs up to the user-controlled maximum number of components, retain only those in which all components exceed the user-specified minimum weight, and select the model with the lowest Bayesian Information Criterion^44^ to prevent overfitting. To quantify the subpopulation structure of each color group from the GMM fit, a weighted entropy-based heterogeneity index can be computed and displayed^45,46^.

Each point can be assigned to a GMM component, which can then be exported and used as a categorical feature. In *Hard Assignment*, the posterior probability (responsibility) of each component is computed, and the data points are assigned to the component with the highest responsibility. Alternatively, *Intersection Thresholding* finds intersection points of adjacent component distributions using Brent’s method^47^ and assigns each point to the component whose region (bounded by these intersections) it falls into.

The bivariate *Feature Distribution* module (Supp Man. Chapter 16) visualizes the first selected feature on the x-axis and the second on the y-axis as a scatter plot. A two-dimensional GMM can be fit to each color group, following the same procedure as in *Feature Histogram*. The correlation coefficient and its p-value are calculated using SciPy, and a linear regression line can be fit using scikit-learn and displayed.

The *Phasor Analysis* module (Supp Man. Chapter 17) plots the phasor coordinates into the universal semicircle^33^. K-means clustering^48^ is performed with a user-specified number of clusters, after standardizing the phasor coordinates since *g* (real part) and *s* (imaginary) may differ in scale. Then scikit-learn is used to find the optimized clustering. A convex hull, a polygon that contains all the points of each cluster, is drawn for each cluster, and the cluster centroid is marked with its coordinates displayed.

In *Dimension Reduction* (Supp Man. Chapter 18), selected numerical features are standardized and projected into two dimensions for visualization. Three popular algorithms are provided: Principal Component Analysis (PCA^49^), t-distributed Stochastic Neighbor Embedding (t-SNE^50^), and Uniform Manifold Approximation and Projection (UMAP^51^). Since hyperparameters affect projection results, users can adjust perplexity and early exaggeration when using t-SNE, while other parameters follow scikit-learn defaults. For UMAP, the number of neighbors and minimum distance are adjustable, while other parameters are left at umap-learn defaults.

In the *Classification* module (Supp Man. Chapter 19), machine learning classifiers including Random Forest, Support Vector Machine, Logistic Regression, and Gradient Boosting are provided. Their performance is evaluated using accuracy, precision, recall, F1 score, and visual diagnostics including confusion matrices and Receiver Operating Characteristic (ROC) curves with corresponding Area Under the Curve (AUC). All classifiers and performance metrics are implemented with scikit-learn, using a fixed random seed to ensure reproducibility. For decision tree–based models (Random Forest and Gradient Boosting), feature importance plots are also generated to rank the contribution of each feature.

A set of model-specific widgets are implemented to facilitate the classification process. Users can select the classifier and specify the proportion of data allocated to training and testing. To address class imbalance, three sampling strategies are provided. By default, stratified sampling that preserves the original class distribution is applied. Alternatively, undersampling^52^ reduces all classes to the size of the smallest class, and oversampling increases all classes to the size of the largest class by randomly duplicating samples from minority classes. The visual channel widget *Color by* is renamed to *Classify by* and is used to form classes based on selected categorical features. User-selected options for both binary and multi-class classification, augmented by one class versus the rest, are available.

### 2.6 Two-photon fluorescence lifetime imaging microscopy of cells

PANC-1 human pancreatic (ATCC, RRID: CVCL_0480) and MCF7 human mammary gland adenocarcinoma (ATCC, RRID: CVCL_0031) cells were cultured in high-glucose Dulbecco’s modified Eagle’s medium supplemented with 10% fetal bovine serum (Gibco) and 1% penicillin/streptomycin (Gibco). Cell lines were regularly tested for mycoplasma contamination. 2×10^5^ cells were plated on 35mm glass-bottom dishes (MatTek) 48 hours prior to imaging and maintained at 37°C and 5% CO_2_. The following metabolic inhibitors were added to the dish as indicated prior to imaging—sodium cyanide (NaCN; inhibits oxidative phosphorylation) (Fisher Scientific): 4mM for 15 minutes; 2-Deoxy-d-glucose (2-DG; inhibits glycolysis) (Sigma Aldrich): 10mM for 2 hours; iodoacetic acid (IAA; inhibits glycolysis) (Sigma Aldrich): 1.5mM for 30 mins. FLIM was performed on a custom-made Ultima Multiphoton Imaging System (Bruker) that consists of an inverted microscope (TI-E, Nikon). The system is coupled to an ultrafast tunable laser source (Insight DS+, Spectra Physics Inc). The fluorescence lifetime images were acquired using time-correlated single-photon counting (TCSPC) electronics (SPC-150, Becker & Hickl GmbH) and imaging was performed using Prairie View Software (Bruker). NAD(P)H and FAD were sequentially excited using excitation wavelengths of 750 nm and 890 nm respectively with the laser power at the sample <10 mW. The samples were illuminated using a 40x objective lens (W.I./1.15 NA /Nikon PlanApo) with a pixel dwell time of 4.8 µs and frame integration of 60s at 256×256 pixels. The photon count rates were maintained at 1–5 × 10^5^ photons/second to ensure adequate photon detection for accurate lifetime decay fits. A 720nm dichroic mirror and bandpass filters separated fluorescence signals from the excitation laser. Emission filters used were bandpass 460/80 nm for NAD(P)H and 500/100 nm for FAD. Fluorescence signals were collected on GaAsP photomultiplier tubes (H7422P-40, Hamamatsu, Japan). The IRF was collected each day by recording the second harmonic generation signal of urea crystals (Sigma-Aldrich) excited at 890nm. For each condition, data were acquired from two replicates, and three to four images were acquired per replicate per condition.

### 2.7 Single-Cell Segmentation

Single-cell segmentation of whole-cell masks was performed using either a customized CellProfiler pipeline or the Cellpose algorithm applied to NAD(P)H fluorescence intensity images. The CellProfiler pipeline, previously validated for both 2D cell and 3D organoid images, began with the manual identification of nuclei as primary objects, which served as seed regions for segmentation. Subsequently, whole-cell boundaries were delineated automatically using a Voronoi-based segmentation approach^53,54^. Alternatively, automated whole-cell masks were generated using Cellpose^7^ (model = cyto3, radius = 30). The resulting masks were then visually inspected and manually refined in Napari^8^ to ensure accurate delineation of individual cells.

### 2.8 SPCImage Analysis

To enable fitting using the SPCImage software (Becker & Hickl GmbH) at the cell level, preprocessing was required: for each cell, the decays of all pixels within the same cell were summed. The resulting cell-level decay was then propagated back to replace the decays of all pixels belonging to that cell, so that the pixel-fitting routine by SPCImage could be applied to fit cell-level decays^19,27^. Because all pixels of a given cell now shared the same decay curve, the binning factor was set to zero. The IRF time-shift was selected manually from one FOV by clicking on representative pixels containing cell decays; upon selection, SPCImage fits the decay and outputs the corresponding shift value. The median shift across those pixels was chosen and applied to all FOVs.

## 3 RESULTS

### 3.1 Fit Validation

To validate the *Lifetime Fit* extractor in FLIM Playground, 25 NAD(P)H FLIM images of PANC-1 cancer cells^19^ were fit using SPCImage and FLIM Playground. SPCImage yielded 6.40 time bins (or 307 ps) as the IRF shift, while FLIM Playground produced 6.53 time bins (or 313 ps). The cost metric (MLE), number of exponential components (2), and time gates were matched between the two methods to ensure a fair comparison. The average photon count across all cells was 4626. Lifetimes and fractions of each component were obtained from fits performed with these two methods.

The comparisons of two representative lifetime features (*τ_m_*, *α*_1_) are shown in Figure 3. Although the mean of *τ_m_* and *α*_1_ calculated by FLIM Playground deviated from the SPCImage fit (Fig 3B, E) on average by -15.4 picoseconds (ps) [95% CI: -16.7 ps, -14.2 ps] and 0.18% [95% CI: 0.1%, 0.2%] respectively, the effect sizes were small (Figure 3A, D), and the FLIM Playground values correlated consistently with the SPCImage values (*r* = 0.99 and *p* < 0.001). A regression line was fit for each of the two features, whose 95% confidence interval for the slope included 1, indicating no evidence of proportional bias (Figure 3C, F).

**Figure 3.**
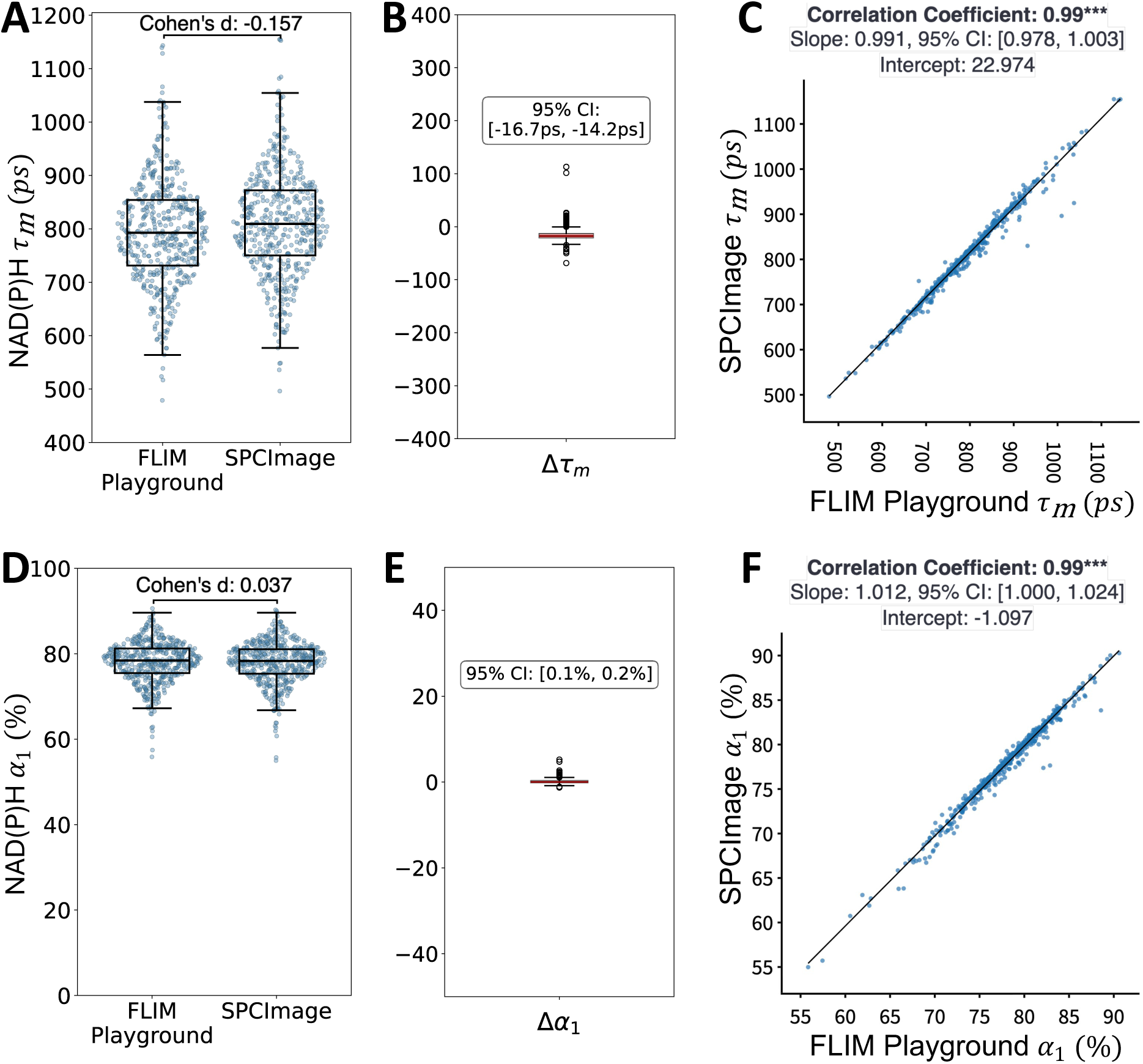
Fit Comparison between FLIM Playground and commercial software (SPCImage). The first row shows the comparison of the NAD(P)H mean lifetime (*τ_m_*) calculated from FLIM Playground and SPCImage. **(A)** Distributions of *τ_m_*are shown for both methods, with Cohen’s d calculated between their means. **(B)** Δ*τ_m_* distribution with the 95% confidence interval annotated. **(C)** A scatter plot annotated with the correlation coefficient between the *τ_m_* values obtained from the two fit methods (*p* < 0.001), exported from the *Feature Distribution* module. A regression line (in black) was fit and plotted. Its slope, confidence interval, and intercept are reported. **(D, E, F)** show the same visualizations and statistics for comparing the proportion of free NAD(P)H (*α*_1_).

### 3.2 Discriminating Inhibitor Treatments in PANC-1 and MCF7 Cells Using FLIM Playground

To demonstrate the integrated workflow of FLIM Playground, the following FLIM dataset was acquired: three inhibitors—cyanide, 2-DG, and IAA—were added to each of the two cell lines (PANC-1 and MCF7), alongside a control condition, and NAD(P)H and FAD channel FLIM images were acquired along with IRFs. Whole-cell masks were made using CellProfiler^21^ based on the NAD(P)H intensity images. Analysis was performed in FLIM Playground, including extracting single-cell features and visualizing and applying models to those features.

The settings were specified using the configuration interface (Supp Figure 1) and were kept consistent across subsequent stages. The FOV identification (Supp Figure 2) and IRF calibration (Supp Figure 3) stages were repeated for each day. Fits were performed in Local mode, and a bi-exponential decay model was applied to both channels with the same cost metric (MLE) and the same time gates ([49, 240]). The fit lifetime features, phasor features, morphology, and texture features were extracted for both channels (Supp Figure 4). All extracted features are shown in Supp Table 1.

The Data Extraction section was concluded by the categorical feature extraction stage, which merges the four single-cell numerical datasets and categories (cell line, treatment, and dish number) were interactively assigned to cells (Supp Figure 5).

The combined dataset was uploaded to the Data Analysis section, and a subset of modules was applied to gain biological insights (Figure 4, 5). In the *Feature Comparison* module, through the shared interactive interface, NAD(P)H *τ_m_*was selected, categories of interest were selected (Figure 4A), and categorical features were assigned to visual channels (Figure 4B). Differences between groups were quantified using effect size calculations, where Glass’ Delta was calculated for selected comparison pairs (Figure 4C), and an effect size threshold was chosen as 0.7 (Figure 4D). This produced a visualization showing the distributions of NAD(P)H *τ_m_* and *τ_ϕ_* for different inhibitors across two cell lines, with replicates encoded in different opacities. Based on the large overlap, technical replicate variability was relatively small. As expected, IAA increased the NAD(P)H *τ_m_* and *τ_ϕ_* compared to the control, whereas inhibition of oxidative phosphorylation by cyanide decreased NAD(P)H *τ_m_* and *τ_ϕ_* for both cell lines (Figure 4E, F).

**Figure 4.**
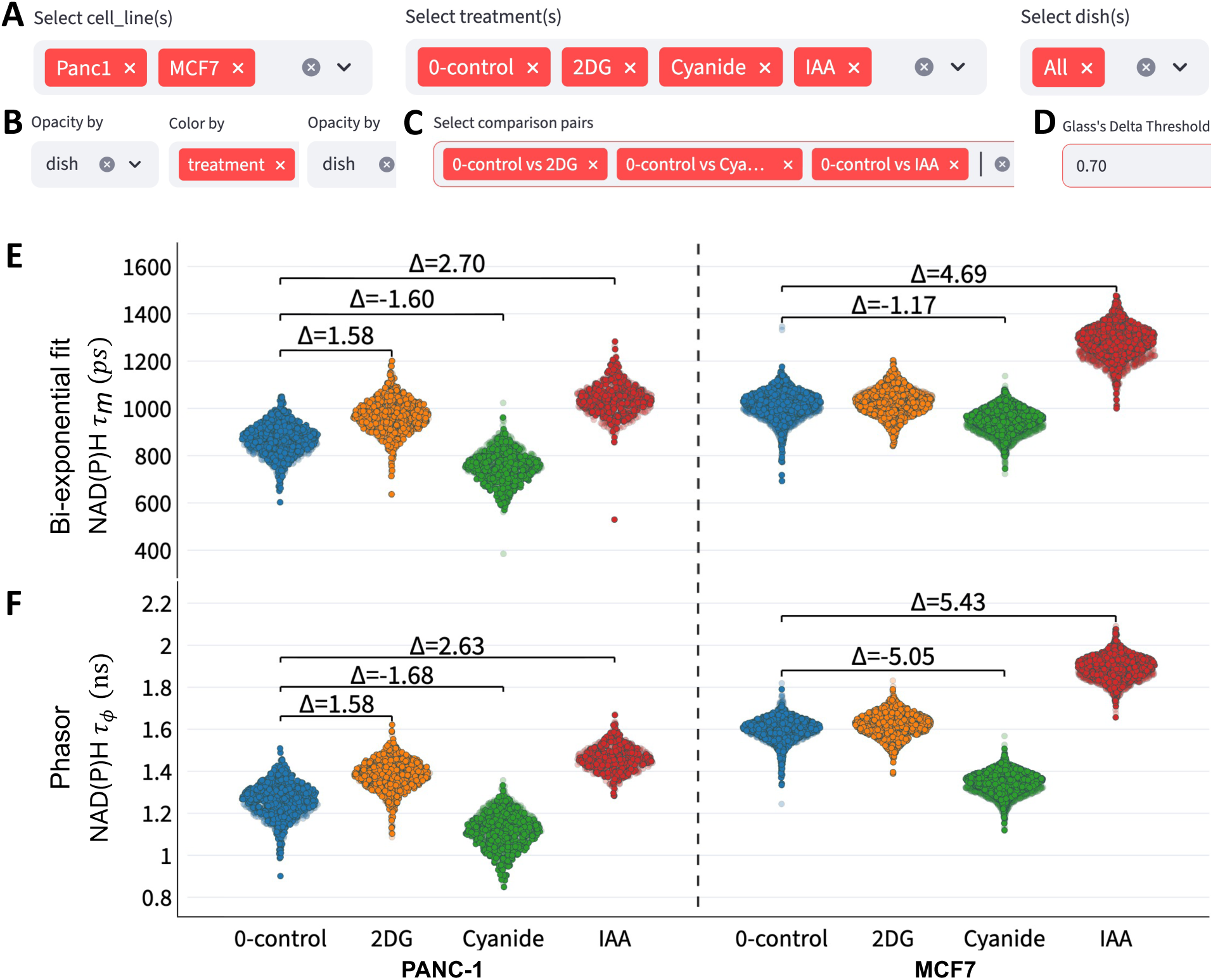
Applying the Feature Comparison module to compare features across categories. **(A)** Filter widgets are rendered for each categorical feature. Two cell lines, three inhibitors plus the control, and both dishes were selected. **(B)** Data separated by cell line, each generating one sub-panel; within each panel, treatments are color-coded. Different opacities indicate variability between dishes. **(C)** Three comparison pairs, control versus each inhibitor, were selected, and **(D)** only effect sizes (Glass’ Delta) exceeding the user-specified threshold (0.7) were shown. **(E)** NAD(P)H *τ_m_* was compared between the inhibitors and the control, separated by cell line (left: PANC-1, right: MCF7). **(F)** Same comparisons plotted for NAD(P)H tau phase (*τ_ϕ_*), a phasor derived lifetime feature.

**Figure 5.**
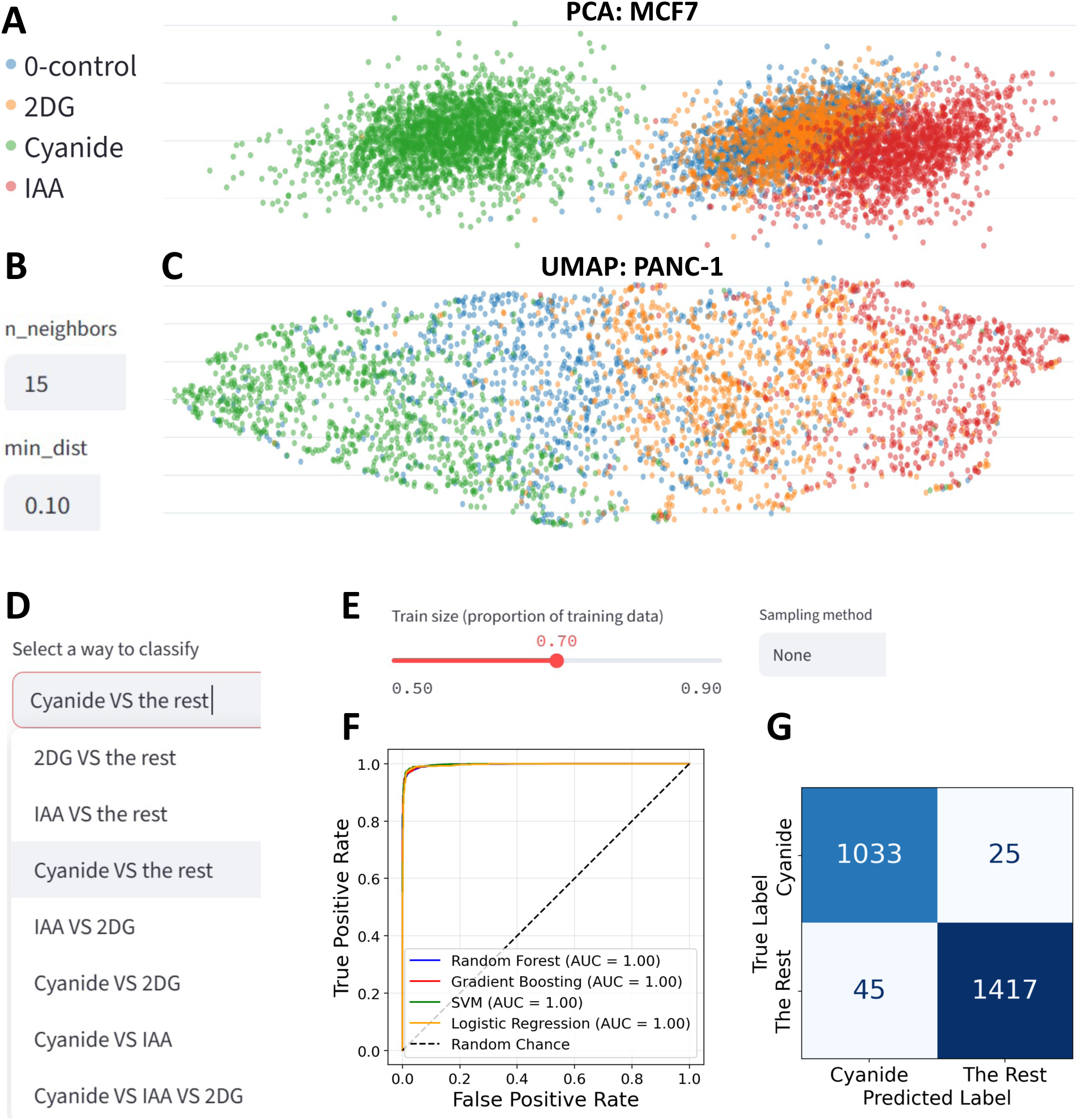
Applying the Dimension Reduction module and the Classification module to differentiate inhibitor treatments. **(A)** PCA was performed on MCF7 cells (axes were truncated). **(B)** Interactive widgets for specifying UMAP hyperparameters (*n_neighbors* = 15, *min_dist* = 0.1); **(C)** UMAP of PANC-1 cells. **(D)** List of all classification options; cyanide versus the rest (two glycolysis inhibitors: 2-DG and IAA) was selected. **(E)** Train–test split: 70% (5,877 cells) training and 30% (2,520 cells) testing and the sampling method used (None: default stratified sampling). **(F)** ROC curves generated from FLIM Playground classifiers. All classifiers were trained on the same numerical feature set used for UMAP. **(G)** Confusion matrix of the Random Forest classifier evaluated on unseen data, showing number of cells within True Positive (upper left), False Negative (upper right), False Positive (lower left), and True Negative (lower right) categories.

The PCA from the *Dimension Reduction* module was demonstrated on MCF7 cells, using all features (Supp Table 1) except single-cell intensity sum to ensure independence from acquisition-related intensity differences (Figure 5A). The separation between control and 2-DG was less distinct, consistent with the small effect size observed between these groups for both *τ_m_*and *τ_ϕ_*. UMAP provided a clearer separation between treatments in PANC-1 cells using the same feature set (Fig 5B, C): the two glycolysis inhibitors (2-DG, IAA) clustered on one side while the oxidative phosphorylation inhibitor (cyanide) localized to the opposite region of the embedding space, and the control lay in the middle.

Whereas dimension reduction methods provided a qualitative view of treatment separation, the *Classification* module was used to quantify the discriminative power of the features. To discriminate between cyanide, the oxidative phosphorylation inhibitor, and glycolysis inhibitors (2-DG and IAA), “cyanide versus the rest” was selected in the classification options list (Figure 5D). The train-test split was 70% and 30% (Figure 5E). Since the classes (cyanide versus 2-DG + IAA) were not substantially imbalanced, the default stratified sampling was used. All classifiers achieved an AUC of 1.0 on the ROC curves (Figure 5F), with the near diagonal confusion matrix (of the Random Forest classifier, Figure 5G), indicating strong discriminative ability between cyanide versus the glycolysis inhibitors (2-DG and IAA).

### 3.3 Analysis of Fluorescence Lifetime Flow Cytometry Data Using FLIM Playground

To validate FLIM Playground, we reanalyzed a previously published dataset of primary human T cells acquired by fluorescence lifetime flow cytometry^18^. This is an example of a dataset without imaging, but with single-cell decays. The data comprise two-dimensional decays in a tabular format: rows are individual cells and columns are successive time bins of the decay. Consistent with the reported results, activated T cells exhibited shorter mean NAD(P)H lifetimes (*τ_m_*) and a higher fractional contribution of the short component (*α*_1_) compared to quiescent cells. The phasor distribution reproduced the clear separation between activation states, and UMAP embeddings of lifetime fit and phasor features likewise resolved two distinct clusters. Random forest classification based on FLIM Playground-extracted features achieved accuracy and AUC values comparable to those previously reported, confirming that the framework faithfully reproduces published lifetime fit, phasor, and analyses results (Supp Figure 6).

## 4. DISCUSSION

FLIM Playground is an open-source, interactive platform that unifies metadata organization, calibration, single-cell feature extraction, visualization, and statistical modeling through an intuitive and code-free graphical interface. In current practice, FLIM analysis involves switching between a fragmented set of tools and processing their outputs to obtain biologically interpretable features and insights, often requiring custom code and expertise. FLIM Playground overcomes this limitation by mapping the hierarchical levels of FLIM data to three semantically meaningful feature classes—identifiers, numerical features, and categorical features (Table 1).

The Data Extraction section adopts a channel-centric framework, in which each channel can be assigned its own extraction methods. Notably, extraction is not limited to lifetime features: morphological and texture descriptors are also supported, enabling a richer characterization of cellular phenotypes than is currently available in existing tools and extending feature extraction to non-FLIM imaging modalities. In addition to single-cell numerical features, identifiers and categorical features are also incorporated into a seamless three-stage extraction interface. Beyond single cells, this framework accommodates any segmented ROI—for instance, organoids—whose boundaries are defined by ROI masks and labeled with unique ROI IDs.

The Data Analysis section then transforms these three feature classes of single-cell datasets—whether produced by Data Extraction or imported as user-provided data (including from other fields)—into a unified interface shared across visualization and analysis methods built on the Python ecosystem. This interface allows users to explore datasets from multiple perspectives—inspecting individual cells, filtering data of interest, and visualizing features through visual encodings that leverage the human visual system’s capacity for rapid, parallel detection of patterns and trends, revealing insights that raw numbers or text may obscure^55^. The interface further couples with module-specific analysis methods and their corresponding interactive components, enabling specialized statistical modeling to reveal biological insights.

To ensure reliability, we benchmarked lifetime fits in FLIM Playground against the widely used commercial package SPCImage and demonstrated that the two yielded consistently correlated lifetimes with small differences. Additionally, to demonstrate the integration and its capabilities, we used FLIM Playground to run the complete FLIM analysis pipeline—from feature extraction through analysis—to representative two-photon NAD(P)H and FAD FLIM images of cancer cell lines (PANC-1 and MCF7) treated with metabolic inhibitors. FLIM Playground was also demonstrated on a fluorescence lifetime flow cytometry dataset of primary human T cells comprised of 2D decays (*i.e.*, non-imaging measurements). FLIM Playground successfully discriminated between different inhibitors and activation states, respectively.

While integrating downstream analysis with data extraction enables users to explore biologically meaningful variation across cells in the same platform, several limitations remain. At present, the software is designed for TCSPC-acquired data; however, support for other time-domain and frequency-domain acquisition modalities could be implemented in future versions. Similarly, additional fit-free methods, such as Laguerre deconvolution^4^, may be incorporated to broaden the feature extraction repertoire. Fitting is currently limited to the cell level due to computational speed constraints, making pixel-level fitting impractical, though FLIM Playground can accept pixel-level prefit lifetime features or allow users to perform cell-level fitting which improves accuracy and precision for cell-level analysis^19^. Although pixel-level phasor calculation, visualization, and texture descriptors of phasor feature maps are technically feasible, they have not yet been implemented. The extraction stage can also be expanded to include a wider range of morphological and texture feature descriptors. Finally, the current version does not support time-lapse or volumetric FLIM or object tracking. Numerical features that capture relationships across time, such as velocity, and their correlation with lifetime features would provide valuable insights and will be considered in future developments. Planned extensions to Data Analysis include hierarchical clustering for exploratory grouping, linear mixed-effects models to account for biological and technical variability, and modality alignment methods that integrate FLIM-derived numerical features with data from other omics or imaging modalities.

FLIM Playground makes advanced FLIM analysis accessible to non-computational practitioners through its intuitive interface, while its modular design facilitates improvements by specialists, enabling the framework to be readily adapted for non-FLIM acquisitions as well. The platform can be expanded to incorporate additional acquisition modalities, feature descriptors, fit-free methods, and analysis modules, supporting a range of applications in quantitative imaging. By unifying extraction, visualization, and analysis in a single platform, FLIM Playground supports biologically meaningful discoveries at the single-cell level and at scale.

## Acknowledgements

The authors would like to thank all members of the Skala Lab for their suggestions, as well as those who kindly proofread the online manual. We are also grateful to Matthew Stefely for designing and creating the FLIM Playground logo and editing figures and Alicia Williams for editing the manuscript.

## Funding

This work was funded by

NIH R01 CA278051,

R01 CA272855,

R01 HL165726,

R37 CA226526,

NSF 2426316

## Author Contributions

Conceptualization: WZ, KS, MCS, RD*,

Methodology: WZ, KS

Investigation: WZ, RD*

Formal analysis: WZ, KS

Funding acquisition: MCS

Writing – Original Draft Preparation: WZ, RD*

Writing – Review & Editing: WZ, KS, MCS, RD*

*corresponding author

## Code and Data Availability

The source code, dataset used, and the software executables are available at https://github.com/skalalab/flim_playground.

