## Supplemental material for "FLIM Playground: An interactive, end-to-end graphical user interface for analyzing single cells with fluorescence lifetime imaging microscopy"

### *Supplemental Figures and Table*

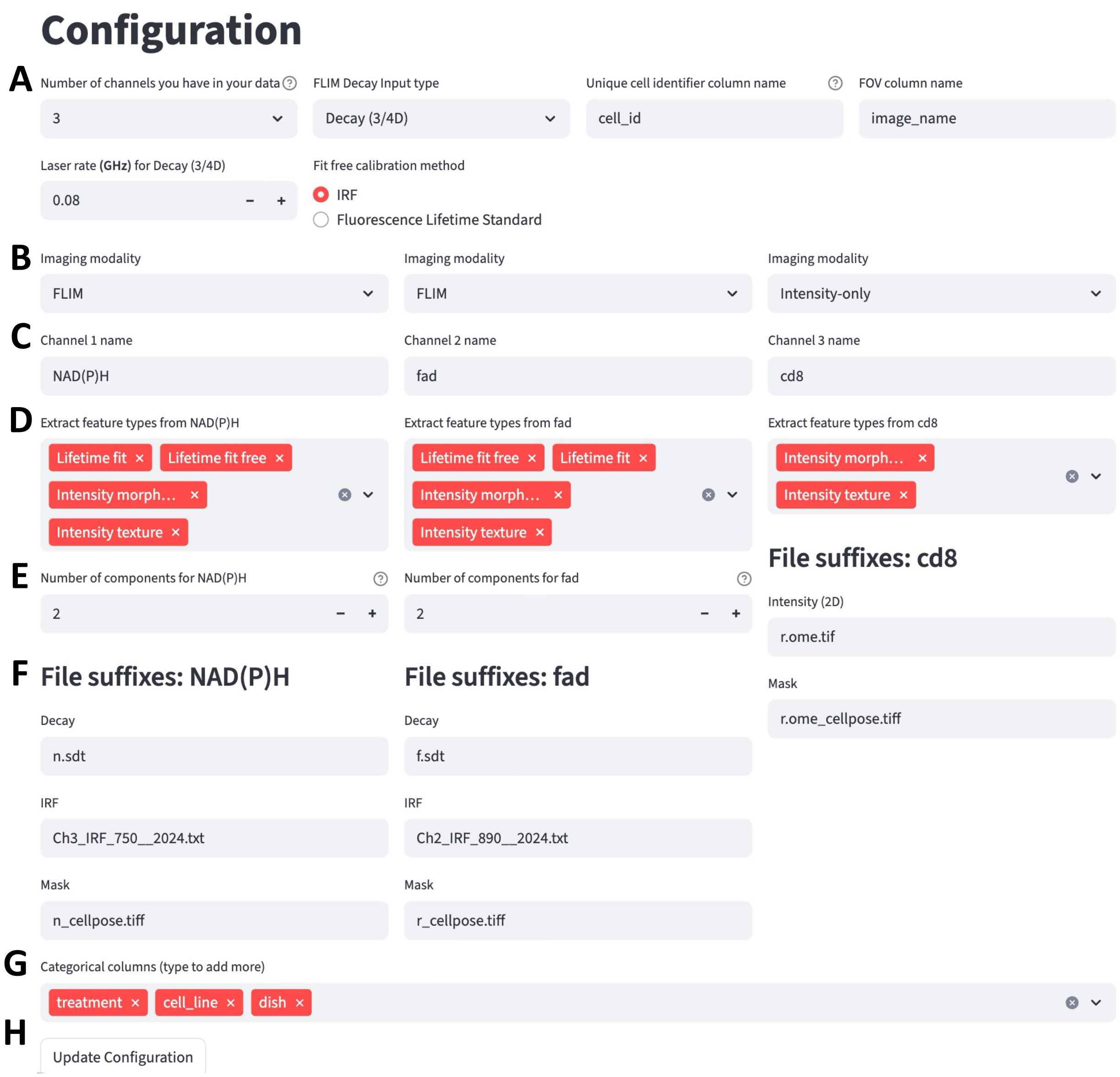

**Supp Figure 1. An example of the configuration interface.** FLIM Playground loads the previously saved configurations and displays them for users to update. **(A)** The first two rows show configurations applied to all acquisition channels: number of channels, decay type, names of the columns that store the unique cell identifiers and field-of-view identifiers, laser frequency, and calibration method for extracting phasor features. Based on the specified number of channels, one column is rendered for each channel that stores channel-level configurations: **(B)** imaging modality, **(C)** channel name, **(D)** feature extractors, **(E)** number of exponential components if Lifetime Fit extractor is selected, and **(F)** file suffixes for each required input file type. Different channels can apply different feature extractors and focus on different cell ROIs. **(G)** Categorical features to be extracted are specified here. **(H)** The configuration can be saved for future use by clicking the update button.

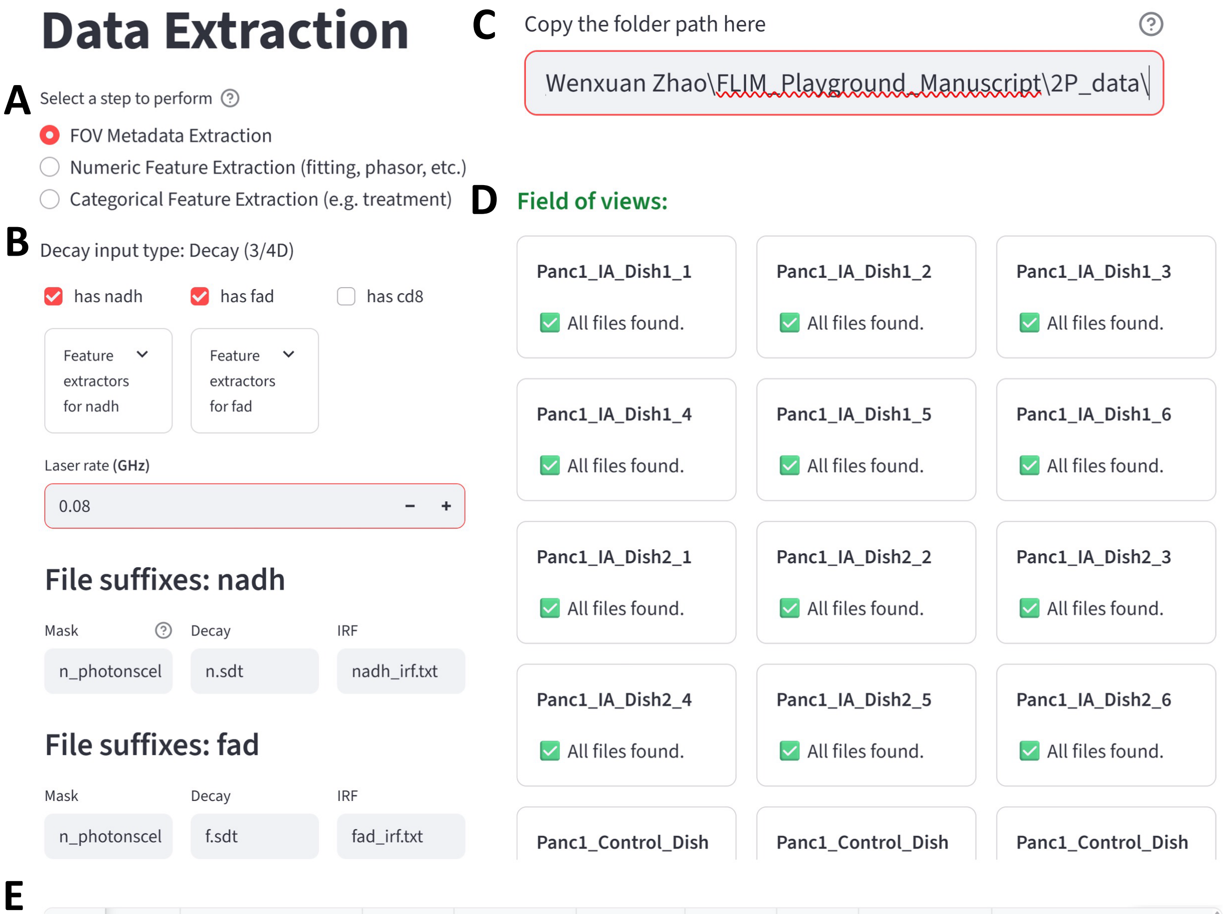

| | y texture | fad_num_components | laser_rate | nadh_channel | fad_channel | time_bins | duration | fov_dimensions | fit_free_calibration_method |
|---|---|---|---|---|---|---|---|---|---|
| 0 | | 2 | 0.08 | 1 | 0 | 256 | 10.0067 | (256, 256) | IRF |
| 1 | | 2 | 0.08 | 1 | 0 | 256 | 10.0067 | (256, 256) | IRF |
| 2 | | 2 | 0.08 | 1 | 0 | 256 | 10.0067 | (256, 256) | IRF |

**Supp Figure 2. Identify FOVs of inhibitor-treated PANC-1 cells. (A)** This was the first of the three stages of the Data Extraction section. **(B)** The parameters were retrieved from a saved file updated by the configuration interface (Supp Figure 1), and the third acquisition channel was deselected. **(C)** After the user specified the folder path containing all the required files, **(D)** a widget card was rendered for each FOV showing the file lookup status (some FOVs were cut off due to space limitation here). **(E)** In addition to the file paths, file metadata and other configurations were checked and saved.

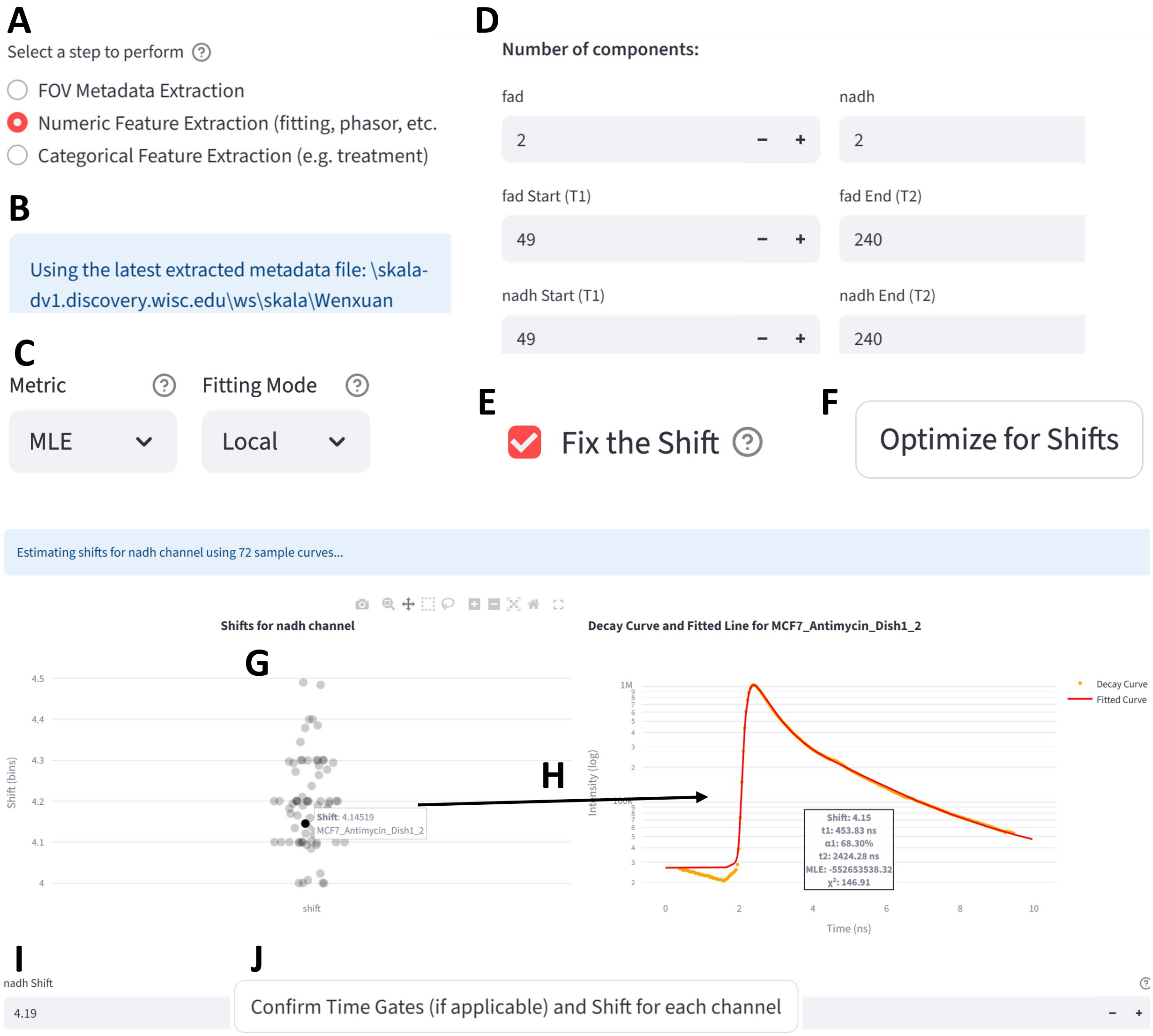

**Supp Figure 3. IRF calibration on FOVs of inhibitor-treated PANC-1 and MCF7 cells.** A calibration interface was displayed for the first step of the **(A)** numerical feature extraction stage. **(B)** It loaded the latest extracted data automatically from the cache. **(C)** The chosen cost metric and the fitting mode, **(D)** number of exponential components, and the time gates of the reconvolution fitting. **(E)** *Fix the Shift* was chosen, so the selected shifts were applied to all FOVs. Once all configurations were confirmed, the user started fitting for shifts on high-SNR decays by clicking the **(F)** *Optimize for shifts* button. **(G)** The distribution of optimized shifts was plotted, allowing the user to hover and click on individual points to view **(H)** the fitted curve alongside the decay curve with diagnostic plots. Users could hover over the decay curve to display the corresponding time bin number, which helped in selecting appropriate time gates. **(I)** The shift selector was automatically populated with the median shift of the distribution, which was selected in this case. Steps **(G–I)** were repeated for the FAD channel as well. The shifts, time gates, and other fitting settings were confirmed by clicking the **(J)** *confirm* button.

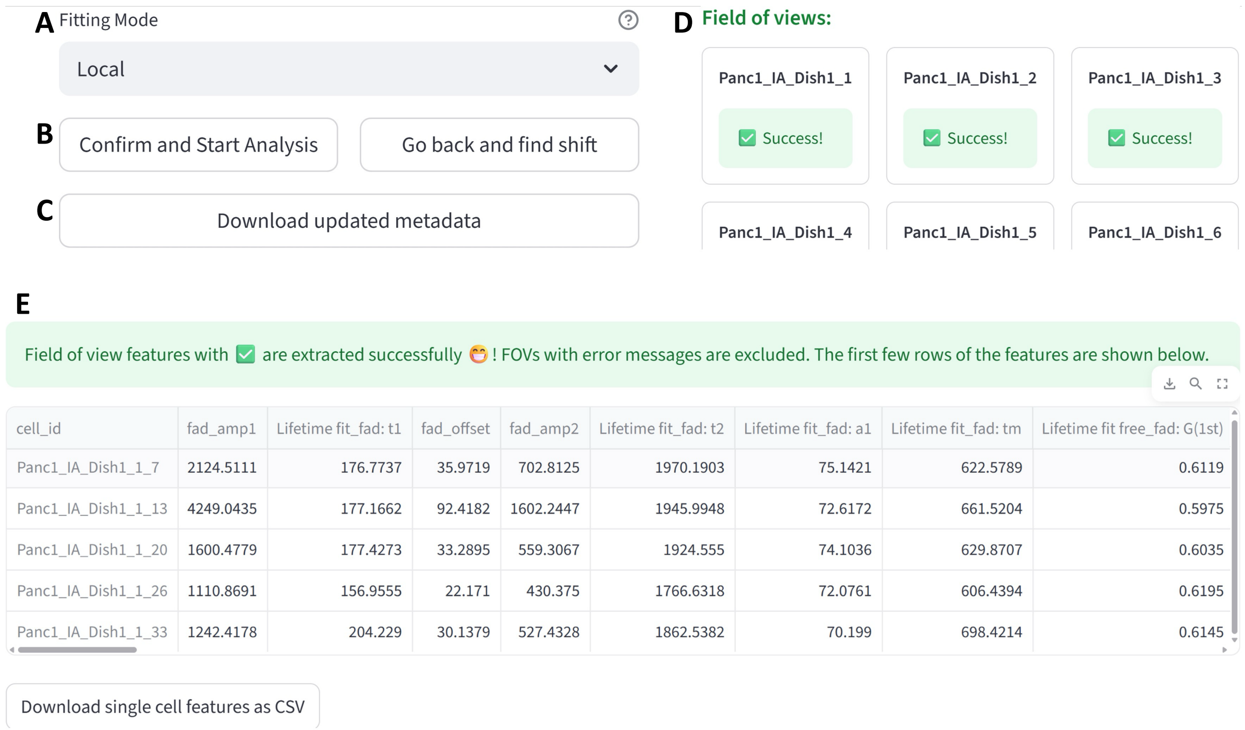

**Supp Figure 4. Numerical feature extraction for FOVs of inhibitor-treated PANC-1 cells. (A)** Users can choose a different fitting mode after IRF calibration. **(B)** They can either confirm the shifts and fitting configurations or choose to go back and recalibrate. **(C)** Update the FOV metadata file with the configuration values. **(D)** After starting the analysis, the selected feature extractors of each acquisition channel were applied to each FOV. The extraction status was displayed, **(E)** and the preview of the extracted dataset was displayed.

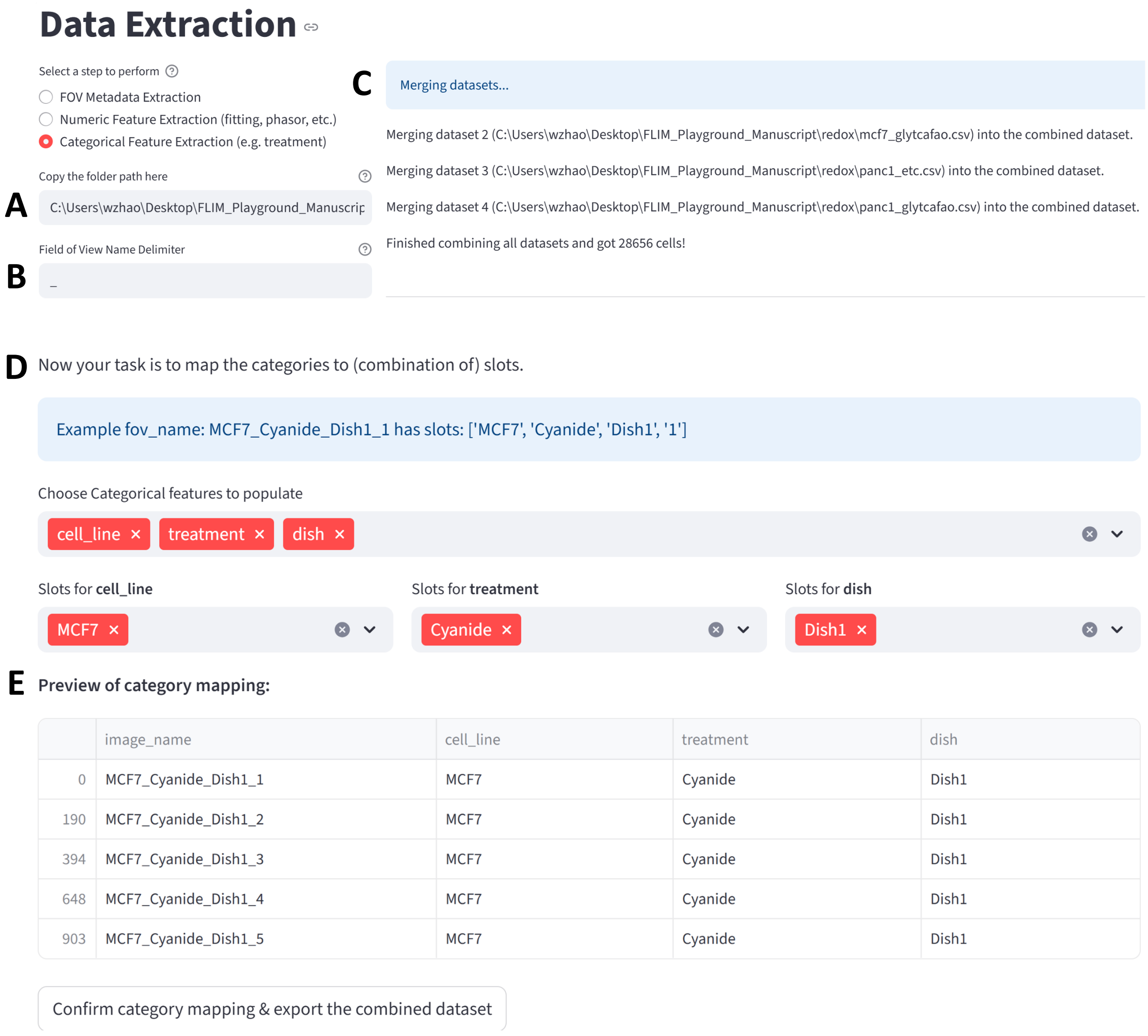

**Supp Figure 5. Categorical feature extraction of PANC-1 and MCF7 single-cell numerical datasets.** **(A)** The folder path containing single-cell numerical datasets extracted by the previous stages of Data Extraction was pasted into the text box. **(B)** The FOV identifiers were segmented by the user chosen delimiter. **(C)** Datasets were merged and the merging status was displayed. **(D)** FLIM Playground parsed the FOV identifiers and interactively assisted the user to assign categorical features to each cell. **(E)** A live preview of such assignment was displayed, and the merged and categorical feature annotated dataset was exported.

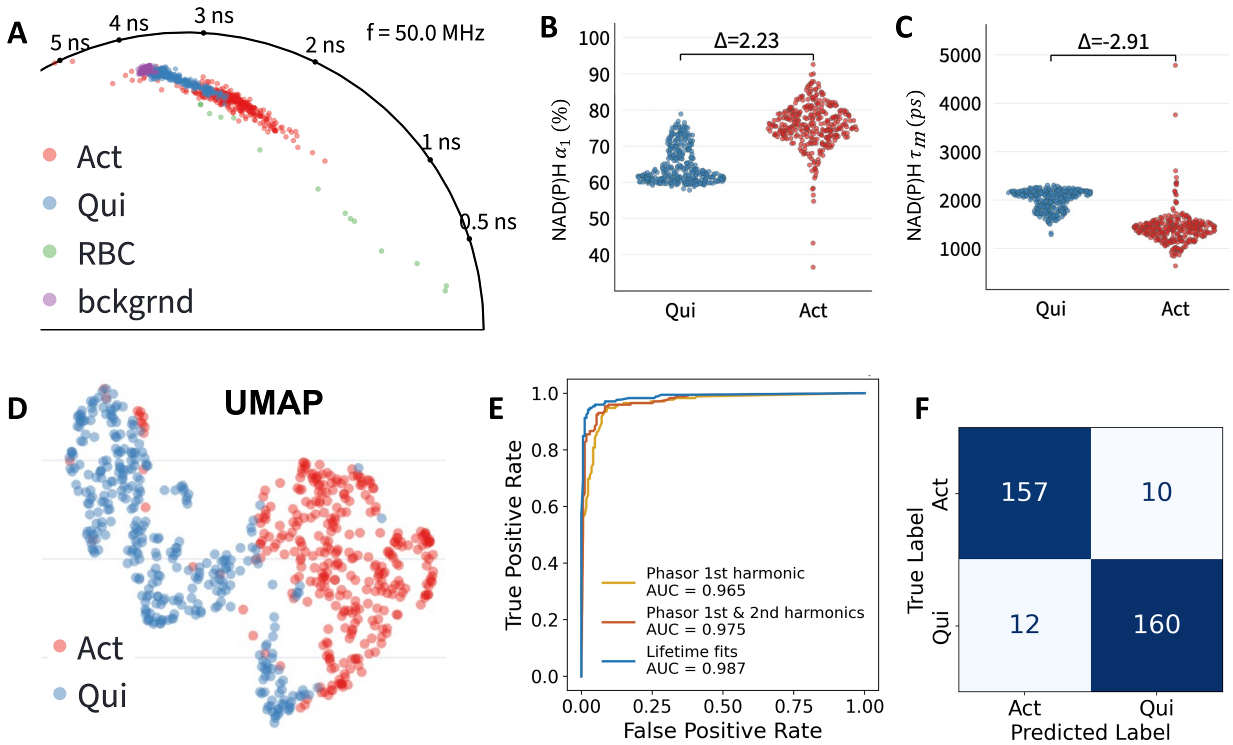

**Supp Figure 6. Discriminating Quiescent versus Activated Primary Human T Cells. (A)** Phasor representation of NAD(P)H lifetime from the primary human T cells measured in flow. The activated cells cluster separately from the control quiescent cells and present with a shorter fluorescence lifetime. Red blood cells (RBCs) have shorter fluorescence lifetimes, and background (bckgrnd) has a longer fluorescence lifetime. Activated cells have **(B)** increased proportion of free ($\alpha_1$) and **(C)** shorter NAD(P)H mean lifetime ($\tau_m$). **(D)** UMAP projection of the NAD(P)H lifetime fit parameters ($\tau_m$, $\tau_1$, $\tau_2$, $\alpha_1$) and phasors (1st, 2nd harmonics) shows separate clustering of the quiescent and activated T cells. **(E)** Random forest classifiers trained on NAD(P)H phasor or lifetime parameters (50% training; 50% testing split) can predict the activation state of the T cells with good accuracy as demonstrated by the area under the curve (AUC) of the receiver operating characteristic (ROC) curves. **(F)** Confusion matrix for the classifier based on NAD(P)H phasors.

**Supp Table 1: Extracted single-cell numerical features from PANC-1 and MCF7 cells across all treatments, computed separately for NAD(P)H and FAD channels.** The features were grouped by selected feature extractors: Lifetime Fit, Lifetime Phasor, Morphology, and Texture. For the FAD channel, the same feature set as NAD(P)H was used, with fit and phasor features derived from FAD-specific decay files and texture descriptors from single-cell FAD intensity images. Since both channels share the same ROI masks, morphology features are identical across channels and therefore omitted for FAD.

| Acquisition Channels | Lifetime Fit | Lifetime Phasor | Morphology | Texture |
|---|---|---|---|---|
| NAD(P)H | $\alpha_1$, $\tau_1$, $\tau_2$, $\tau_m$ | $g$, $s$, $g_2$, $s_2$, $m$, $\phi$, $\tau_\phi$, $\tau_m$ | area, perimeter, solidity, eccentricity, major_axis_length, minor_axis_length, circularity | Granularity: 1,3,5,7,9, Radial Distribution: 1,2,3,4, Mass Displacement, Intensity Sum |
| FAD | $\alpha_1$, $\tau_1$, $\tau_2$, $\tau_m$ | $g$, $s$, $g_2$, $s_2$, $m$, $\phi$, $\tau_\phi$, $\tau_m$ | None | Granularity: 1,3,5,7,9, Radial Distribution: 1,2,3,4, Mass Displacement, Intensity Sum |

### *Supplemental Manual*

**19 Classification** **92**

### 1 Quick Start

Welcome to the FLIM Playground 🥳🎉🥂! This is an interactive graphical user interface (GUI) that allows you to extract single-cell features from fluorescence lifetime imaging microscopy (FLIM) raw data (Data Extraction) and analyze extracted features or datasets extracted via other methods using a built-in repertoire of methods (Data Analysis).

To quickly try out different analysis methods, download this sample dataset and try the online demo. If you prefer to use your own data, read this chapter on data analysis configuration to learn how to configure the system.

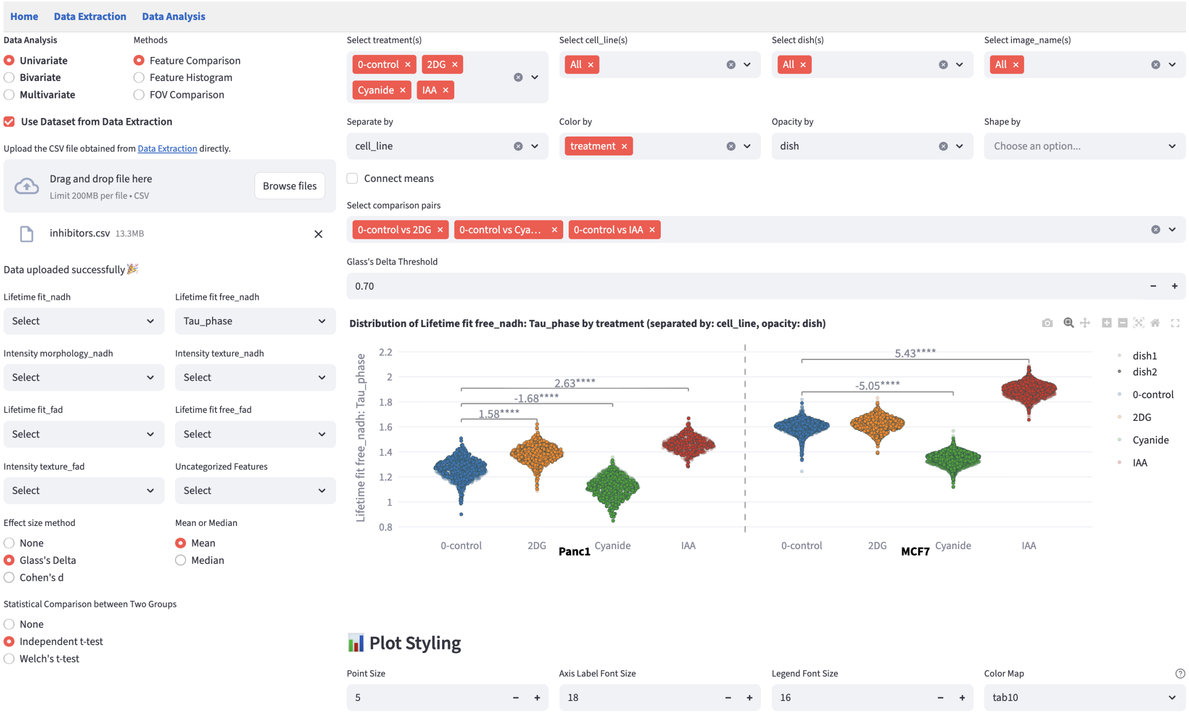

Due to the limitation in local file system access, extracting features from raw data is not available in the online demo. Read on to discover how FLIM Playground transforms raw fluorescence decay data into biological insight through an integrated, interactive workflow.

The latest version of this FLIM Playground manual is available online.

#### 1.1 Installation

FLIM Playground is built entirely in Python and is open-source.

##### 1.1.1 Download

Download the desktop app from GitHub and double-click it to run. Releases are currently available for Windows 11 and Mac OS 26. If your operating system is not either of these, you can build it yourself by following the instructions below.

##### 1.1.2 Build it yourself

1. Clone the repo and navigate into the repository once cloned.
2. Install the Python environment
   - Install uv if not yet installed
   - run `uv sync`
3. Build the app
   - run `pyinstaller Flim-Playground.spec --clean`

#### 1.2 Introduction

Fluorescence lifetime imaging microscopy (FLIM) measures the time it takes for a fluorescent molecule to emit light (return to the ground state) after being excited by a pulse of light (enter the excited state). It is sensitive to changes in fluorophore microenvironment including, pH, temperature, and conformational changes due to protein-binding and the presence of quenchers[1]. Coupled with modern automated cell-segmentation methods[2], FLIM enables single-cell analyses that can reveal biological heterogeneity.

> **i Instrumentation**
>
> To acquire FLIM data, a light source—typically a pulsed laser for time-domain methods or a modulated continuous-wave source for frequency-domain methods—is used to excite the fluorophore of interest. The emission is detected using instrumentation capable of resolving fluorescence decay, such as time-correlated single-photon counting (TCSPC), time-gated, or phase/modulation-based detection. In time-domain FLIM, the delay between excitation and photon arrival is measured, and often a histogram is built, with the x-axis representing the delay time and the y-axis representing the number of photons falling into each time bin. Compared to intensity images, FLIM has an

additional dimension of time (e.g., 256 time bins per 12.5 nanoseconds).
In frequency-domain FLIM, the phase shift and modulation depth of the emission relative to the excitation are determined.

#### 1.3 Challenges

A diverse set of tools — both open-source and commercial, ranging from libraries to code-free graphical user interfaces (GUIs) — is available to extract and analyze FLIM data, providing alternative methods and therefore flexibility to FLIM researchers. Examples include PhasorPy[3], an open-source library for analyzing fluorescence lifetime using the phasor approach; FLUTE[4], an open-source GUI for interactive phasor analysis; FLIMPA[5], an open-source phasor analysis GUI enabling batch processing, ROI-based quantification, and experiment-level comparison through manual assignment; FLIMLib, an open-source generic curve fitting library that can be used to fit fluorescence lifetime decay data; SPCImage[6], a commercial software for fitting and phasor features.

However, while some tools offer code-free interfaces, users still need to write custom code—either to prepare data in the proper format as input, or to further process their outputs for downstream analysis. The fragmentation between tools arises because each focuses on only a subset of data levels: pixel, cell ROI, channel, field of view, and experiment.

##### 1.3.1 Data Levels

- **Pixel**:
  - A single decay curve encoded in vendor-specific file formats (e.g., Becker & Hickl, PicoQuant, etc.)
- **Region of Interest (ROI)**
  - Mask with cell labels
- **Channel**
  - Different fluorophores
  - Fluorophore-specific calibration files
  - Masks focusing on different parts of the cell (e.g., whole cell, cytoplasm, nucleus, stain, etc.)
  - Different feature extraction methods (fitting, phasor, morphology, texture)
- **Field of View (FOV)**
  - Input decay types (e.g., 2D, 3/4D, pixel-level lifetime features already fitted)
- **Experiment**

– Different treatments, time points, cell lines, etc., and combinations thereof

An integrated framework should take into account all data levels ■ ■ ■ ■ ■ (•) while maintaining the flexibility to handle various input types (◦). **It should provide a level of abstraction to address fragmentation from data levels and input types**.

Additionally, the use of FLIM is rapidly evolving, and new methods are being developed all the time. An integrated framework should allow users to choose among alternative methods seamlessly, be the backbone of iterative explorations integral to research, and be ready to incorporate new methods: **The provided level of abstraction should address fragmentation from extraction and analysis methods**.

Finally, many of the existing tools, including GUI-based software, are not cross-platform, which limits their accessibility. The Installation addresses this last challenge.

A closer parallel to FLIM Playground's integrated approach is the combination of CellProfiler[7], which extracts per-object morphological and texture features, and CellProfiler Analyst[8], which ingests these outputs or other feature tables for visualization and statistical analysis. However, FLIM Playground stands apart in that it can extract lifetime features from time-resolved data, alongside morphological and texture features from intensity images. Its general Data Analysis section incorporates FLIM-specific methods (e.g., phasor analysis) and provides statistical models that address additional analysis aspects such as data heterogeneity, in addition to training machine learning classifiers.

#### 1.4 Method

##### 1.4.1 Feature Classes

Tabular data columns can be categorized into three feature classes:

- Identifiers: unique row identifier and (optional) field of view identifier that allow us to investigate the biological heterogeneity at the single-cell level
- Categorical features: conceptually help us group the rows (e.g., `treatment` will group the rows into different treatment groups)
- Numerical features: quantify the differences/similarities between data groups

**Science, from the data perspective, is about closing the conceptual categorical gaps with quantitative measurements.**

At data levels, the data are processed to extract the feature classes in Data Extraction, and the classes are used in the Data Analysis section.

Increasing complexity

| Data Levels | Data Processing | Feature Classes |
|---|---|---|
| Experiment | Assign category labels (e.g., treatment, cell line, etc.) to each cell; perform data analysis on labeled cell groups. | **Categorical features** |
| Field of View | Assign FOV identifier to each cell, obtain unique cell identifiers | **Identifiers** |
| Acquisition Channel | Apply **per-channel** ROI mask, calibration and extraction methods to cell-level signals | **Numerical features** |
| Cell ROI | Aggreate pixel-level decays to obtain cell-level signals | N/A |
| Pixel | Read pixel-level decays encoded in different formats | N/A |

#### 1.4.2 Design

FLIM Playground has two independent sections:

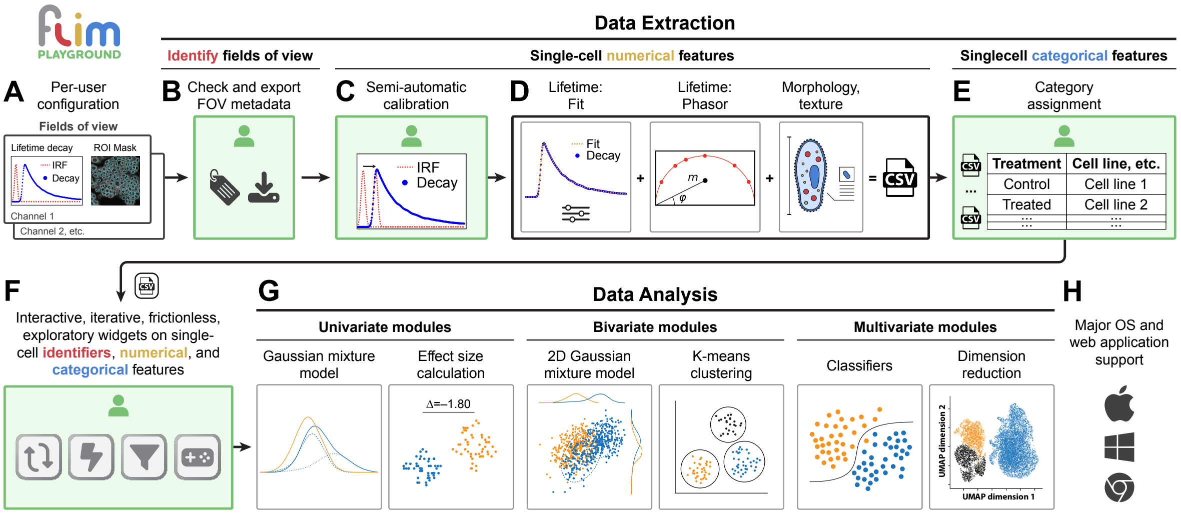

##### 1.4.2.1 Data Extraction

Data Extraction extracts single-cell features from the raw data. It adopts a framework that offers channel-level flexibility in input types and extraction methods without incurring too much overhead for users. Following the above categorization, it is divided into the following steps:

- Data Extraction Configuration (A): allows users to choose among alternative input types and extraction methods (extractors).
- Metadata organization (B): extracts the field of view identifiers and their configurations
- Numerical Feature Extraction: extracts single-cell numerical features based on user-selected extractors. More extractors can be integrated in the future.
  - Calibration (C): calibrate for IRF shift or use fluorescence lifetime standard
  - Alternative lifetime extractors (D):
    * Fitting
    * Phasor
  - Alternative intensity-based extractors (D):
    * Morphology
    * Texture
- Categorical feature extraction (E): extracts single-cell categorical features and combines experiment-level datasets.

###### 1.4.2.2 Data Analysis

Data Analysis analyzes features—whether extracted through Data Extraction or by other methods—using visualizations and statistical modeling. It deploys a shared framework (F) built to handle the feature classes across all analysis methods, enabling the same interactive and frictionless exploration experience and allowing new methods to be integrated in the future easily.

- Data Analysis (G) goes in-depth into how FLIM Playground handles the feature classes and Data Analysis Config goes through how users can configure FLIM Playground to analyze datasets that are not extracted by Data Extraction.

Here is the list of analysis methods incorporated into FLIM Playground, grouped by the number of numerical features they take as inputs:

- Univariate analysis
  - Feature Comparison
  - Feature Histogram
  - Field of View Comparison
- Bivariate analysis
  - Feature Distribution
  - Phasor Analysis
- Multivariate analysis
  - Dimension Reduction

– Classification

Both sections are built in Python and as self-contained executables ready to run on major operating systems and in browsers (**H**).

##### 1.4.3 Summary

**FLIM Playground** resolves these challenges with an interactive code-free graphical user interface (GUI) that spans the full pipeline. It integrates validation checks that guide users at every step, and has a built-in repertoire of analytical methods with interactive widgets that encourage hypothesis-driven, iterative exploration of large datasets. It is built on a modular architecture that enables incorporation of new algorithms in the future.

### Part I

### Data Extraction

### 2 Overview

FLIM Playground transforms raw FLIM data into single-cell numerical and categorical features, ensuring smooth integration with downstream analysis. All steps are seamlessly connected through interactive widgets, with built-in error checking and reporting to ensure correctness.

A high-level overview of the data extraction process is shown below.

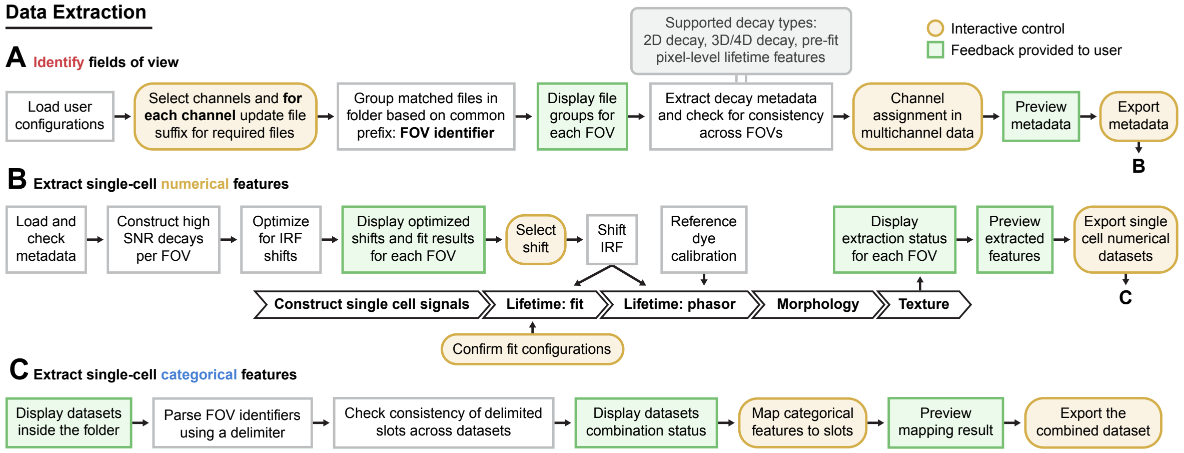

#### 2.1 Stages

Users can configure the system once and apply it to future data. After configuration, users can extract features from raw data using the following steps, inspired by the categorization:

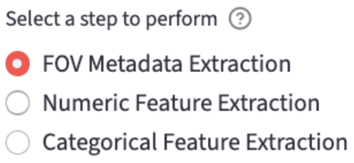

1. FOV Metadata Organization (A): each experiment session consists of multiple fields of view (FOVs). This step helps users organize the metadata of the FOVs.

2. Numerical feature extraction (B): it calibrates and then extracts single-cell numerical features.

   - Calibration
     - Fit Calibration
     - Fit Free Calibration
       - IRF Shift
       - Fluorescence lifetime standard
   - Per-channel feature extraction includes:
     - Lifetime fitting features: fit an exponential decay curve to the measured decay and extract lifetime features per cell ROI.
     - Phasor features: calculate the phasor features such as the phasor coordinates. It also provides phasor analysis.
     - Morphology features: calculate morphology features based on the ROI mask.
     - Texture features: calculate texture features based on the intensity image and the ROI mask.

3. Categorical feature extraction (C): extract the categorical features such as the treatment, time point, etc. based on the FOV name.

- Users can then upload the extracted datasets to perform Data Analysis.

#### 2.2 Input File Types

##### 2.2.1 Decay

See decay types for more details.

- 2D decay:
  - `.csv`: a tabular data sheet with each row representing a cell and each column representing a time bin.
- 3D/4D decay: besides the spatial dimensions and the time dimension, the additional dimension in the 4D array represents acquisition channels.
  - `.sdt`: Becker & Hickl
  - `.ptu`: PicoQuant

##### 2.2.2 Mask

Because of the single-cell focus of FLIM Playground, cell-level masks are needed for each channel. Channels can share the same mask, or use different masks (when they use different masks, the mask ID for each cell should match across masks).

The cell-level mask can focus on different parts of the cell region, such as the whole cell, the cytoplasm, the nucleus, or the stain part.

For an example, see here.

- `.tiff`/`.tif`: a 2D array with background labeled as `0` and each cell ROI region labeled as a unique positive integer. That integer will be part of the unique cell identifier.

##### 2.2.3 IRF

IRF must be a 1D array. Currently, the supported formats are (extendable to more formats in the future):

- `.txt`: it uses `np.loadtxt` to parse a txt file. It returns a 1D array if the IRF numbers are in one row or in one column.

##### 2.2.4 SPCImage t1

If the decay type is 3D/4D pixel-prefitted, users can provide pixel-prefitted lifetime feature files.

- `.asc`: a 2D array in spatial dimensions outputted by SPCImage, with each (row, column) containing the value of the lifetime feature (e.g. `t1`: the first component's lifetime) of that pixel.
- Depending on the chosen number of lifetime components, the other pixel-prefitted feature files are included and their names are inferred from the `t1` file, by replacing `t1` in the file name with `t2`, `t3`, `a1[%]`, etc.

##### 2.2.5 Fluorescence Lifetime Standard

If the fluorescence lifetime standard-based calibration is chosen, users are expected to provide the fluorescence lifetime standard file.

- `.tiff`/`.tif`: a 3D array, with one dimension representing the time axis.

##### 2.2.6 Intensity (2D)

If the channel's imaging modality is `Intensity-only`, users are expected to provide a 2D intensity image in:

- `.tiff`/`.tif`: a 2D array.

#### 2.3 Limitations

- To simplify the workflow and due to the framework chosen, FLIM Playground does not support pixel-level fitting and fit-free analysis, and there are many other open-source tools that can do so (for a comprehensive list, see here). Instead, it does cell-level fitting and fit-free (phasor) feature extraction by summing up all the decays belonging to the same cell ROI to get one cell-level decay as a preprocessing step[9]. It also accepts prefitted pixel-level lifetime features from SPCImage and aggregates pixel-level lifetime features to cell-level lifetime features.

### 3 Configuration

> **Note**
>
> Users can configure the system once, and it will be applied to future data.

Constructing a framework that balances flexibility and usability is about choosing the right abstraction level. FLIM Playground chooses acquisition *channel* from data levels as the focus and adopts the **channel-centric framework**. Settings are divided into two categories: cross-channel configurations and per-channel configurations.

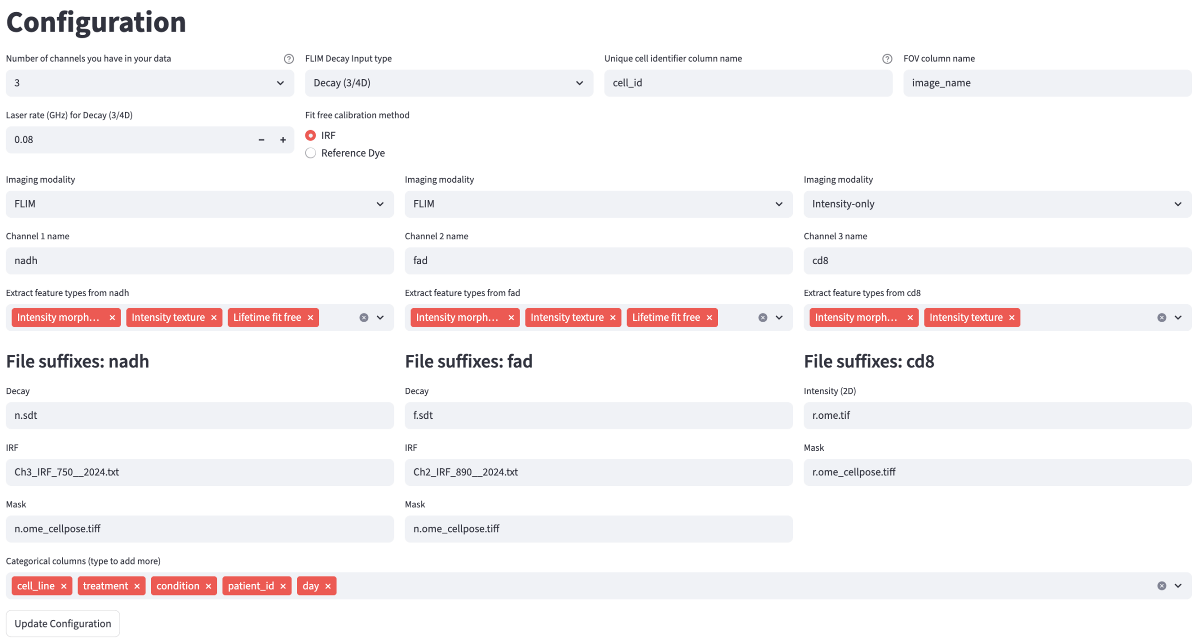

#### 3.1 Cross-Channel Configurations

##### 3.1.1 Categorical Features

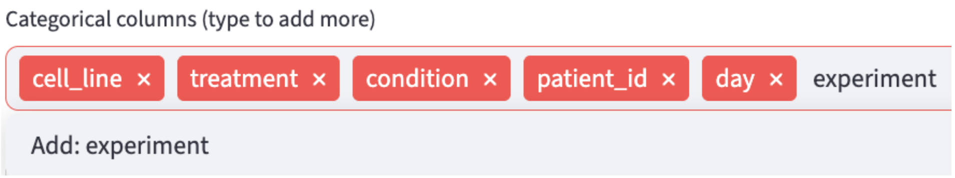

Categorical features are useful to organize the data into groups of interest, and will be used extensively in the Data Analysis section. Users can specify the potential categorical feature names to be extracted in the Categorical Feature Extraction step.

##### 3.1.2 Decay Types

FLIM Playground supports different decay types, including 2D decay, 3D/4D decay, and 3D/4D pixel-prefitted decay, through user specification.

###### 3.1.2.1 2D Decay

In the system developed by Samimi et al.[10], single cells flow through and a decay curve is acquired for each cell by aggregating all the photon arrival time delays deemed to be from the same cell. In exchange for spatial distribution, the acquisition speed is increased enormously. The output of the system is a tabular data sheet with each row representing a cell and each column representing a time bin. **FLIM Playground currently supports 2D decay in the CSV format.**

###### 3.1.2.2 3D/4D Decay

In TCSPC-based FLIM, the data is stored in a 3D/4D array and in special format (e.g. `.sdt`, `.ptu`). In addition to the spatial dimensions and the time dimension (`XYT`), the channel dimension may also be present, making it 4-dimensional (`CXYT`). **FLIM Playground currently supports both `.sdt` and `.ptu` files up to those 4 dimensions.** It reads the `duration` and the number of `time bins` from the decay file metadata. Some systems can also record a time stack, which is not supported yet.

###### 3.1.2.3 3D/4D pixel-prefitted

Since FLIM Playground currently does **not** support pixel-fitting, users have the option to provide pixel-prefitted outputs from SPCImage in the format of `.asc`. See here for how FLIM Playground processes SPCImage outputs to obtain cell-level lifetime fitting features.

##### 3.1.3 Identifiers

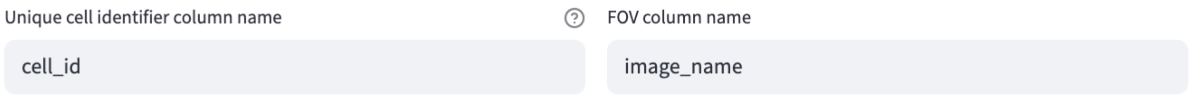

###### 3.1.3.1 Cell Identifier

Users can specify the column name for the unique identifier of the cell to be stored in the output dataset. The cell identifier is constructed as `{FOV Identifier}_{cell_label}`, where the `cell_label` is the unique integer label in the mask or the row number if the decay type is 2D decay.

###### 3.1.3.2 FOV Identifier

Users can specify the column name for the field of view identifier. If the decay type is 2D decay, users may want to specify the column name as `exp_name`, because the FOV is not an image.

##### 3.1.4 Decay Info

###### 3.1.4.1 Laser frequency

Users can specify the laser frequency in GHz. For example, 0.08 GHz equals 80 MHz. It will be used in the phasor analysis. If you do not intend to use the phasor analysis, you can fill in any number.

###### 3.1.4.2 2D Decay-specific

Since the 2D decay file does not provide the `duration` and the number of `time bins` per laser pulse interval, users need to specify them.

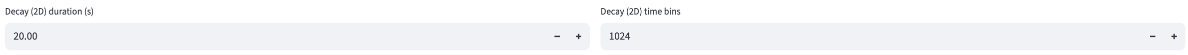

##### 3.1.5 Calibration Method

It supports two calibration methods for phasor analysis:

- IRF shift-based calibration
- fluorescence lifetime standard-based calibration

Since the instrumental response function (IRF) or the fluorescence lifetime standard is measured for each channel, they are set in the per-channel configurations. The fluorescence lifetime standard's lifetime is channel-agnostic.

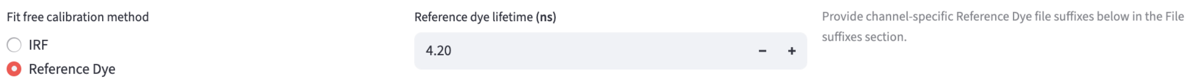

##### 3.1.6 Number of Channels

Each FOV is assumed to have the same number of channels. The configuration will dynamically adjust the per-channel configurations based on the specified number of channels. Currently, the maximum number of channels is 4, and it can be extended to house more channels.

#### 3.2 Per-Channel Configurations

##### 3.2.1 Channel Name

Users can customize the channel name based on, for example, the name of the fluorophore.

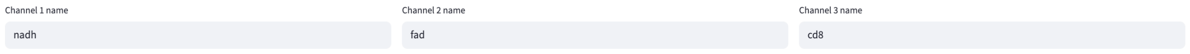

##### 3.2.2 Imaging Modality

Users can specify the imaging modality for each channel. Currently, two modalities are supported:

- `FLIM`: The signal is time-resolved, i.e., has a time dimension, and is expected to be from one of the decay types.
- `Intensity-only`: The signal is a spatial map of intensity values (2 spatial dimensions), i.e., has no time dimension. **Currently, the only supported signal type is a 2D intensity image in the format of `.tiff`/`.tif`.**

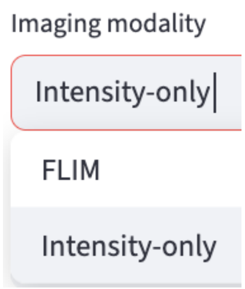

Tip

This setup allows users to image some channels in FLIM mode, and others in non-FLIM mode. It also allows users to extract and analyze their data if they image in non-FLIM microscopic systems such as epifluorescence, confocal, brightfield, etc. by selecting `Intensity-only` for all channels.

Note

If the decay type is 2D decay, the imaging modality for all channels is restricted to `FLIM`.

##### 3.2.3 Feature Extractor

Four types of feature extractors are available if the imaging modality is `FLIM`:

- Lifetime fit
- Lifetime fit free
- Intensity Morphology
- Intensity Texture

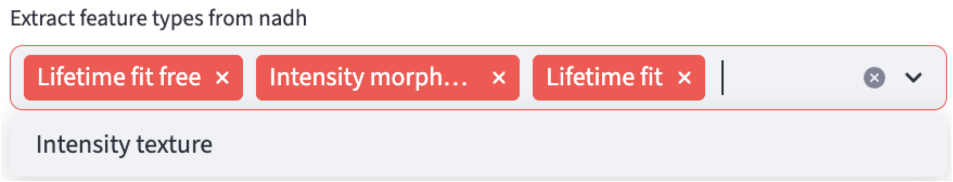

Note

If the decay type is 2D decay, the Intensity Morphology and Intensity Texture feature extractors are **not** applicable.

The Intensity Morphology and Intensity Texture feature extractors are available if the imaging modality is `Intensity-only`.

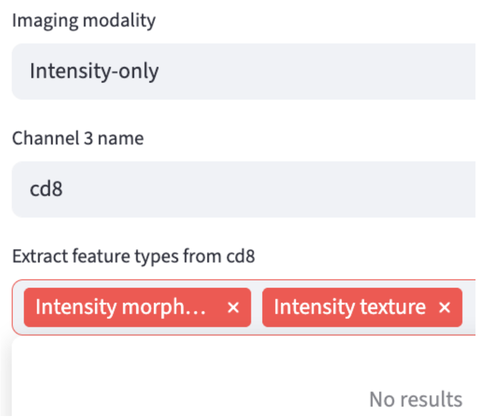

Each channel can be assigned its own set of feature extractors. For example, if users do not intend to do lifetime extraction, they can de-select the `Lifetime fit` and `Lifetime fit free` options.

##### 3.2.4 Number of Components

If users select the `Lifetime fit` under this channel, they can specify the number of components to fit.

##### 3.2.5 File Suffix

The FOV Metadata Organization step looks for associated input files for all channels of a given FOV using file suffixes. Based on the decay type, calibration method, and the selected feature extractors, the system automatically generates the file suffixes for required input files.

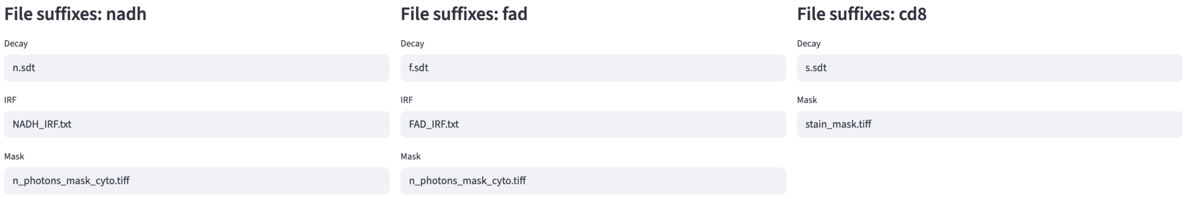

In the example above, channels `nadh` and `fad` were asked to provide an IRF file, while channel `cd8` did not, because `Lifetime fit` was selected for the former two channels and not for the latter. The file suffixes of the IRF files were different for each of the two channels, but they shared the same ROI mask file suffix because they used the same ROI mask. The `cd8` channel used a different ROI mask than the other two channels, indicated by the mask file suffix.

For detailed information on what file formats are supported for each file type (Mask, IRF, etc.), see input file types.

For detailed information on how the file suffixes are used, and what each file type (e.g. `IRF`, `Mask`, etc.) is, see the FOV Metadata Organization page.

#### 3.3 Save

Finally, users can save the configuration for future use by clicking the `Update Configuration` button.

### 4 Field of View Metadata

It is the first step in the Data Extraction workflow. It organizes the metadata of each field of view (FOV)—including file paths for all required inputs and decay information—into a single metadata file. This file is then used for numerical feature extraction, as well as for users' own bookkeeping and troubleshooting.

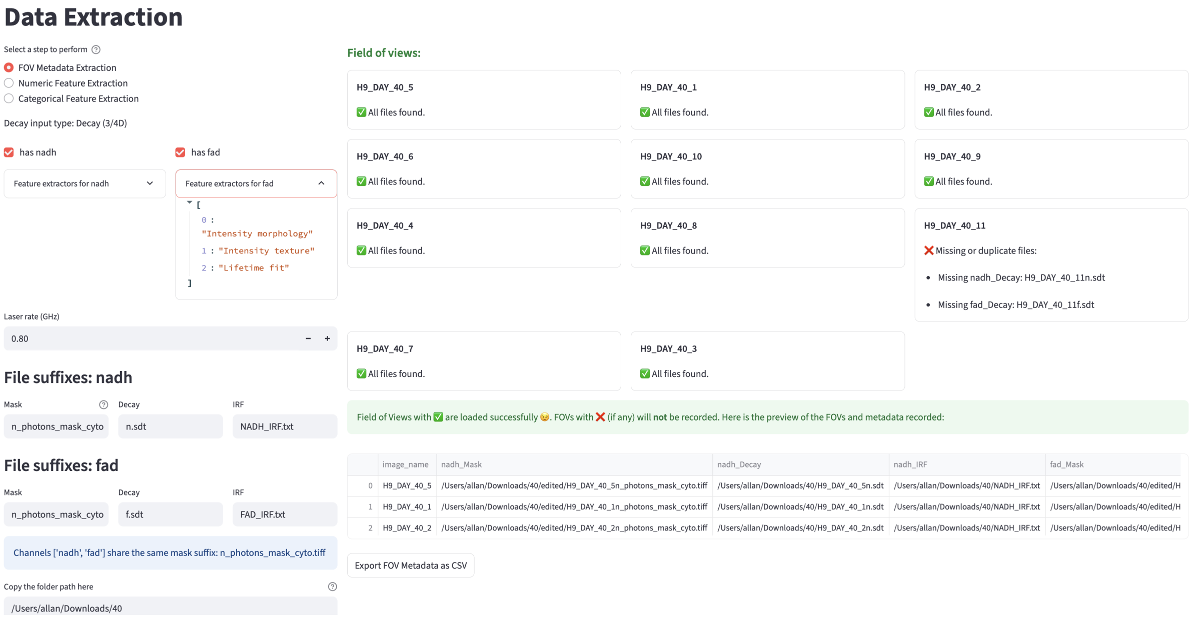

It is divided into two sections:

- The left 1/3 of the screen is the metadata configuration panel, inherited from the Data Extraction Configuration step.
- The right 2/3 shows the extraction status for all FOVs.

#### 4.1 Metadata

Metadata are configured earlier in the Data Extraction Configuration step and copied here. Some fields remain editable for flexibility, while others are fixed to ensure usability. Non-editable fields can be modified in the Data Extraction Configuration step. After saving your modifications, return to this page and refresh it to see the changes.

##### 4.1.1 Decay Type

See decay types for more details. The decay type is not editable here.

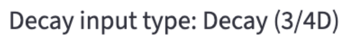

##### 4.1.2 Channel Names

See channel name config for more details. The names are not editable here, though users can choose which channel to include.

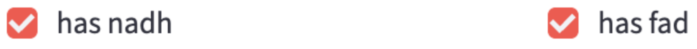

##### 4.1.3 Feature Extractors

See feature extractor config for more details. The feature extractors for each channel are shown but not editable here.

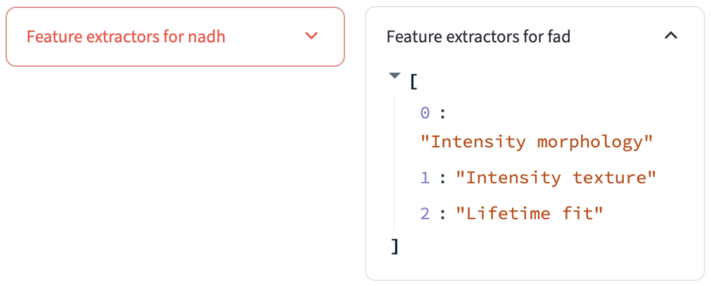

##### 4.1.4 Decay Info

See decay info config for more details. The decay information is editable here.

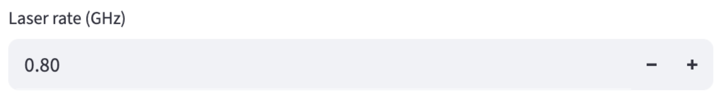

###### 4.1.4.1 2D Decay-specific

Since the 2D decay file does not provide the `duration` and the number of `time bins` per laser pulse interval, users need to specify them.

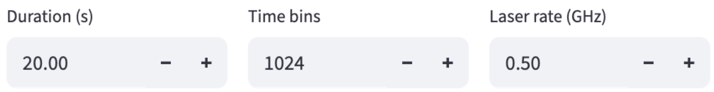

##### 4.1.5 File Suffix

Copied from the file suffix config. Users can edit the file suffixes here.

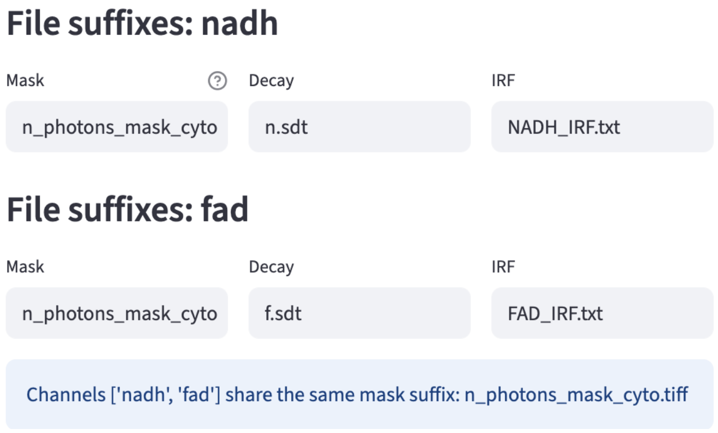

##### 4.1.6 Fluorescence Lifetime Standard

If the selected fit free calibration method is fluorescence lifetime standard, users are asked to specify the fluorescence lifetime standard file path for each channel and the fluorescence lifetime standard's lifetime, which are copied from the data extraction configuration.

##### 4.1.7 Folder Path

Finally, users can specify the folder path that contains all the required input files.

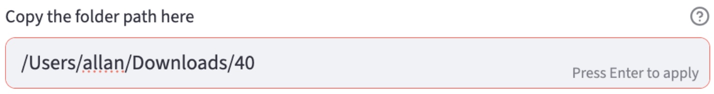

#### 4.2 Metadata Extraction

##### 4.2.1 FOV File Paths

The first step is to find the fields of view:

- It finds all the files recursively in the folder path that ends with the 1st file suffix of the 1st channel.
- The prefix of all the matched files is considered to be the field of view name.
  - FOV name = file name - 1st file suffix

Then, it uses the found FOV names to find all other files:

- The file name to search = FOV name + other file suffixes for each file suffix

The only exceptions are the calibration files, because they are not FOV-specific. They will be searched for based on their file suffixes only.

###### 4.2.1.1 Success

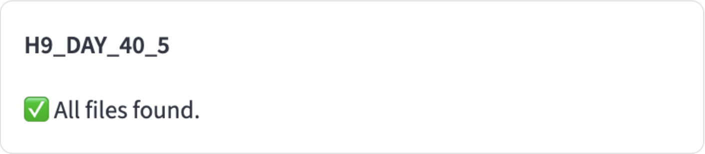

###### 4.2.1.2 Missing

When it cannot find the file based on the file name (FOV name + file suffix for that file type) inside the folder path, it will complain.

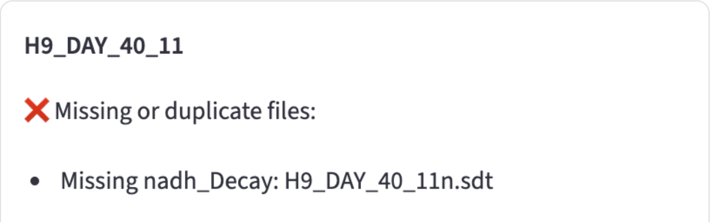

###### 4.2.1.3 Duplicate

Because the calibration file is not FOV-specific, it will be searched based on the file suffixes only. If there are multiple files that match the same file suffix, FLIM Playground will be confused about which one to use.

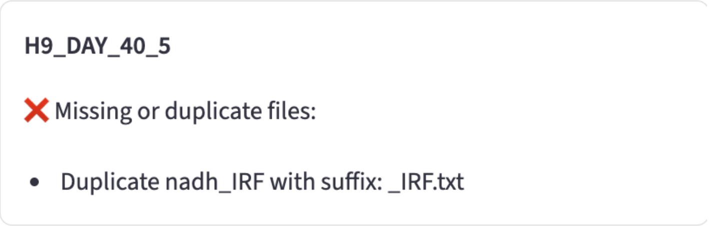

Fields of view with ✅ are loaded successfully 🎇. FOVs with ❌ (if any) will not be recorded.

> **Tip**
>
> If the issue persists after renaming the files to match the required prefix, force-quit and relaunch FLIM Playground to clear its internal cache.

##### 4.2.2 Decay Info

If the decay type is 3D/4D decay, FLIM Playground will try to infer the `duration` and the number of `time bins` per laser pulse interval from the decay file metadata. It will also check for inconsistencies across all decay files.

###### 4.2.2.1 Channel Assignment

If the decay file associated with the channel found by the previous step is a 4D array, then FLIM Playground will try to infer the channel number for that channel. For each channel name, it will list all non-empty channels as potential channels to be assigned. If there is only one, the assignment is automatic. If there are multiple, users need to select the channel intended. It will also check for inconsistencies across all decay files about their dimensions and complain if there are any.

###### 4.2.2.2 Fluorescence Lifetime Standard

If the file path to the fluorescence lifetime standard is specified, FLIM Playground will try to look for and read the file. It expects a `tiff` or `tif` file that contains a 3D array, with one of them as the time dimension. It will try to match the time dimension with the time dimension of the decay file. If it cannot find the matching time dimension, it will complain.

Error: Reference dye file 'test.tif' not found in folder. Please check the file name.

Error: Cannot find the time axis (256 time bins) in the reference dye file dimensions: (56, 512, 512)

Once the time axis is matched, the file path to the fluorescence lifetime standard, the fluorescence lifetime standard's lifetime, and the time axis will be recorded.

##### 4.2.3 Result Preview and Export

If there is a non-zero number of FOVs with ✅ and channel assignment and fluorescence lifetime standard checks (if applicable) are passed, a preview of the metadata data sheet will be shown for users to check.

| | image_name | nadh_Mask | nadh_Decay | nadh_IRF | fad_Mask |
|---|---|---|---|---|---|
| 0 | H9_DAY_40_5 | /Users/allan/Downloads/40/edited/H9_DAY_40_5n_photons_mask_cyto.tiff | /Users/allan/Downloads/40/H9_DAY_40_5n.sdt | /Users/allan/Downloads/40/NADH_IRF.txt | /Users/allan/Downloads/40/edited/H |
| 1 | H9_DAY_40_1 | /Users/allan/Downloads/40/edited/H9_DAY_40_1n_photons_mask_cyto.tiff | /Users/allan/Downloads/40/H9_DAY_40_1n.sdt | /Users/allan/Downloads/40/NADH_IRF.txt | /Users/allan/Downloads/40/edited/H |
| 2 | H9_DAY_40_2 | /Users/allan/Downloads/40/edited/H9_DAY_40_2n_photons_mask_cyto.tiff | /Users/allan/Downloads/40/H9_DAY_40_2n.sdt | /Users/allan/Downloads/40/NADH_IRF.txt | /Users/allan/Downloads/40/edited/H |

Export FOV Metadata as CSV

As an example, the following metadata are extracted:

- `image_name`: The `image_name` column is the name of each FOV, which is specified in the fov identifier config.
- `nadh_Mask`, `nadh_Decay`, `nadh_IRF`, `fad_Mask`, `fad_Decay`, `fad_IRF`: the file paths of the mask, decay, and IRF files for the `nadh` and `fad` channels.
- `nadh_input_type`, `nadh_imaging_modality`, `fad_input_type`, `fad_imaging_modality`: the decay type for each channel. If they share the same imaging modality, the input types should be the same. The only imaging modality supported by FLIM Playground is `FLIM`, therefore the input_type for all channels should be from one of the decay types.
- `nadh_Lifetime fit free`, `nadh_Intensity morphology`, `nadh_Lifetime fit`, `fad_Intensity morphology`, `fad_Intensity texture`, `fad_Lifetime fit`: feature extractors for each channel.
- `nadh_channel`, `duration`, `fad_channel`, `time_bins`, `laser_rate`: the decay info for each channel and info shared by all channels.

The user can click the `Export FOV Metadata as CSV` button to export the metadata as a CSV file.

### 5 Numerical Feature Extraction

In this step, FLIM Playground extracts *single-cell* numerical features from the raw data in a folder. For each channel, the features to be extracted are specified in the user-selected feature extractors. The available feature extractors are:

- Lifetime fit: fit exponential models to decay curves to estimate lifetimes and their fractional contributions by minimizing an objective function.
- Lifetime fit free: transform decay curves to phasor space using Fourier transform to obtain phasor coordinates.
- Intensity Morphology: morphological features of single-cell ROI masks
- Intensity Texture: texture features of single-cell ROI intensity images

#### 5.1 Input

A CSV file extracted in the fov metadata extraction step that contains the metadata of the FOVs including:

- required file paths
- decay type
- decay info
- selected feature extractors for each channel
- fit free calibration method (if fit free calibration is required)

Users can either upload a previously extracted metadata file (left), or use the cached metadata file just extracted in the fov metadata extraction step (right).

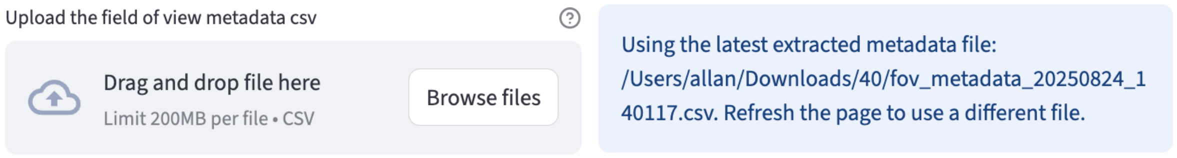

For `Lifetime fit` and `Lifetime fit free` feature extractors, the first step is always calibration.

#### 5.2 Calibration

Calibration in FLIM is essential because raw decays are convolved with the instrument response function (IRF)—the timing profile of the detection system's response to an ultrashort light pulse that broadens and shifts the measured decay—so without correcting for these, fitted lifetimes and phasor positions are biased and not comparable across days, samples, or instruments.

##### 5.2.1 Fit Calibration

During different experiments, the IRF may shift differently with respect to the measured decay. Before applying reconvolution fitting, the IRF shift needs to be estimated for each channel if the `Lifetime fit` feature extractor is selected for the channel.

It is performed in two steps:

1. Gather the high signal-to-noise ratio (SNR) decay curves
2. For each curve, perform reconvolution fitting and set the shift value as the free parameter to be optimized/fitted.

The distribution of the shift values is displayed as an interactive scatter plot. When clicking on a point, the corresponding decay curve with the fitted curve and the key statistics are displayed for diagnostics. Based on the distribution or using certain prior knowledge, users can specify the shift value applied to all fields of view (if `Fix the Shift` is selected, see fitting options), or use the fov-specific shift value found by the fitting (if `Fix the Shift` is deselected).

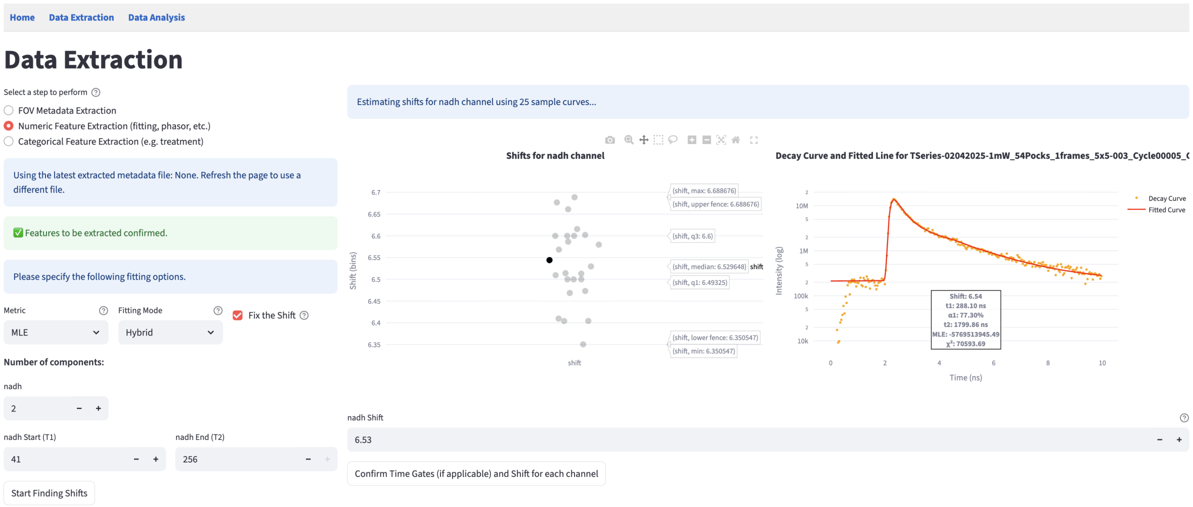

###### 5.2.1.1 Gather High SNR Decay Curves

The high SNR decay curves are constructed automatically based on the decay type. If it is in 2D format, a total of the brightest 30 curves that are below `100000` photons are selected and evenly distributed across all fields of view (one CSV file is considered as one FOV). If it is in 3D/4D format, the fields of view are images, and one curve is constructed for each image that includes all the non-zero pixels within the ROI mask.

###### 5.2.1.2 Reconvolution Fitting

Fitting is essentially an optimization problem: it minimizes the difference between the fitted curve and the measured curve. The fitted curve is modeled by an $n^{th}$ component exponential function convolved with the shifted IRF, and the difference is modeled as an objective metric. Therefore, users are provided with controls over two parts of the fitting process through the fitting options panel:

1. How to construct the objective metric
2. How to perform the optimization

Implementation-wise, FLIM Playground uses the `lmfit` package that takes generic objectives to perform the optimization process that is flexible enough to handle the reconvolution fitting.

If at least one channel has `Lifetime fit` feature extractor selected, the fitting options panel is displayed.

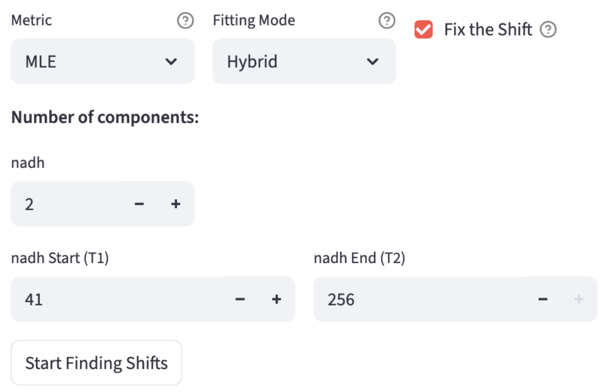

Let's break down the fitting options one by one.

###### 5.2.1.2.1 Number of Components

The number $n$ in the $n^{th}$ component exponential function:

$$I(\mathbf{t}) = \sum_{i=1}^{n} A_i \, e^{-\mathbf{t}/\tau_i}, \qquad \tau_i > 0$$

Therefore, $n$ determines the parameters to be fitted: the amplitudes $A_i$ and the lifetimes $\tau_i$. $\mathbf{t}$ is the time axis of the decay curve calculated by the `duration` and `time bins` from the decay info:

$$\mathbf{t} = [0,\ \Delta t,\ 2\Delta t,\ \dots,\ (N-1)\Delta t], \quad \text{where } \Delta t = \frac{T}{N}.$$

It supports $n = 1, 2, 3$ components.

Additionally, the constant offset parameter ($Z$) that represents a time-independent background (i.e., room light) is also optimized[11].

$$\hat{y}(t) = [I(t) \otimes \mathrm{IRF}(t)] + Z$$

###### 5.2.1.2.2 Time Gates

Due to the dead time of the system certain time bins from the head and/or tail of the decay curve are not reliable. The T1 (head) and T2 (tail) gates are used to select the time range of the decay curve to be fitted. Users can inspect the decay curve by clicking the full-screen mode button of the plot on the right of the shift result. Hover-based interaction is implemented so users can see the time bin numbers to have a better sense of the time range.

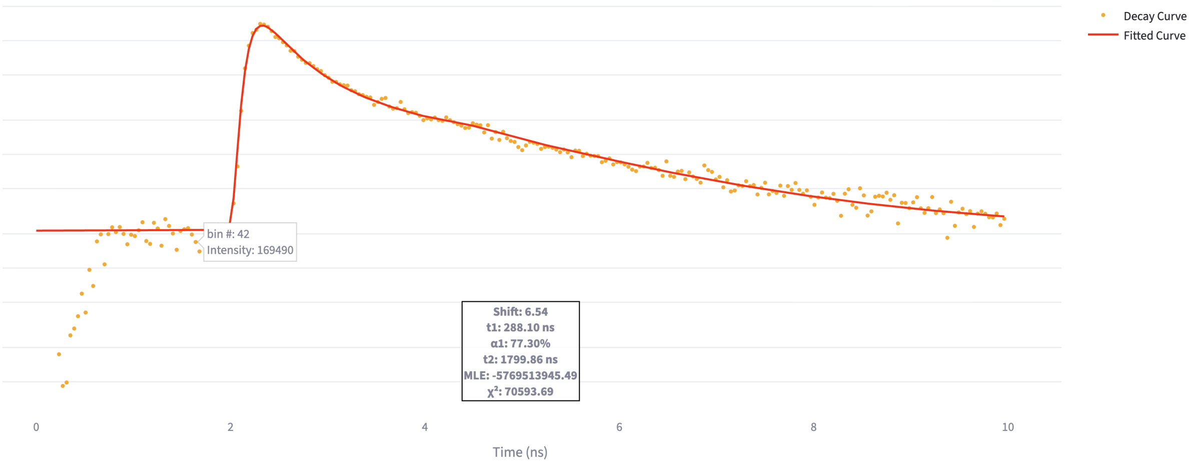

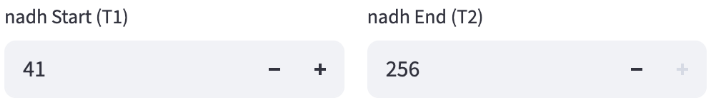

Only the time bins within the T1 and T2 gates are used to calculate the cost metric.

###### 5.2.1.2.3 Metric

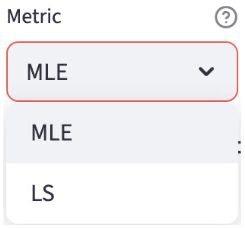

Maximum Likelihood Estimation (`MLE`) estimates the parameters by maximizing the likelihood function, which is the probability of the measured data given the model parameters. A mathematically convenient way to do this is to minimize the negative log-likelihood function:

$$\mathrm{NLL}(\theta; t_s, t_e) = -\sum_{n=0}^{N-1} \mathbf{1}_{[t_s,t_e]}(t_n)\left[y_n \log \hat{y}(t_n) - \hat{y}(t_n)\right].$$

$\mathbf{1}_{[t_s,t_e]}$ is the indicator function that is 1 if $t_n$ is within the time gates $[t_s, t_e]$, and 0 otherwise. $y_n$ is the observed count at time $t_n$, and $\hat{y}(t_n)$ is the model prediction at $t_n$.

Least Squares (`LS`) estimates the parameters by minimizing the sum of the squared differences between the measured curve and the model prediction.

###### 5.2.1.2.4 Fitting Mode

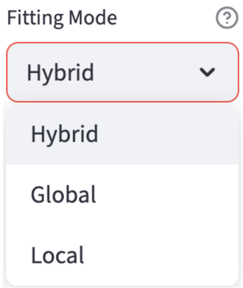

After constructing the objective metric, the fitting mode is used to determine the optimization algorithm. It is a trade-off between the speed and the effort to avoid local minima.

- `Global`: It uses the `differential evolution` algorithm, a derivative-free, population-based but slow global optimizer.
- `Local`: It uses the `leastsq` (least squares with Levenberg-Marquardt) algorithm if the chosen metric is `LS`, otherwise it uses the `nelder` (Nelder-Mead) algorithm.
- `Hybrid`: The most time-consuming option but combines the best of both worlds. It uses the `differential evolution` to find a good initial guess to all the parameters, and then uses the `Local` to drill in.

##### 5.2.2 Fit Free Calibration

Users get to choose between the following two methods to calibrate the IRF shift in the configuration step.

The first step of the calibration is to **subtract an offset value and clip each decay curve to 0 if the time bin is negative**. The offset $C$ is estimated as the mean signal over the final 10% of time bins in each decay ('tail').

Then, depending on the chosen calibration method, the following two methods are available:

###### 5.2.2.1 Shift IRF

It is performed similarly to the fit calibration steps, but in the second step, the shift is not optimized (fitted). It shares the same interface as the fit calibration step, where a scatter plot of the shift values is displayed for each channel, only that the plot is not interactive to show the fit. Instead, it outputs the shift that maximizes the cross-correlation between the IRF and each decay. If `Fix the Shift` is selected, users can specify the shift value that will be applied to all fields of view. Otherwise, the shift value is chosen to be the one that maximizes the cross-correlation between the IRF and the decay.

> **ℹ Note**
>
> If both the `Lifetime fit` and `Lifetime fit free` feature extractors are selected for this channel, the optimized IRF shift derived from the fit calibration step is reused for the `Lifetime fit free` feature extractor.

The signals of the shifted IRF are deconvolved from the offset-subtracted decay curves using `phasor.phasor_divide` from the phasorpy package.

###### 5.2.2.2 Fluorescence lifetime standard

Because the fluorescence lifetime standard is measured in the same system as the decay curves and we know its lifetime, there is no need to account for the IRF shift and the offset. Therefore, after subtracting the offset, `phasorpy`'s `lifetime.phasor_calibrate` function is used to calibrate all the decay curves behind the scenes when calculating the phasor coordinates.

##### 5.2.3 Confirm Calibration

Once users are satisfied with the calibration settings (fitting options and shift values), they can click the `Confirm Calibration` button at the bottom of the page.

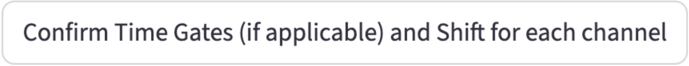

Once confirmed, the left panel prompts users to either apply the calibration settings to the downstream extractors or calibrate again. Users can also save the updated settings (fitting options and shift values) to the metadata file.

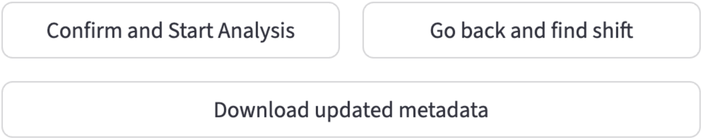

#### 5.3 Feature Extraction

If the selected feature extractors do not require IRF shift calibration, or users have finished the calibration, they can proceed to the feature extraction step by clicking the `Confirm and Start Analysis` button.

Similar to the fov metadata extraction step, FLIM Playground extracts the features and displays the extraction status for each field of view.

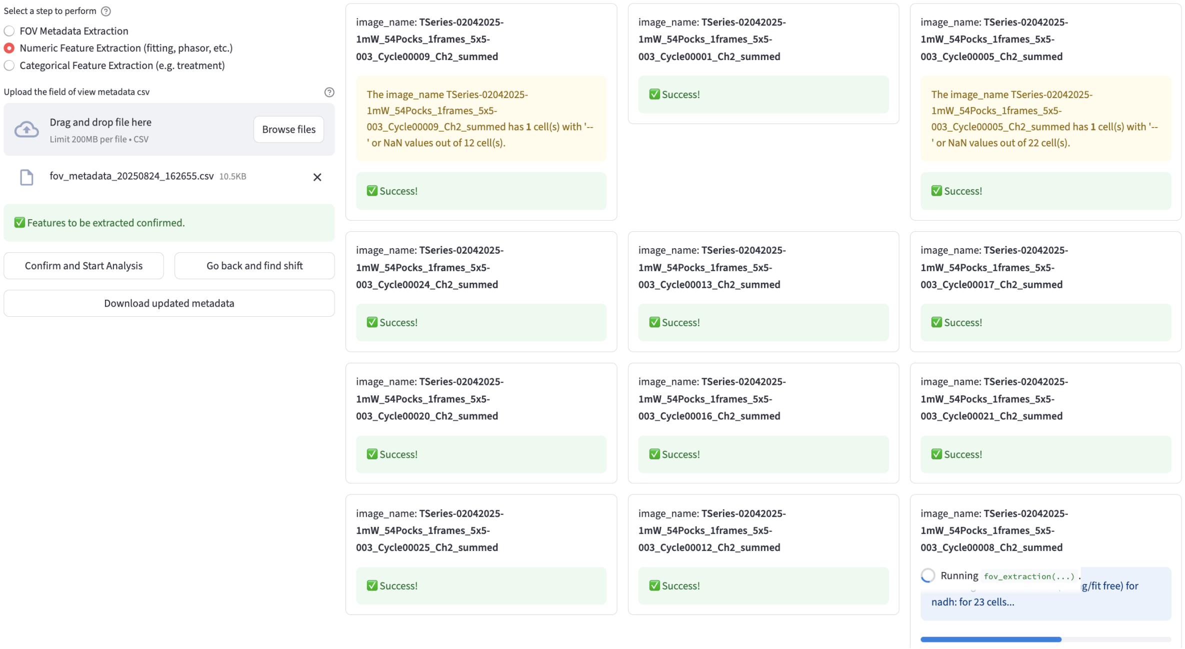

A progress bar is rendered to show the feature extraction progress for each FOV (bottom right).

Warnings are displayed if cells have `NaN` values for some features. They are *not* excluded from the final CSV file.

#### 5.4 Save Results

Once the feature extraction is finished, users can save the results to a CSV file by clicking the `Download` button at the bottom of the page.

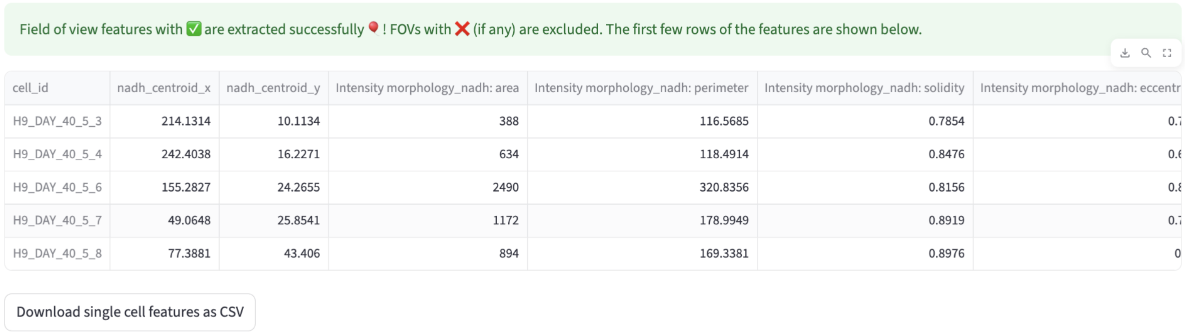

### 6 Lifetime Features (Fit)

Biological samples often contain mixtures of fluorescent species, each with its own characteristic lifetime. By modeling the decay as a sum of exponentials, the analysis can separate and quantify these different contributions. For example, NADH in cells exists in both free (short lifetime, ~0.4 ns) and protein-bound (longer lifetime, ~2–3 ns) states; by fitting the fluorescence decay with a bi-exponential model, one can estimate the fraction of each state, which provides direct insight into cellular metabolism and energy production pathways.

FLIM Playground extracts cell-level lifetime fitting features in this feature extractor, including the fraction of each lifetime component ($\alpha_i$), their lifetimes ($\tau_i$), and mean lifetime ($\tau_{mean}$).

$$I(t) = \sum_{i=1}^{n} A_i \, e^{-t/\tau_i}$$

#### 6.1 Fitting

If the decays have not been pre-fitted, FLIM Playground applies the confirmed fitting options (number of components, time gates, metric, and fitting mode) to the same reconvolution fitting process, as described in the irf shift calibration step, with the only difference being that the IRF shift is no longer a free variable to be optimized. To recap, the reconvolution fitting minimizes the difference between the measured curve and the fitted curve modeled by a multi-exponential model convolved with the IRF, quantified by the cost metric.

Otherwise, FLIM Playground uses the pre-fitted values to calculate the fitting features.

In the final dataset, fitting features are prefixed by the combination of the feature extractor name (i.e. `Lifetime fit`) and the channel name, allowing Data Analysis to group the features. For example, `Lifetime fit_nadh: a1` means the fraction of the first lifetime component for the NAD(P)H channel.

##### 6.1.1 Preprocessing

It sums up all the pixel decays belonging to the same cell ROI labeled by the ROI mask as one decay curve[9]. The ROI summing reduces variability and bias and allows for short integration times at acquisition in exchange for sub-cellular resolution (i.e., pixel-level). Users do not need to specify the bin factor (each pixel sums up the surrounding pixels' decay curves) to account for insufficient photon counts. Also, this summing lets a single-threaded CPU application finish in a reasonable amount of time.

It also shifts the IRF using the shift values from the IRF shift calibration step. To shift an IRF, FLIM Playground upsamples the IRF 10 times using linear interpolation to fill the gaps. Then it shifts the IRF by the shift values $\times 10$ and downsamples the IRF back to the original size.

##### 6.1.2 Fraction of components

In addition to the absolute amplitudes of each component directly from the fitting result, FLIM Playground calculates the fraction of each component, $\alpha_i$, as the amplitude of the component divided by the sum of all amplitudes. This normalization allows lifetime to be independent of the absolute intensity of the signal.

##### 6.1.3 Mean lifetime

The mean lifetime, $\tau_{mean}$, is calculated as the weighted average of the lifetimes of all components, where the weights are the fractions of each component.

$$\tau_{mean} = \sum_{i=1}^{n} \alpha_i \tau_i$$

#### 6.2 Pixel-prefitted

Currently, FLIM Playground supports pixel-prefitted lifetime features from SPCImage. The pixel-level lifetime fitting features are expected to be stored in 2D arrays in spatial dimensions, with each row and column having the value of a lifetime feature outputted from SPCImage. SPCImage assigns `0` for pixels that are not fitted (e.g. thresholded out). To avoid them biasing the results, FLIM Playground uses `np.ma.masked_array` to create a masked array and disregards them when calculating the averages using `np.ma.average`.

Then it uses the ROI mask to calculate the cell-level lifetime features by averaging the pixel-level features within each ROI.

### 7 Lifetime Features (Fit-Free)

Phasor analysis provides a fast, model-free view of lifetimes by transforming fluorescence decays into phasor coordinates through a Fourier transform. This approach is considered "fit-free" because it does not require iterative curve fitting or predefined decay models—each decay is mapped directly to a point on the phasor plot. It has the nice property that mono-exponential decays fall on the universal semicircle while mixtures lie inside as linear combinations, so clusters reveal distinct species and their fractional contributions. FLIM Playground currently extracts cell-level phasor features: `g`, `s`, `Tau_phase`, `Tau_m`, `g_2nd` (2nd harmonic), and `s_2nd`.

Fit-free features in the final dataset are prefixed by the combination of the feature extractor name (i.e. `Lifetime fit free`) and the channel name, allowing Data Analysis to group the features. For example, `Lifetime fit free_nadh: G` means the $g$ coordinate for this cell in the NAD(P)H channel.

#### 7.1 Preprocessing

It sums up all the pixel decays belonging to the same cell ROI labeled by the ROI mask as one decay curve[9]. The phasor features are calculated from the summed decay curve.

If the chosen calibration method is IRF shift calibration, FLIM Playground shifts the IRF using the shift values and subtracts the estimated offset as described in the IRF shift calibration step.

#### 7.2 Phasor features

##### 7.2.1 Step 1: raw phasor coordinates

The reference implementation from phasorpy assumes the decay curve comes from one period and time bins are evenly spaced (untruncated). So, their calculation is frequency ($f$) and time ($T$) independent. However, there are cases when the decay curve is truncated due to the deadtime of the system. For example, when the laser frequency is $f = 0.08$ GHz, the theoretical period is $T = 12.5$ ns, but in practice the time window is truncated to $T = 10$ ns. FLIM Playground applies a custom implementation to the truncated decay curve. (The `sample_phase` option can be used to account for the truncation but it does not handle the second or higher harmonics)

###### 7.2.1.1 Untruncated decay curve

It uses the phasor_from_signal function from the phasorpy package to calculate the uncalibrated phasor coordinates g_raw and s_raw, g_raw_2nd and s_raw_2nd, the first and second harmonics' real and imaginary parts, from the cell decay curve.

Either g_irf, s_irf, g_irf_2nd, and s_irf_2nd from the IRF, or ref_mean, ref_real, ref_imag from the fluorescence lifetime standard, depending on the chosen calibration method, are calculated using the same function.

Mathematically, the raw phasor coordinates are calculated as (copied from the phasorpy documentation):

$$F_{\mathrm{DC}} = \frac{1}{K}\sum_{k=0}^{K-1} F_k, \quad g_{\mathrm{raw}} = \frac{1}{F_{\mathrm{DC}}}\frac{1}{K}\sum_{k=0}^{K-1} F_k \cos\left(2\pi h\frac{k}{K}\right), \quad s_{\mathrm{raw}} = \frac{1}{F_{\mathrm{DC}}}\frac{1}{K}\sum_{k=0}^{K-1} F_k \sin\left(2\pi h\frac{k}{K}\right)$$

where $K$ is the number of time bins, $F_k$ is the decay curve, and $h$ is the harmonic number.

###### 7.2.1.2 Truncated decay curve

The phasor coordinates of the IRF and the decay curves of both harmonics are calculated using the time axis $t_k = k\,(T/K)\ \ (k = 0, \ldots, K-1)$ and the angular frequency $\omega = 2\pi\, h f$:

$$g_{\mathrm{raw}} = \frac{\sum_{k=0}^{K-1} F_k \cos(\omega\, t_k)}{\sum_{k=0}^{K-1} F_k}, \quad s_{\mathrm{raw}} = \frac{\sum_{k=0}^{K-1} F_k \sin(\omega\, t_k)}{\sum_{k=0}^{K-1} F_k}$$

This formula is equivalent to the untruncated formula when $fT = 1$.

For the fluorescence lifetime standard, the phasor coordinates are calculated using phasorpy's phasor_from_signal and the sample_phase option is set to $\omega t_k$.

##### 7.2.2 Step 2: calibrated phasor coordinates

To get the calibrated phasor coordinates g, s, g_2nd, and s_2nd, FLIM Playground uses either

```
from phasorpy import phasor
g, s = phasor.phasor_divide(g_raw, s_raw, g_irf, s_irf)
g_2nd, s_2nd = phasor.phasor_divide(g_raw_2nd, s_raw_2nd, g_irf_2nd, s_irf_2nd)
```

or

```
from phasorpy import lifetime
g, s = lifetime.phasor_calibrate(g_raw, s_raw, ref_mean, ref_real, ref_imag, frequency=laser_
g_2nd, s_2nd = lifetime.phasor_calibrate(g_raw_2nd, s_raw_2nd, ref_mean, ref_real, ref_imag,
```

The `laser_rate` is specified during the fov metadata extraction and the configuration interface.

##### 7.2.3 Step 3: phasor-derived lifetime

`Tau_phase` and `Tau_m` are calculated using the first harmonic's phasor coordinates g and `s`.

```
import numpy as np
w = 2 * np.pi * laser_rate
phi = np.arctan2(s, g)
m = np.sqrt(g**2 + s**2)
tau_phase = 1/w * np.tan(phi)
tau_m = 1/w * np.sqrt(1/m**2 - 1)
```

$$\phi = \text{atan2}(s, g), m = \sqrt{g^2 + s^2}, \tau_\phi = \frac{\tan\phi}{\omega}, \tau_m = \frac{1}{\omega}\sqrt{\frac{1}{m^2} - 1}$$

### 8 Morphological Features

FLIM Playground extracts single-cell morphological features based on the cell region of interest (ROI) mask of each *channel*. By inputting the ROI mask to `skimage.measure.regionprops`, the following features are extracted:

- `area`: the number of pixels in the ROI.
- `perimeter`: length of the ROI boundary in pixels.
- `solidity`: Solidity $= \frac{\text{\# ROI pixels}}{\text{\# convex hull pixels}}$, where the convex hull is the smallest convex polygon that contains the ROI. A value close to 1 means the shape has few concavities (more solid), while lower values indicate more irregular boundaries.
- `eccentricity`: equals the eccentricity of the ellipse that has the same second moments as the region. It is given by the ratio of the focal distance (the distance between the foci) to the length of the major axis: $e = \frac{c}{a}, \quad e \in [0, 1)$, where $a$ is the semi-major axis length, $b$ is the semi-minor axis length, and $c = \sqrt{a^2 - b^2}$ is the focal distance. When $e = 0$, the ellipse reduces to a circle. It quantifies how elongated the ROI is, with 0 being a perfect circle and 1 being a line.
- `major_axis_length`: the length of the major axis of the ellipse that has the same second moments as the region: $2a$. It represents the longest dimension of the ellipse that approximates the region's shape, essentially describing its maximum elongation.
- `minor_axis_length`: the length of the minor axis of the ellipse that has the same second moments as the region: $2b$. It represents the shortest dimension of that ellipse, capturing the region's minimum elongation.
- `circularity`: describes how close the ROI is to a perfect circle. It is defined as circularity $= \frac{4\pi A_{\text{ROI}}}{P_{\text{ROI}}^2}, \quad$ circularity $\in (0, 1]$, where $A_{\text{ROI}}$ is the area of the region and $P_{\text{ROI}}$ is its perimeter. A value of 1 corresponds to a perfect circle.

Feature names in the final dataset are prefixed by the combination of the feature extractor name (i.e. `Intensity morphology`) and the channel name, allowing Data Analysis to group the features. For example, `Intensity morphology_nadh: area` means the area of each cell in the NAD(P)H channel.

### 9 Textural Features

FLIM Playground extracts single-cell texture features based on the intensity image and the ROI mask from each acquisition *channel*. For an example application, refer to this paper that used CellProfiler to extract the texture features.

The first step is to create a single-cell intensity image for each cell based on the ROI mask and the intensity image. Then a list of texture features (the list may be extended in the future) is calculated:

#### 9.1 Granularity

$Granularity_n$ is the percentage of intensity contributed by bright objects of $n$ pixels in radius. It is computed by performing a morphological opening with a disk of radius n on the cell intensity image and measuring the intensity lost as a percentage of the total intensity of the original cell ROI.

$n = 1, 3, 5, 7, 9$ are computed.

Semantically, higher granularity at smaller scales implies higher fragmentation.

#### 9.2 Radial Distribution

To calculate the radial distribution, each cell intensity image is partitioned into $4$ concentric rings, and the intensity fraction over the total intensity for each ring is calculated. For example, a higher fraction in ring 1 implies that signals are more concentrated at the center of the cell, whereas a higher fraction in ring 4 implies that signals are more concentrated at the cell edge.

#### 9.3 Mass Displacement

Mass displacement is the Euclidean distance (in pixels) between the intensity-weighted centroid of the cell ROI and the geometric centroid of the same ROI.

#### 9.4 Intensity Sum

Intensity sum is defined as the sum of pixel intensities within the cell intensity image.

In the final dataset, feature names are prefixed by the combination of the feature extractor name (i.e. `Intensity texture`) and the channel name, allowing Data Analysis to group the features. For example, `Intensity texture_nadh: granularity_1` means the percentage of intensity removed when bright objects of 1-pixel diameter are removed in the NAD(P)H channel.

### 10 Categorical Feature Extraction

Based on the datasets extracted in the numerical feature extraction step, FLIM Playground:

- merges the datasets into a single dataset
- assigns categorical features to each cell in the merged dataset

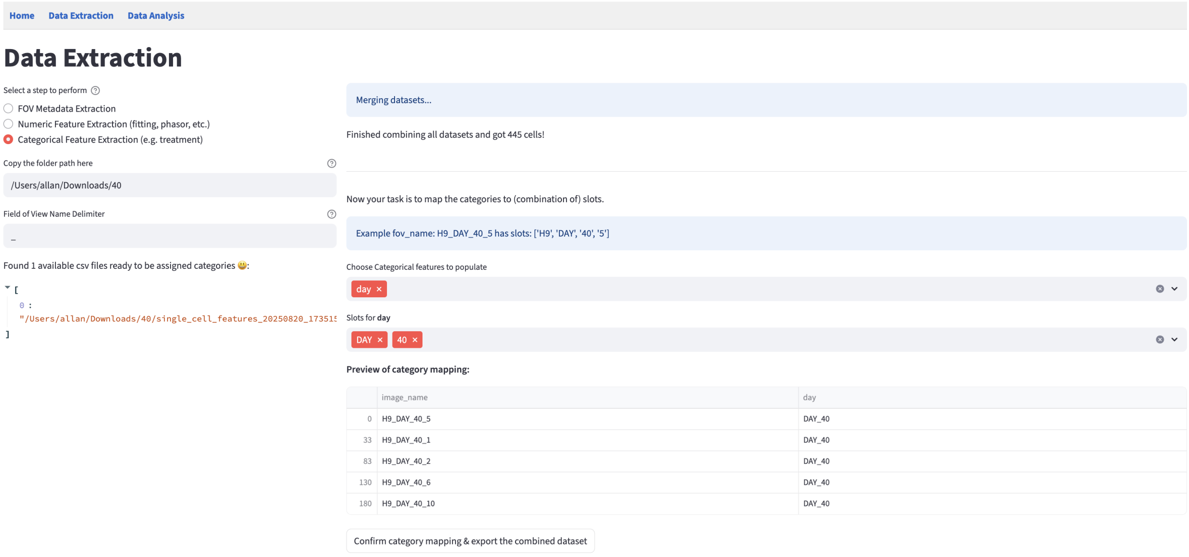

#### 10.1 Merging Datasets

It scans the specified folder path and finds all CSV files. It uses the cell identifier column to check for duplicates both within each dataset and across all datasets.

> **! Important**
>
> **The FOV identifier uniquely identifies a field of view and is assumed to contain all information necessary to extract the categorical features (e.g., treatment, time point, etc.), by including segments separated by a certain delimiter (e.g., _). Categorical features are assigned from individual segments or their combinations.** Example: The FOV `Panc1_Cyanide_dish1`

denotes that it comes from the Panc1 cell line, under the Cyanide treatment, in the 1st dish.

Therefore, it checks if all field-of-view names include the same number of segments to determine the eligibility of categorical feature assignment.

Field of View Name Delimiter

_

The image_name column in /Users/allan/Downloads/DATA/donor6.csv has different number of parts. For example, check fov_name: HC220113_1n.

The image_name column in /Users/allan/Downloads/DATA/donor5.csv has different number of parts. For example, check fov_name: HC130319_1n.

Found 7 available csv files ready to be assigned categories 😄:

```
[
  0 : "/Users/allan/Downloads/DATA/donor8.csv"
  1 : "/Users/allan/Downloads/DATA/donor9.csv"
  2 : "/Users/allan/Downloads/DATA/donor1.csv"
  3 : "/Users/allan/Downloads/DATA/donor2.csv"
  4 : "/Users/allan/Downloads/DATA/donor3.csv"
  5 : "/Users/allan/Downloads/DATA/donor7.csv"
  6 : "/Users/allan/Downloads/DATA/donor4.csv"
]
```

It merges the datasets that pass the checks by finding common columns and vertically concatenating them.

Merging datasets...

Merging dataset 2 (/Users/allan/Downloads/DATA/donor9.csv) into the combined dataset.

Merging dataset 3 (/Users/allan/Downloads/DATA/donor1.csv) into the combined dataset.

Merging dataset 4 (/Users/allan/Downloads/DATA/donor2.csv) into the combined dataset.

Merging dataset 5 (/Users/allan/Downloads/DATA/donor3.csv) into the combined dataset.

Merging dataset 6 (/Users/allan/Downloads/DATA/donor7.csv) into the combined dataset.

Merging dataset 7 (/Users/allan/Downloads/DATA/donor4.csv) into the combined dataset.

Finished combining all datasets and got 3037 cells!

#### 10.2 Assigning Categorical Features

Available categorical features to be assigned are drawn from the user-specified configuration.

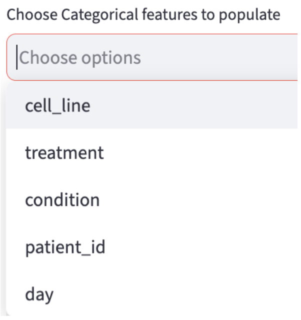

Based on the assumptions, it breaks down the FOV identifier into individual fields and facilitates the assignment.

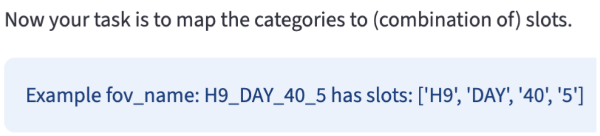

A live preview of the assignment is displayed.

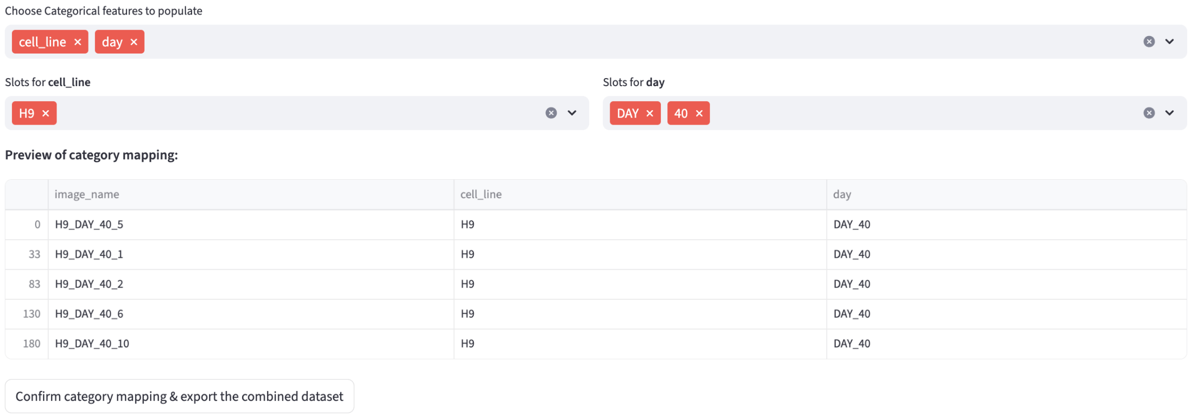

And the merged dataset with categorical features is available for download.

### Part II

### Data Analysis

### 11 Overview

We are not done yet 😱! FLIM Playground also provides a suite of methods to analyze and visualize the extracted data (either using the Data Extraction section or in users' own way). They are designed to be interactive and frictionless ⚡ so that users can perform hassle-free exploration of their data with, hopefully, fun 😎. Inspired by the same data categorization in Data Extraction, the Data Analysis section designs an architectural blueprint that all in-house analysis methods build on. It is modularized so that new methods and features can be added in the future easily.

#### 11.1 Methods

- Depending on the number of numerical features in the analysis, the methods are categorized into 3 groups:
  - Univariate Analysis
    - Feature Comparison
    - Feature Histogram
    - Field of View Comparison
  - Bivariate Analysis
    - 2D Feature Distribution
    - Phasor Analysis
  - Multivariate Analysis
    - Dimension Reduction
    - Classification

All the methods share a set of interactive widgets to support the general workflow. They also have their own method-specific widgets, the descriptions of which are provided in the corresponding method pages.

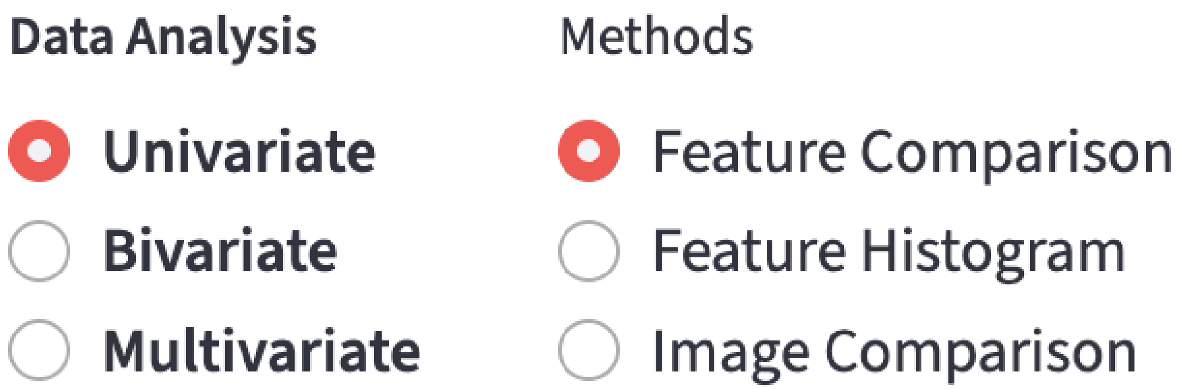

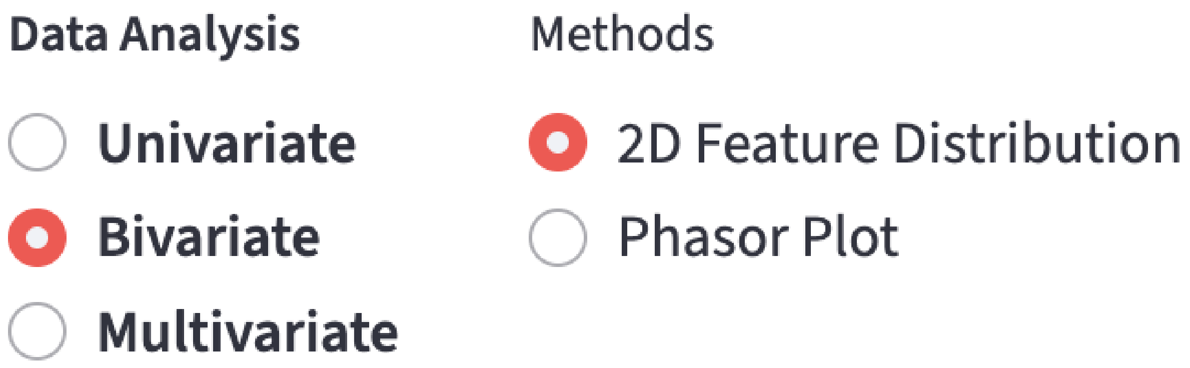

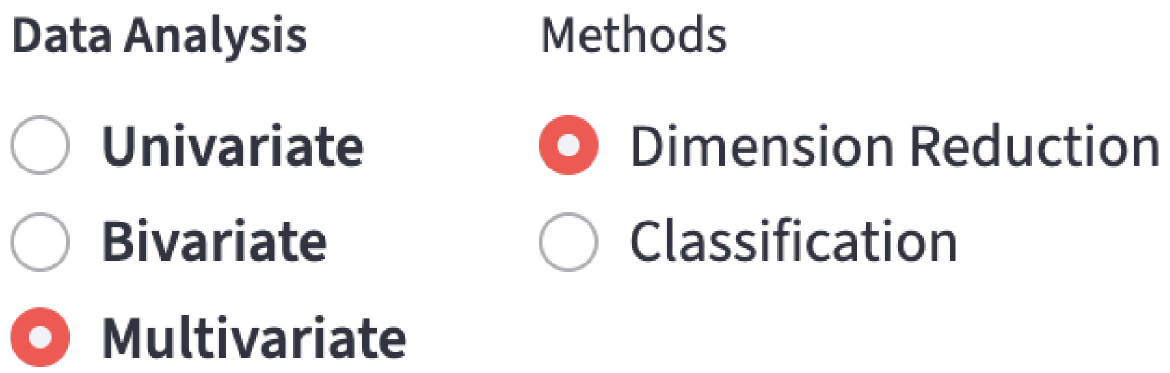

#### 11.2 Input

Users can upload the dataset output from the Data Extraction directly or *their own datasets* in **CSV** (Comma Separated Values) format after finishing the interactive configuration setup.

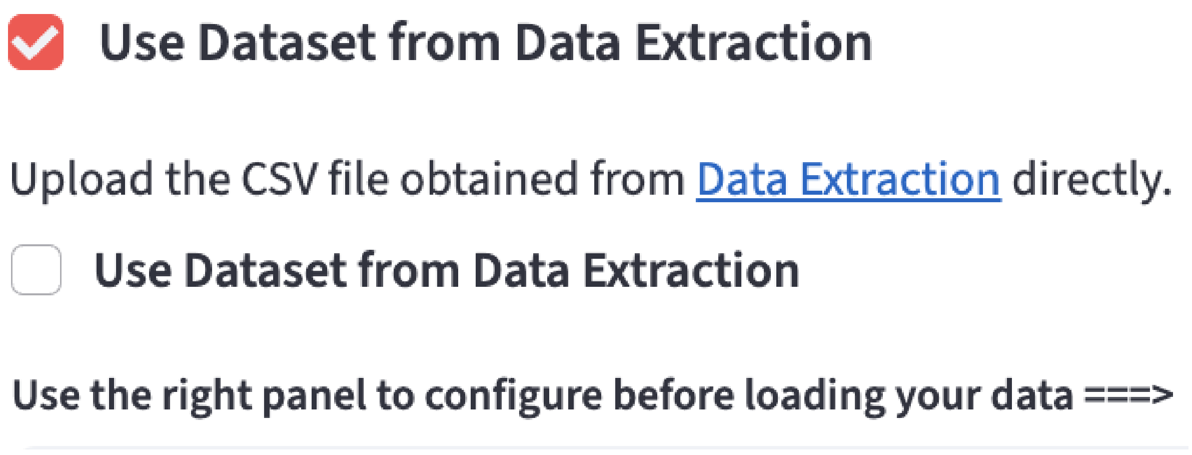

##### 11.2.1 Requirements

> **Important**
>
> In either case, following the categorization, the dataset should have:
>
> - a column that uniquely identifies each row
> - an (optional) field of view identifier column
>
> - a set of numerical features
> - zero or more categorical features (e.g. treatment, day, patient id, etc.)

The unique row id and field of view id are used to help users identify the row (e.g. a single cell) and field of view (e.g. a single image) of data of interest through the built-in hover-based interaction. Numerical and categorical features are used to render shared widgets. Internally, FLIM Playground will check whether the dataset fulfills the requirements and output meaningful warning or error messages.

> **Note**
>
> Warning messages will not prevent the analysis but error messages will.

##### 11.2.2 Warning Messages

- `Empty columns`: will be dropped.
- `Duplicated columns`: Only the first occurrence of the duplicated column will be kept. Other occurrences will be dropped.

- `Duplicated rows based on the unique row id`: Only the first occurrence of the duplicated row will be kept. Other occurrences will be dropped.
- `Columns with NaN values`: won't be dropped, just a warning message.
  - The analysis will be performed on the rows that are not NaN in the selected numerical features.

##### 11.2.3 Error Messages

- Missing a column that uniquely identifies each row
- Cannot identify any numerical feature column
  - It uses `pd.api.types.is_numeric_dtype` to check if a column is numerical.

#### 11.3 Shared Interactive Widgets

A list of shared widgets is provided to support the general workflow in analysis.

##### 11.3.1 General Workflow

- select numerical feature(s) on the left
- subset the data to find data of interest on the top
- use visual channels widgets to look at the data in different ways
- change the plot style
- hover to find data points of interest

The visualization and analysis results are updated in real time to reflect users' interactions with any of the above widgets.

##### 11.3.2 Numerical Feature Widgets

The Data Extraction recognizes numerical features using `pd.api.types.is_numeric_dtype` internally. Then it groups numerical features based on the feature extractor type and the channel it belongs to. For user-provided datasets, the numerical features are grouped based on the user-specified configuration. Ungrouped numerical features are put into the `Uncategorized Features` group. One selection widget is rendered for each numerical feature group.

> **Tip**
>
> If you cannot find a certain numerical feature under any numerical widgets including `Uncategorized Features`, please inspect the dataset (e.g., using Excel filters) and look for non-numeric values in that column. One non-numeric value (e.g., "–") will prevent it from being recognized.

###### 11.3.2.1 Univariate Analysis

One single-select widget is rendered per group; choosing a feature in any group clears the selections and resets the others to `Select`.

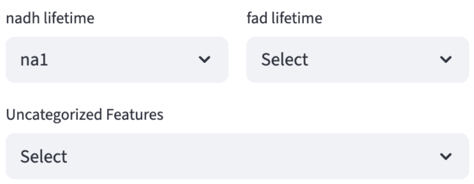

###### 11.3.2.2 Bivariate Analysis

Two sets of selection widgets are rendered to allow maximum flexibility (the two features can be from the same or different feature groups). Each set of selection widgets behaves like the selection widgets in the univariate analysis. The first selected feature will be hidden in the second set of selection widgets.

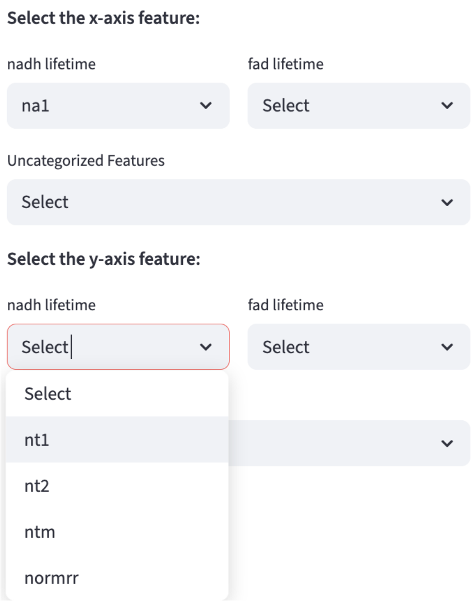

###### 11.3.2.3 Multivariate Analysis

One set of selection widgets, each can select multiple features from a feature group, is rendered. A special value `All` is introduced so that users can conveniently select all features under the feature group. If users select `All`, all the other options will be cleared, and vice versa.

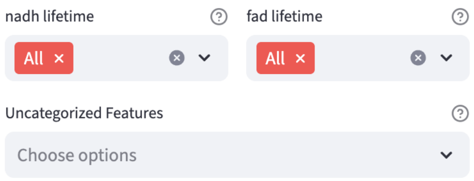

##### 11.3.3 Categorical Feature Widgets

FLIM Playground recognizes the categorical features in the uploaded dataset based on the user-specified configuration if the dataset is not extracted by Data Extraction. Otherwise, it recognizes categorical features specified in the Data Extraction configuration.

###### 11.3.3.1 Filter Widgets

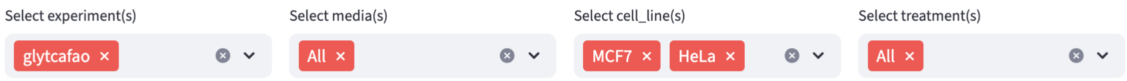

For complex datasets that are collected over multiple days, experiments, treatments, etc., it is useful to filter the data to focus on a subset of the data (data of interest). One filter widget is rendered for each categorical feature so that users have the flexibility to filter the data based on combinations of categorical features. `All` is a special option that include all categories of the selected categorical feature. Once it is selected, all the other options are cleared, and vice versa.

###### 11.3.3.2 Visual Channels Widgets

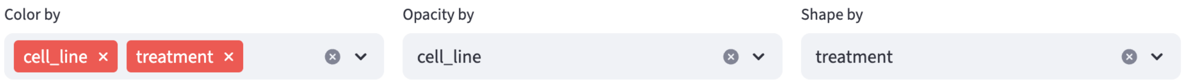

Human vision is wired for rapid, parallel pattern and trend detection; good visual encodings (how to map data to visual elements such as color, shape, opacity, etc.) harness this to surface insights that raw numbers or text obscure[12]. FLIM Playground provides `color`, `opacity`, and `shape` channels for visualizations that support them:

###### 11.3.3.2.1 Color by

Color is supported in all methods except for Classification. In `Color by`, users can select multiple categorical features, and each unique combination of available categories (determined by the filters in the Filter Widgets) in selected features is assigned a distinct color. Groups created by `Color by` show up in the x-axis (if multiple features, categories from each feature are delimited by `::`).

> **Note 1: How to order the groups**
>
> The order of the groups in x-axis and in the legend is determined by:
>
> - the order of the selected features: categories of the first feature appear before those of the second, and so on and so forth. For example, the two treatments of `Panc1` are together, because it is sorted first on the cell lines, then on treatments. If you want the treatments to be together, in `Color by` you can select `treatment` then `cell_line`.
> - *Numeric-alphabetical sort*: within each feature, the category order is determined by the number inside (e.g. `1` in `Panc1`) first, then the alphabetical order of the category name. Therefore, although `P` is after `M` alphabetically, the `1` is before `7` numerically, making `Panc1` appear before `MCF7`. This may be helpful when you have data from different days or hours. The default string sort will put `Day 100` before `Day 30`, and this is not what we want.

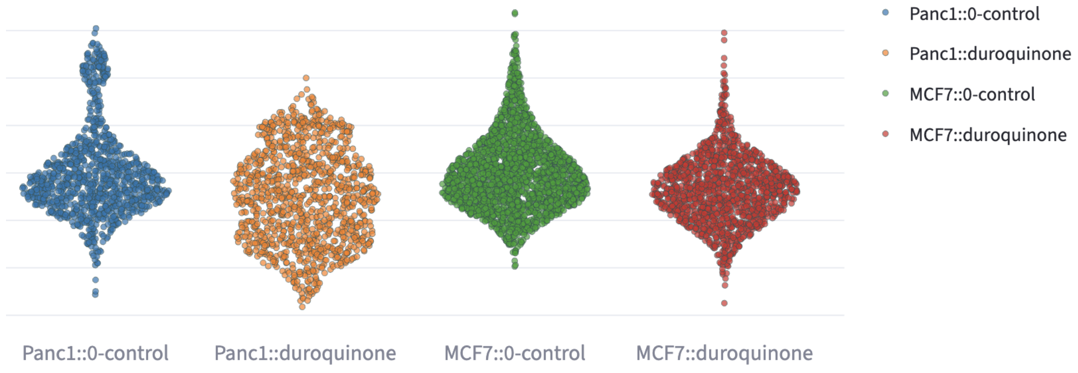

###### 11.3.3.2.2 Opacity and Shape by

- Opacity and shape are supported in all point-based visualizations (e.g. Feature Comparison, 2D Feature Distribution, Phasor Analysis, Dimension Reduction). In `Opacity by` and `Shape by`, users can select one categorical feature, and each unique category is assigned a distinct opacity or shape. The order of the shape and opacity is also sorted numeric-alphabetically.

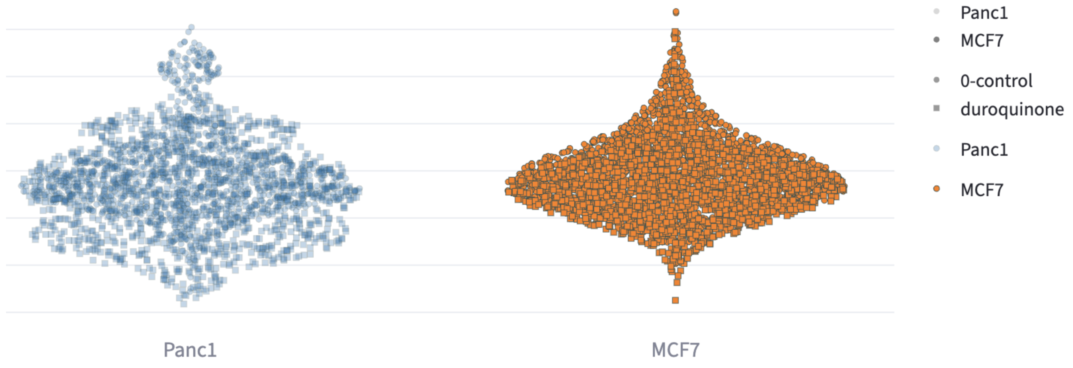

##### 11.3.4 Unique ID Hover

- In all point-based visualizations, when hovering over a point, the unique id of the point will be shown.

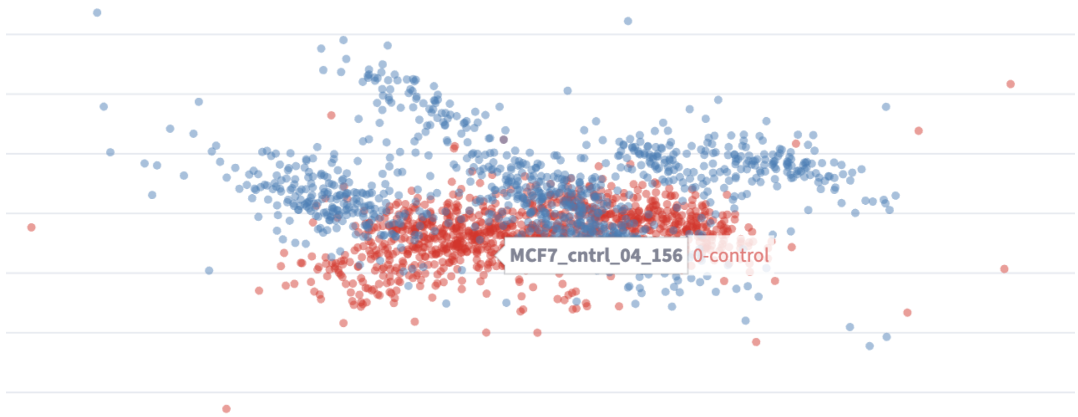

##### 11.3.5 Plotting Configuration Widgets

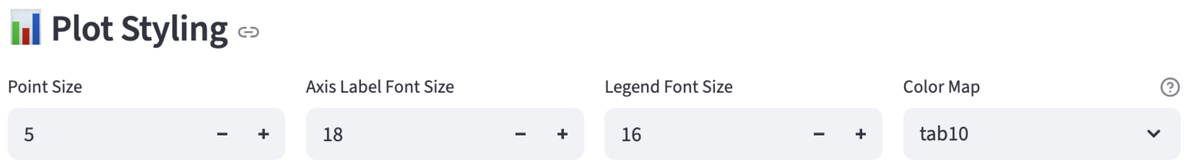

Users can interactively configure the following plot parameters:

- point size (if point-based)
- axis label size

- legend size
- the colormap used to color different groups
  - `colorblind`, `tab10`, `tab20`, `Set1`, `Set2`, `Set3`, `Pastel1`, `Pastel2`, `Accent`, `viridis`, `plasma`, `inferno`, `magma`, `cividis`

### 12 Configuration

FLIM Playground provides an interactive configuration workflow to help users prepare their datasets that are not extracted by Data Extraction so that they fulfill the requirements. The following sections use the iris dataset as an example: the `flower_id` is added to uniquely identify each row (i.e. a single flower). The other columns are: a categorical feature `species`, and a set of numerical features `sepal_length`, `sepal_width`, `petal_length`, and `petal_width`.

Specifically, the steps are:

- On the left side of the screen, uncheck `Use Dataset from Data Extraction`.

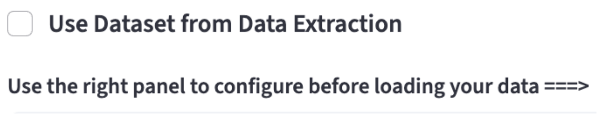

#### 12.1 Identifiers Config

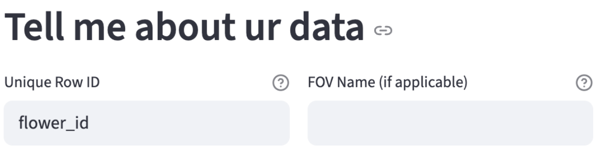

- The dataset needs to have a unique identifier column to identify each row. In this case, `flower_id` is the unique identifier column. This dataset does not have a field of view column, so we leave it blank.

#### 12.2 Categorical Feature Config

- Add categorical columns you want FLIM Playground to recognize here. Those columns are converted to strings internally.

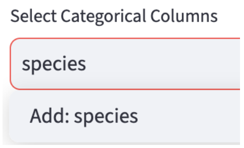

The default categorical feature list does not include `species`, so we add it manually.

#### 12.3 Numerical Feature Config

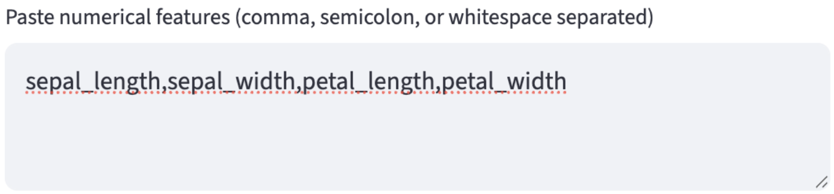

- Users can copy the numerical features from Excel or any text editor that opens their CSV files. The features can be separated by `,`, `;`, or whitespaces including newlines.
- After hitting the `ctrl/cmd + enter` key, the numerical features are parsed and added to `Available Features`. Users can add/delete groups and drag features from the `Available Features` to the groups they've just created. Any ungrouped features are added to a group called `Uncategorized Features`.

##### 12.3.1 Example

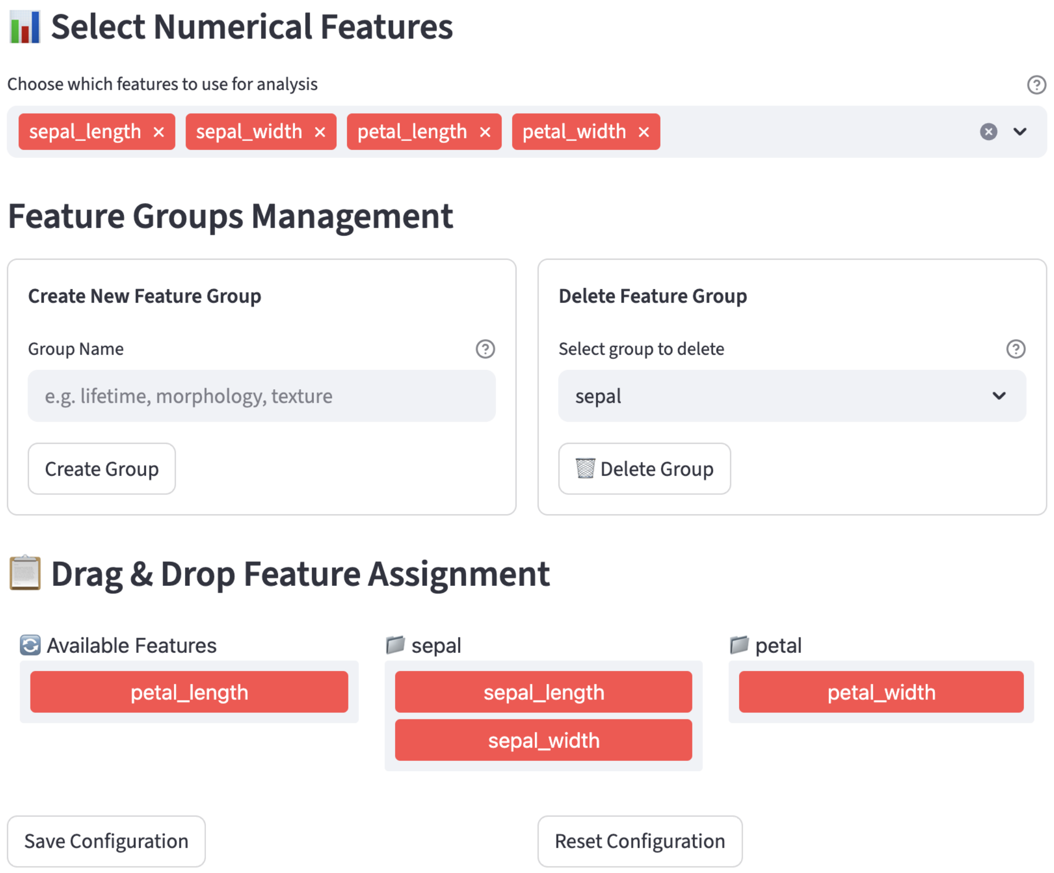

#### 12.4 Save

- After users finish the configuration and click the `Save Configuration` button, FLIM Playground saves the configuration for processing current and future datasets.

#### 12.5 Reset

Users can reset the configuration to the default settings.

### Part III

### Univariate Analysis

### 13 Feature Comparison

This method allows users to compare the distributions of a single numerical feature across groups. It can help users spot overarching trends between distributions, identify within-distribution shape-related characteristics such as skewness or bimodality that might signal sub-populations, and diagnose potential outliers.

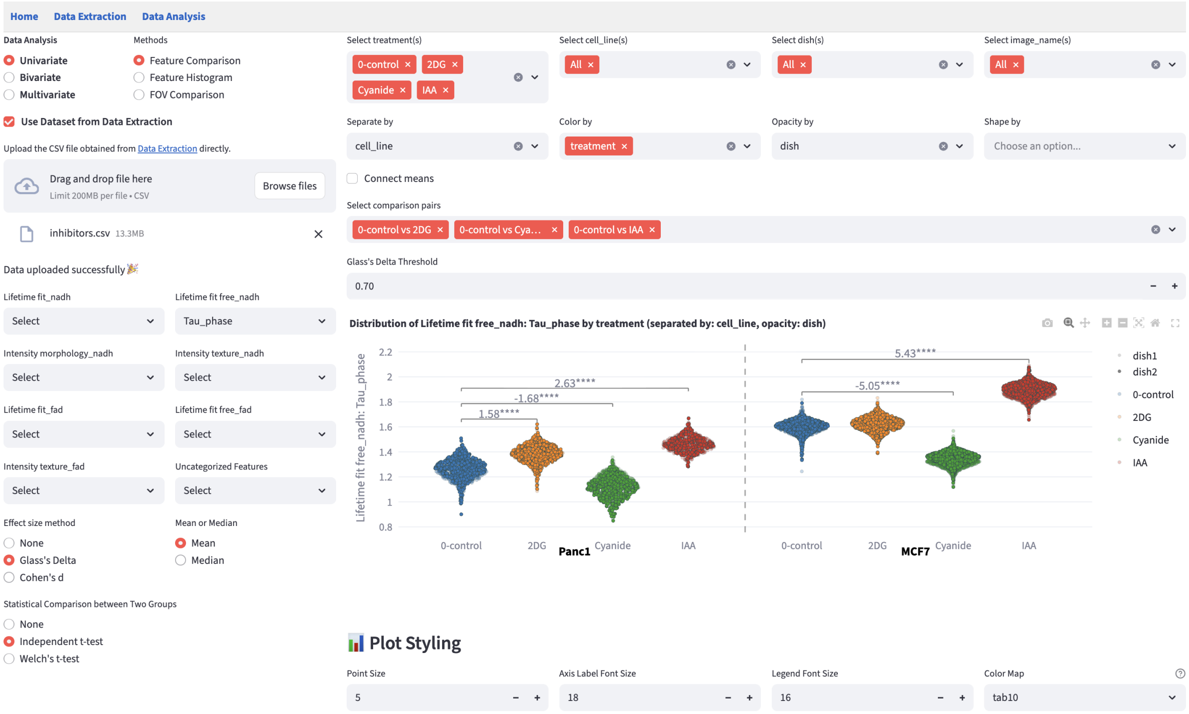

#### 13.1 Shared Interface Components

- On the left, users can use the selection widgets to select the numerical feature to be compared. Exactly one feature can be selected from all feature groups.
- On the top right, users can use the filters to subset the data to find the groups of interest.
- Below the filters, users can apply the visual channels widgets that allow users to `Color by`, `Opacity by`, and `Shape by` categorical features.
- On the bottom right, users can change the plot style using the plot styling widgets.

#### 13.2 Separate by

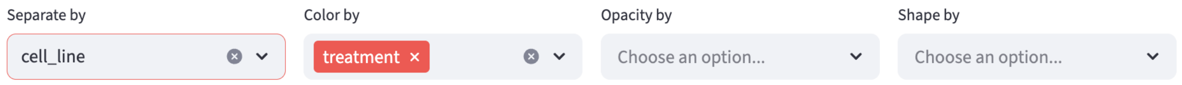

`Separate by` takes effect before `Color by`. `Separate by` divides the plot into sections separated by a dashed line, with each section labeled by one of the available categories of the selected categorical feature after filtering. The selected feature of `Separate by` is automatically removed from available features in `Color by`. The section order is determined by the order of the categories which are sorted numeric-alphabetically.

Then, visual channels (color, shape, opacity) and effect size annotation are applied within each section. For example, the visualization below was separated by `cell_line`, with each section labeled by the bold text of a cell line. Within each section, the x-axis rendered the `Color by` categories (`treatments`) so colors were consistent across sections. Effect size calculation and annotation (only effect size $> 0.5$ is shown) were performed within each section.

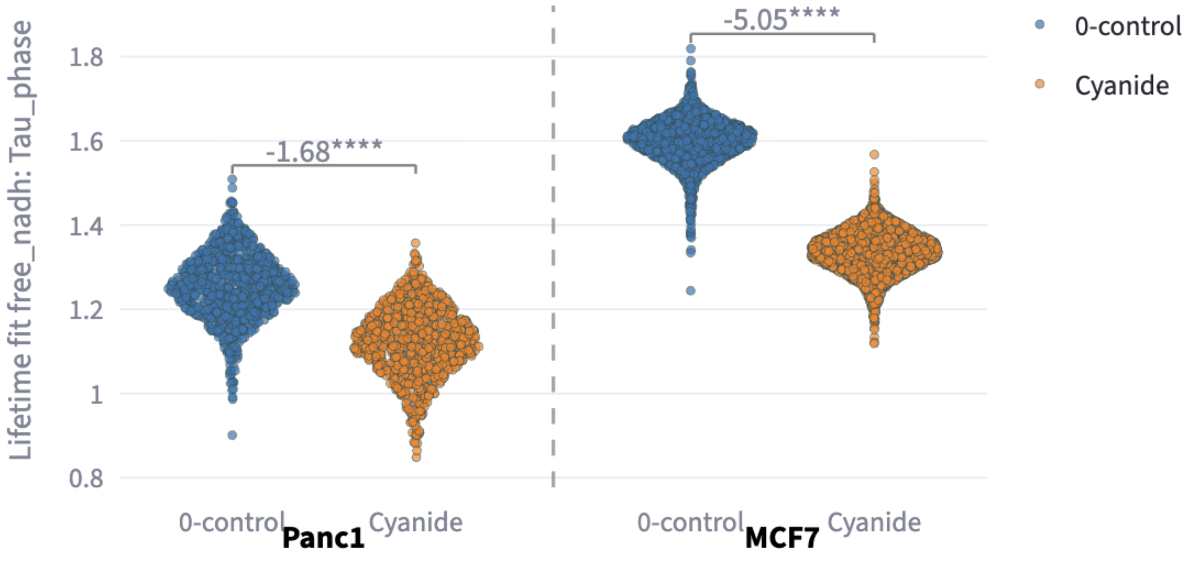

If `Separate by` is left blank and both `cell_line` and `treatment` are selected in `Color by`, then colors are assigned to each cell line and treatment combination, and effect sizes are calculated across all combinations of group pairs (only pairs with effect size $> 0.5$ are shown).

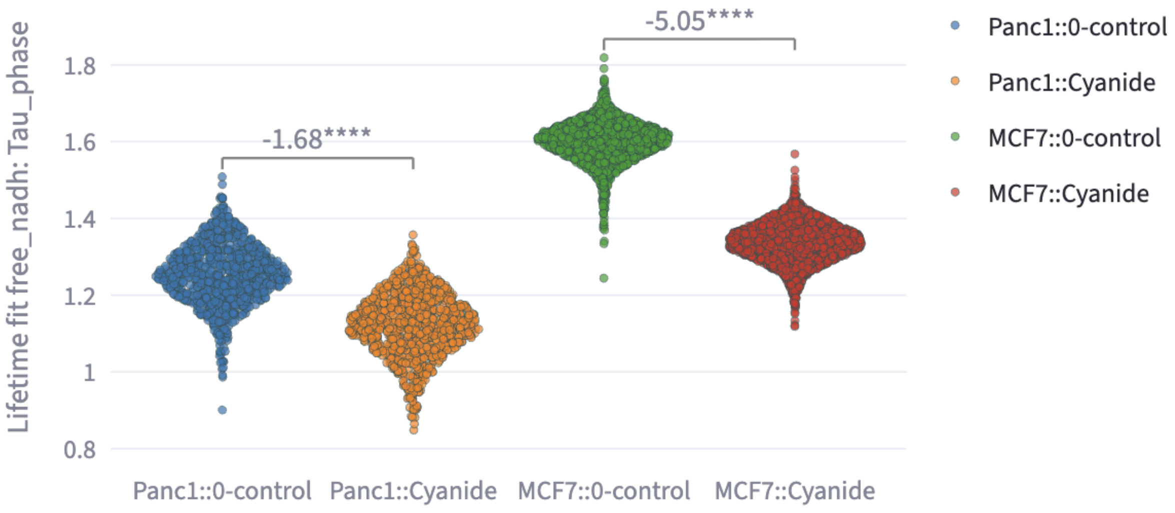

#### 13.3 Comparison Widget

Statistical tests and effect size calculations are performed on the selected comparison pairs. FLIM Playground uses the groups on the x-axis populated by Color by to create comparison groups, starting from the leftmost group. For example, if there are 5 groups, then $\binom{5}{2} = 10$ comparisons will be created. The number of groups can get combinatorially large and many of the comparisons are not meaningful. Therefore, two widgets are provided to filter the comparison groups:

- A selection widget that shows all possible comparisons by default and users can deselect groups they do not need.

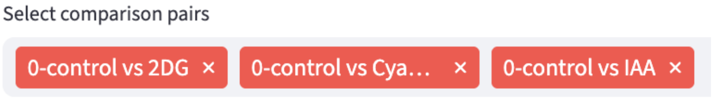

- A threshold by effect size widget that filters out comparisons below the threshold.

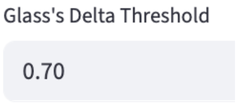

#### 13.4 Effect Size

Complementary to statistical tests, which address "is there a difference?", effect size metrics answer "how large is the difference?", enabling cross-study comparison. FLIM Playground offers two effect size calculation methods, Glass's Delta and Cohen's d, each can be in mean or median form.

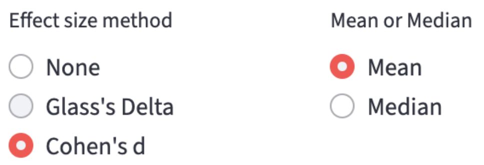

##### 13.4.1 Glass's Delta

Mean ($\bar{X}$) form:

$$\Delta_G = \frac{\bar{X}_{\text{treatment}} - \bar{X}_{\text{control}}}{s_{\text{control}}}, \qquad s_{\text{control}} = \sqrt{\frac{1}{n_c - 1}\sum_{i=1}^{n_c}(X_{control_i} - \bar{X}_{\text{control}})^2}$$

Median ($\tilde{X}$) form:

$$\widetilde{\Delta}_G = \frac{\tilde{X}_{\text{treatment}} - \tilde{X}_{\text{control}}}{\text{MAD}_{\text{control}}}, \qquad \text{MAD}_{\text{control}} = 1.4826 \times \text{median}\left|X_{control_i} - \tilde{X}_{\text{control}}\right|, \qquad i = 1, \ldots, n_c$$

> **Note**
>
> MAD stands for median absolute deviation. In order to make MAD asymptotically consistent with the standard deviation under normality, it is scaled by $C = 1/\Phi^{-1}(0.75) \approx 1.4826$. Implementation-wise, it uses `median_abs_deviation` from `scipy.stats`, and `scale` option is set to `"normal"` to account for the constant multiplier $C$. THE MEDIAN FORM of both Glass's Delta and Cohen's d IS NOT REPORTED IN THE LITERATURE.

> **Important**
>
> Which group is treatment and which is control matters! From the formula of $\Delta_G$, switching the order not only flips the sign of the result, but also changes the denominator, which is $s_{\text{control}}$, the standard deviation of the *control*. Currently, FLIM Playground treats the group on the left as control and the group on the right as treatment (the negative $\Delta_G$ below implies this). In the future, it may support user-customizable x-axis order so that users can arrange the groups to make sure the control group is on the left. Or users can use Cohen's d that does not assume the treatment-control

structure.

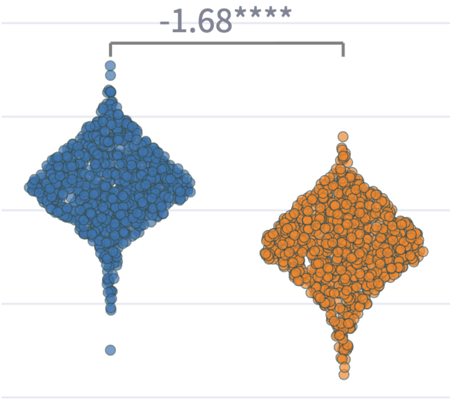

**Note**

To keep the point-based visualization style that supports interactive hover while showing the violin-plot-like distribution, Sina plot is used: it uses `gaussian_kde` from `scikit-learn` to make sure the width of the point distribution is proportional to the kernel density.

##### 13.4.2 Cohen's d

Mean ($\bar{X}$) form:

$$d = \frac{\bar{X}_1 - \bar{X}_2}{s_p}, \qquad s_p = \sqrt{\frac{(n_1 - 1)s_1^2 + (n_2 - 1)s_2^2}{n_1 + n_2 - 2}}$$

Median ($\tilde{X}$) form:

$$\tilde{d} = \frac{\tilde{X}_1 - \tilde{X}_2}{s_{\tilde{p}}}, \qquad s_{\tilde{p}} = \sqrt{\frac{(n_1 - 1)\,\mathrm{MAD}_1^2 + (n_2 - 1)\,\mathrm{MAD}_2^2}{n_1 + n_2 - 2}}$$

#### 13.5 Statistical Test

Statistical tests are performed on the selected comparison pairs that exceed the effect size threshold (if users choose to calculate effect sizes). The statistical tests supported are:

- Independent t-test (student's t-test, for equal variances)

- Welch's t-test (for unequal variances)

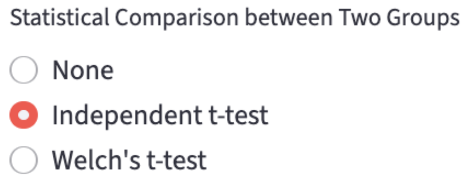

FLIM Playground uses `scipy.stats.ttest_ind` to perform the independent t-test and `scipy.stats.ttest_ind` with `equal_var=False` to perform the Welch's t-test.

#### 13.6 Comparison Annotation

For each selected comparison pair that exceeds the effect size threshold (if any), the statistical significance, effect size, or both are annotated on the plot. It selects the lowest eligible position (over all points in both groups and over existing annotations) to annotate. If both statistical significance and effect size calculation are performed, the annotation is shown in the format of effect size followed by p-value. If only one is performed, then only the performed method's result is shown.

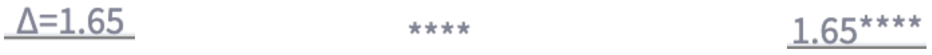

### 14 Feature Histogram

Feature histograms give an instant readout of each group's center, spread, skew and any multi-peaked shape that raw tables hide. FLIM Playground plots a histogram for each group based on the selected numerical feature. Additionally, the Gaussian Mixture Model mode can help find subcomponents of each histogram in an unsupervised way. To quantify the distribution heterogeneity, normalized H-index is calculated for each group.

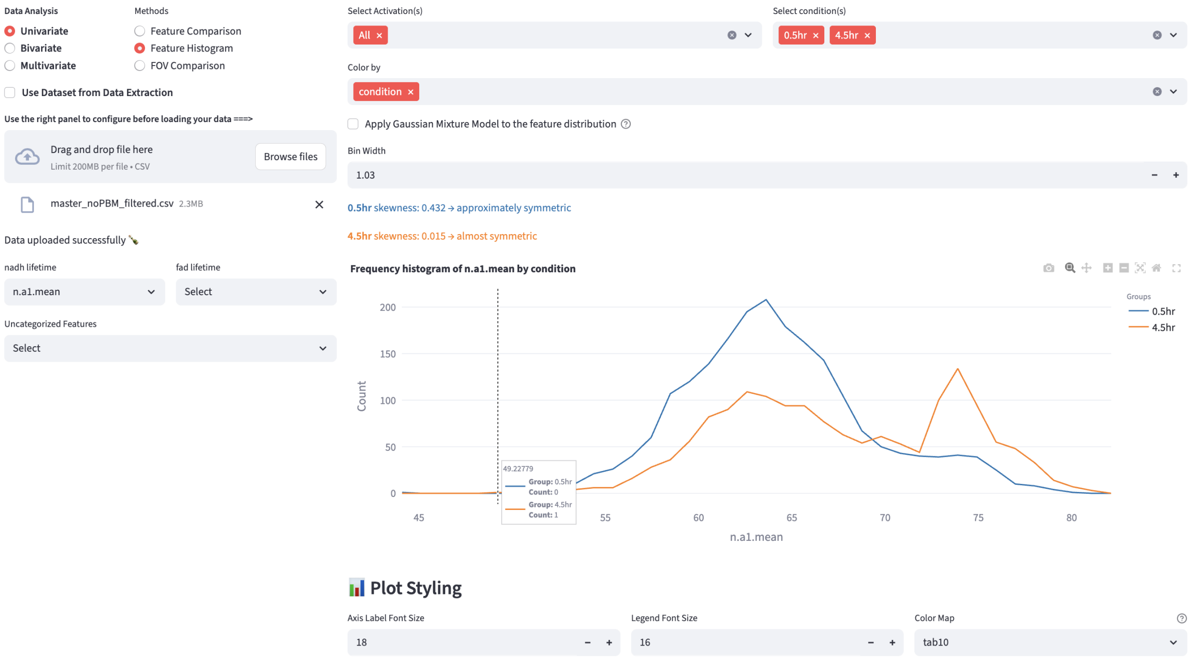

#### 14.1 Shared Interface Components

- On the left, users can use the selection widgets to select the numerical feature to see the histogram/GMM distribution. Exactly one feature can be selected from all feature groups.
- On the top right, users can use the filters to subset the data to find the groups of interest.

- Below the filters, users can apply the visual channels widgets that allow users to `Color by` categorical features. `Opacity by` and `Shape by` are *not* supported because there are no points in the histogram/GMM plot.
- On the bottom right, users can change the plot style using the plot styling widgets.

#### 14.2 Histogram Mode

With a single click, users can switch between the histogram mode and the Gaussian Mixture Model mode. When the checkbox is unchecked, the histogram mode is selected.

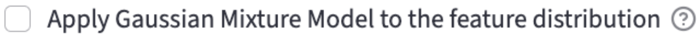

Histogram mode plots a histogram of the selected numerical feature for each group. The histogram bin width is user-controllable, with the default bin width determined automatically as the minimum bin width between the ‘sturges’ and ‘fd’ estimators (see here for more details). The maximum bin width is set to one-third of the range of the data. The step size is set to $1/50$ of the range.

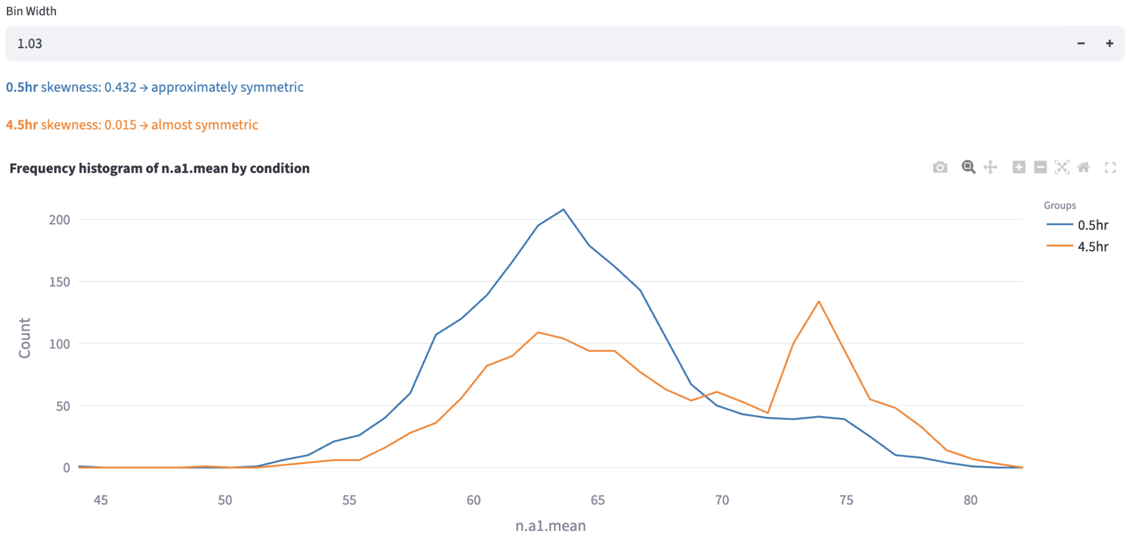

Additionally, the skewness of the distribution is computed and displayed[13].

A rule of thumb categorization of skewness is as follows:

- Strongly left-skewed: < -1
- Moderately left-skewed: -1 to -0.5
- Approximately symmetric: -0.5 to -0.25 and 0.25 to 0.5

- Almost symmetric: -0.25 to 0.25
- Moderately right-skewed: 0.5 to 1
- Strongly right-skewed: > 1

However, the skewness is just one way to quantify the distribution shape. In the example above, the orange distribution was clearly bimodal, but the skewness alone could not capture this. In applications where the distribution is multi-modal, and each mode can represent certain subpopulations (e.g. a cell type, a cell state, a cell cycle phase, etc.), users can use the Gaussian Mixture Model mode to find subcomponents of each distribution in an unsupervised way.

#### 14.3 Gaussian Mixture Model Mode

**GMM Components for 0.5hr:**

| Component | Mean | Std. Dev. | Weight |
|---|---|---|---|
| 1 | 62.18 | 3.51 | 0.68 |
| 2 | 68.67 | 4.66 | 0.32 |

**GMM Components for 4.5hr:**

| Component | Mean | Std. Dev. | Weight |
|---|---|---|---|
| 1 | 63.37 | 3.58 | 0.59 |
| 2 | 73.84 | 2.68 | 0.41 |

H-index for 0.5hr: 0.468. H-index for 4.5hr: 0.485.

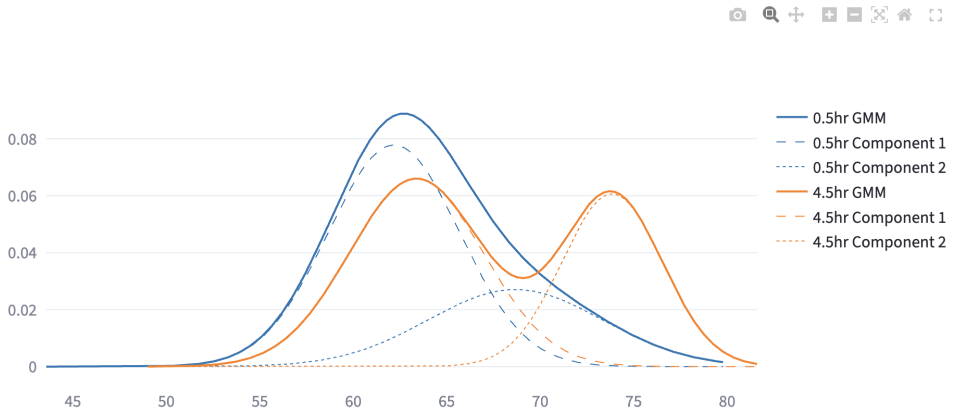

A Gaussian Mixture Model (GMM) represents a dataset as a weighted sum of multiple Gaussian (normal) distributions, each defined by its own mean ($\mu$) and (co)variance ($\sigma^2$). By estimating the component parameters and their weights (mixing coefficients) from the data—commonly using the Expectation-Maximization (EM) algorithm—GMMs can capture complex, multimodal distributions that a single Gaussian cannot.

##### 14.3.1 Fit Gaussian Mixture Models

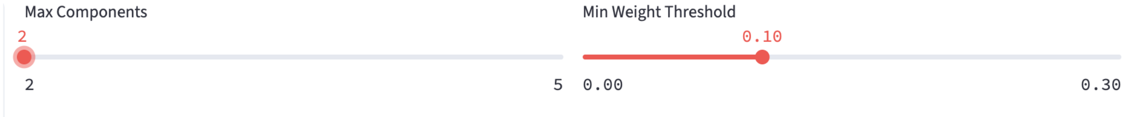

Figure 14.1: Gaussian Mixture Model Configuration

- Fits GMMs on the selected numerical feature up to `Max Components` components, each of which has its component weights ($w_i$), means ($\mu_i$) and variances ($\sigma_i^2$).
- Retains only *valid* models in which every component has weight at least `Min Weight Threshold` (the weights of all components add to 1).
- FLIM Playground will choose the valid model with the lowest BIC score that penalizes the model complexity to avoid overfitting.

##### 14.3.2 Heterogeneity index

To quantify the subpopulation structure, based on the GMM fit, FLIM Playground computes an "H□index" for each group as a weighted entropy□distance: each component contributes according to its weight's uncertainty $(-w_i \log w_i)$[14] scaled by how far its mean $\mu_i$ sits from the overall mixture mean $\bar{\mu} = \sum_{i=1}^{k} w_i \, \mu_i$, normalized by the standard deviation of $\mu_i$s.

$$H = \sum_{i=1}^{k} (-w_i \log w_i) \, d_i, \qquad d_i = \frac{|\mu_i - \bar{\mu}|}{\sigma}$$

##### 14.3.3 Classification

- Users can choose to perform intersection thresholding or hard assignment to determine which GMM component each data point belongs to. It appends a new categorical feature column called `GMM_Group` (recognizable by methods in Data Analysis as a categorical feature) to the filtered dataset, the values of which are `{color_group}_1`, `{color_group}_2`, etc. In the example above, the color groups were created by 0.5hr and 4.5hr because the user colored the data by `hour`. The classification result can be saved by clicking:

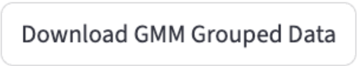

###### 14.3.3.1 Intersection thresholding

- Once the GMM is estimated, thresholds can be obtained by locating the intersection points of adjacent component densities:

$$w_1\,\mathcal{N}(x;\mu_1,\sigma_1^2) \;=\; w_2\,\mathcal{N}(x;\mu_2,\sigma_2^2)$$

  where $w_1, \mu_1, \sigma_1$ are the mixture weight, mean, and standard deviation of the first Gaussian component, respectively, and $w_2, \mu_2, \sigma_2$ those of the second component.
- The above equation can be formed as a quadratic equation and solved for the unique root lying between $\mu_1$ and $\mu_2$. In practice, we bracket this interval and apply a robust one□dimensional root-finding routine (e.g. Brent's method) to compute intersection points. Data points are then assigned to the class corresponding to the interval in which they fall.

> **!** Important
>
> In rare cases when there is more than one intersection point, the data point is assigned using hard assignment rather than intersection thresholding.

###### 14.3.3.2 Hard assignment

- For each data point, it computes the posterior probability (responsibility) of each component, and assigns the data point to the component with the highest responsibility.

$$\text{label}(x) = \arg\max_i \gamma_i(x)$$

$$\text{where} \quad \gamma_i(x) = \frac{w_i\,\mathcal{N}(x \mid \mu_i, \sigma_i^2)}{\sum_{j=1}^{K} w_j\,\mathcal{N}(x \mid \mu_j, \sigma_j^2)}$$

##### 14.3.4 Example

Pham et al. used GMM with hard thresholding to create gates using NAD(P)H $a_1$ and optical redox ratio. These gates were used to define metabolically stressed and fit cells in their study of the impact of cryopreservation on T cell metabolism.

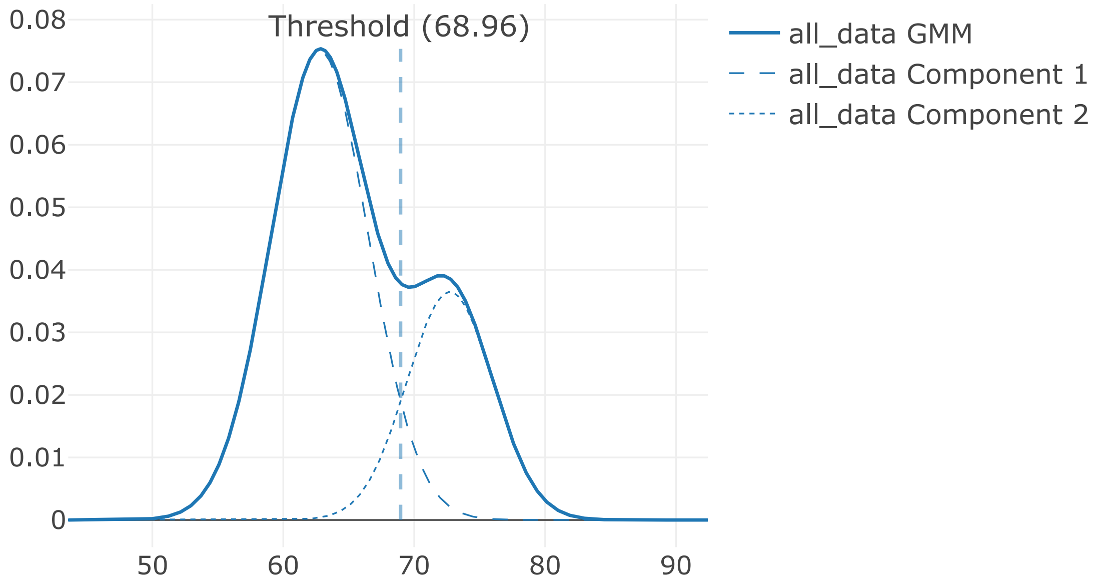
Threshold (68.96)
0.08
0.07
0.06
0.05
0.04
0.03
0.02
0.01
0
50
60
70
80
90
all_data GMM
all_data Component 1
all_data Component 2

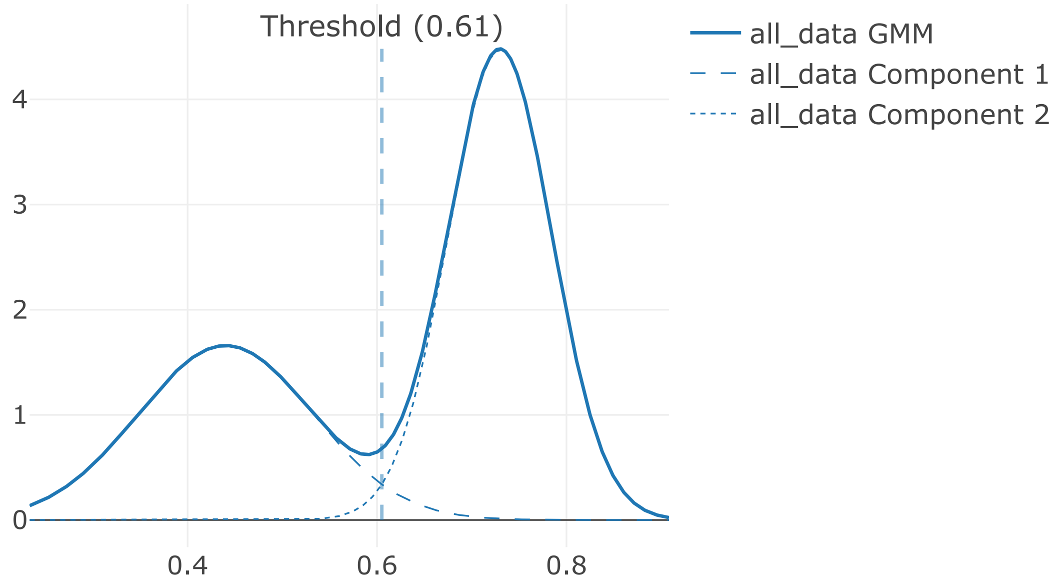

They compared the GMM-derived gate with cell viability by Trypan blue staining:

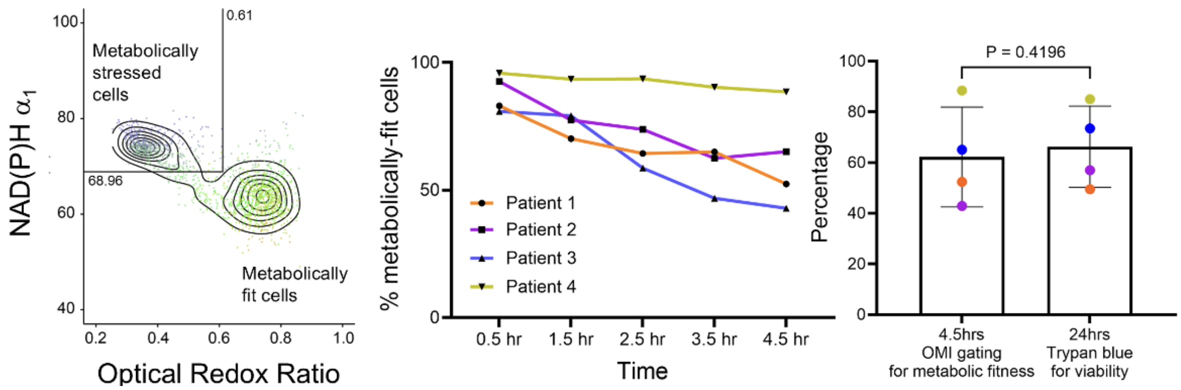

It suggests that optical metabolic imaging (OMI) could serve as an early, non-destructive indicator of T cell viability post-thaw.

### 15 Field of View Comparison

In imaging datasets like FLIM data, field of view (FOV) is an important categorical feature that documents where a certain set of rows (e.g. cells) originate. Visualizing each FOV side by side on a numerical feature can help identify potential FOV-level outliers.

FLIM Playground finds the FOV identifier column from the dataset using the `FOV column name` field in data analysis config (dataset not extracted by Data Extraction) or data extraction config (dataset extracted by Data Extraction).

If the FOV identifier column is not found in the dataset, then a warning message will be shown:

**Warning: We cannot find the fov column from your dataset.**

If found:

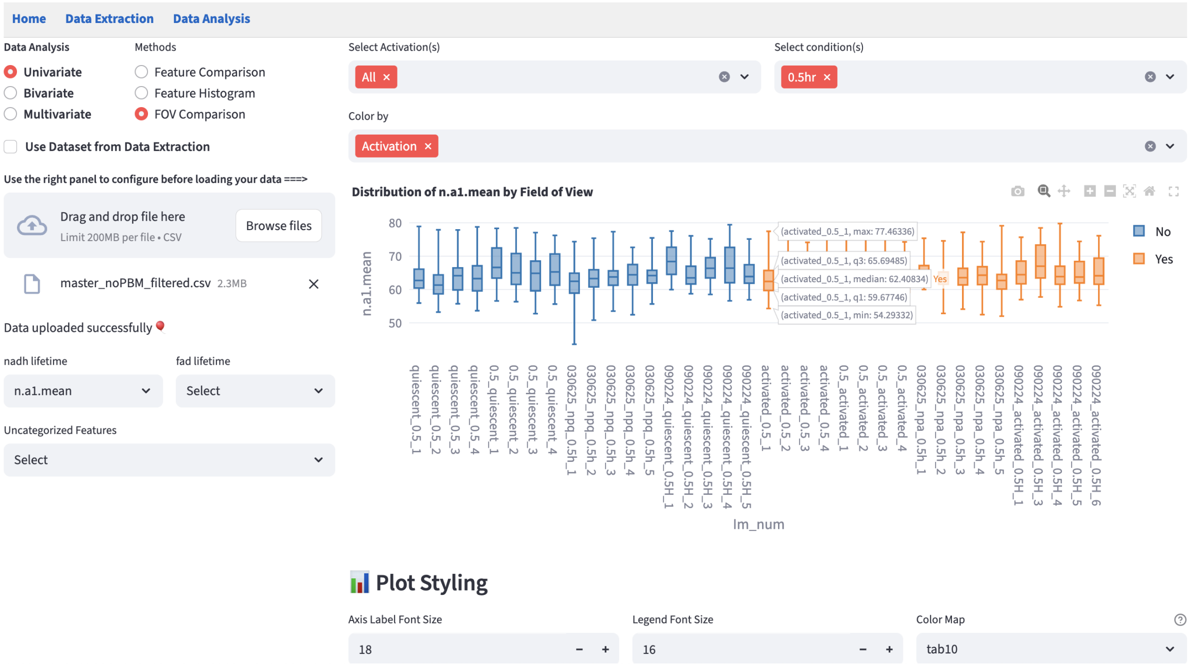

#### 15.1 Shared Interface Components

- On the left, users can use the selection widgets to select the numerical feature to see the boxplot of each FOV. Exactly one feature can be selected from all feature groups.
- On the top right, users can use the filters to subset the data to find the groups of interest.
- Below the filters, users can apply the visual channels widgets that allow users to `Color by` categorical features. `Opacity by` and `Shape by` are *not* supported because there are no points in the boxplot.
- On the bottom right, users can change the plot style using the plot styling widgets.

Besides the shared components, this module does not have any method-specific components.

#### 15.2 Example

Sometimes, the FOV-level boxplots may show consistent divergence across different imaging days, which may suggest that there is experimental or data extraction inconsistency (e.g. shifts are not optimized correctly for one day versus the others). Below is an example that, despite the same conditions, systematically diverged across days, but was consistent within the same day.

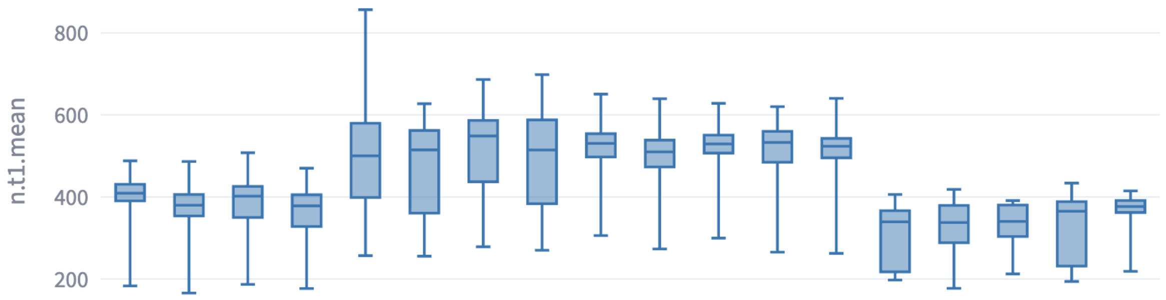

### Part IV

### Bivariate Analysis

### 16 Feature Distribution

This method extends the univariate Feature Histogram method to visualize the distribution of two numerical features on a scatter plot. It helps reveal patterns such as correlations, clusters, trends, or outliers.

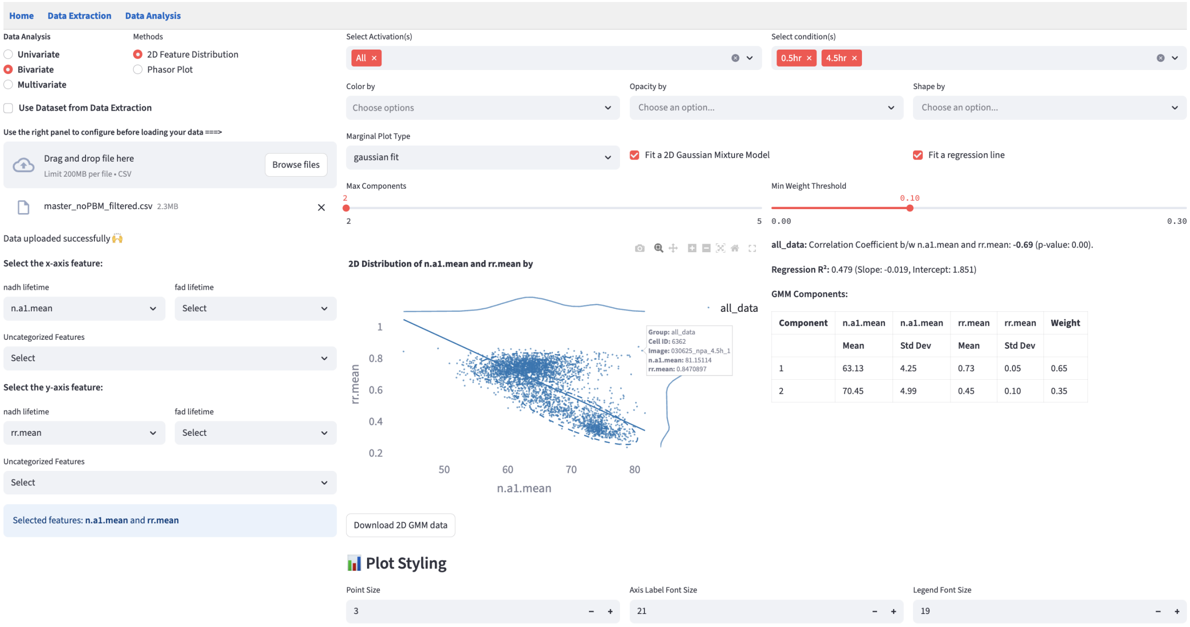

#### 16.1 Shared Interface Components

- On the left, users can use the selection widgets to select the two numerical features on the x-axis and y-axis.
- On the top right, users can use the filters to subset the data to find the groups of interest.
- Below the filters, users can apply the visual channels widgets that allow users to `Color by`, `Opacity by`, and `Shape by` categorical features.
- On the bottom right, users can change the plot style using the plot styling widgets.

#### 16.2 Marginal Distribution

Marginal distributions (the distribution of a single feature) are plotted at the top and the right of the scatter plot. Users can choose the plot type from the dropdown menu.

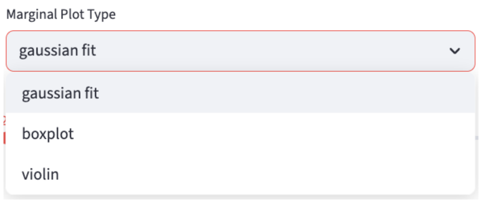

`gaussian fit`: provides a kernel-density estimate using Gaussian kernels. It uses `gaussian_kde` from `scipy.stats` and sets all parameters to default values.

#### 16.3 2D Gaussian Mixture Model

Using the same method as fitting a 1D Gaussian Mixture Model, users can fit a 2D GMM by specifying `Max Components` and `Min Weight Threshold` for each color group.

Ellipses that capture approximately 95% of the points in each subcomponent are drawn. The color of the ellipses is determined by the color group that the subcomponent belongs to.

> **ℹ How ellipses are drawn**
>
> Each ellipse represents a subcomponent of the GMM.
>
> - Center of the ellipse is the mean of the subcomponent $(\mu_x, \mu_y)$.
> - The ellipse is rotated by $\theta = \arctan 2(v_{1y}, v_{1x})$, where $\mathbf{v}_1 = \begin{bmatrix} v_{1x} \\ v_{1y} \end{bmatrix}$ is the first eigenvector of the covariance matrix, so that the major axis of the ellipse is aligned with the eigenvector corresponding to the largest eigenvalue.
> - The semi-axis lengths are $r\sqrt{\lambda_1}$ and $r\sqrt{\lambda_2}$, where $\lambda_1$ and $\lambda_2$ are the covariance eigenvalues (variances along the principal axes) scaled by $r = \sqrt{\chi^2_{\mathrm{df}=2,\, p=0.95}}$ (Mahalanobis distance) that captures approximately 95% of the points in the subcomponent.

All the components below will be rendered for each color group, which is created by user-chosen categorical features in `Color by`.

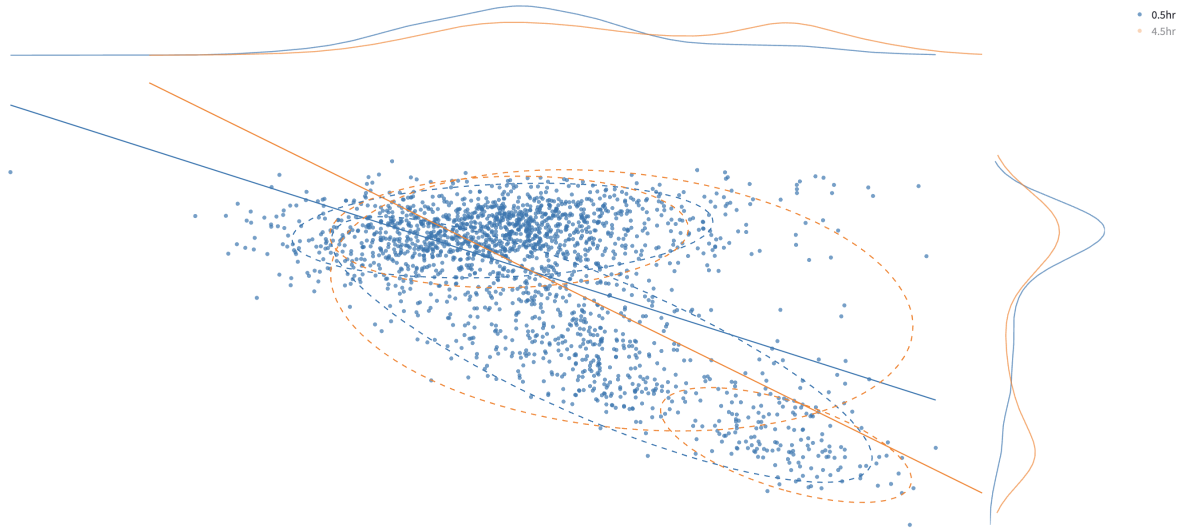

**Tip**

Users can click on a legend group to hide/show the data points in that group. In the above example, to avoid cluttering the plot, the orange group is hidden.

Tables of subcomponents' means, standard deviations, and weights are reported on the side.

**0.5hr:** Correlation Coefficient b/w n.a1.mean and rr.mean: **-0.56** (p-value: 0.00).

**GMM Components:**

| Component | n.a1.mean Mean | n.a1.mean Std Dev | rr.mean Mean | rr.mean Std Dev | Weight |
|---|---|---|---|---|---|
| 1 | 66.71 | 5.29 | 0.54 | 0.11 | 0.36 |
| 2 | 62.83 | 4.12 | 0.74 | 0.04 | 0.64 |

**4.5hr:** Correlation Coefficient b/w n.a1.mean and rr.mean: **-0.76** (p-value: 0.00).

**GMM Components:**

| Component | n.a1.mean Mean | n.a1.mean Std Dev | rr.mean Mean | rr.mean Std Dev | Weight |
|---|---|---|---|---|---|
| 1 | 73.43 | 3.12 | 0.39 | 0.07 | 0.39 |
| 2 | 63.87 | 4.33 | 0.72 | 0.07 | 0.61 |

The classification result can be saved by clicking:

Download 2D GMM data

The classification is done by hard assignment to determine which GMM component each data point belongs to. It appends a new categorical feature column called `2D_GMM_group` to the filtered dataset, the values of which are `{color_group}_group1`, `{color_group}_group2`, etc. The downloaded dataset keeps all the features plus the new categorical feature column, which is recognized by any method in Data Analysis as a categorical feature like others. The rows that pass the filters are included.

#### 16.4 Regression line

A linear regression line is fitted to the data points in each color group. The key statistics are reported on the side.

**Regression R²:** 0.311 (Slope: -0.014, Intercept: 1.570)

$R^2$ tells how much of the variability in $y$ is captured by the model, compared to just predicting the $\bar{y}$.

> **i** $R^2$
>
> - $R^2 = 1 \rightarrow$ perfect prediction.
> - $R^2 = 0 \rightarrow$ model predicts no better than the mean.

### 17 Phasor Analysis

This method focuses on visualizing the distribution of the 1st harmonic or 2nd harmonic phasor coordinates, extracted by the Lifetime fit free extractor, and performs optional clustering to highlight subpopulations or structural patterns.

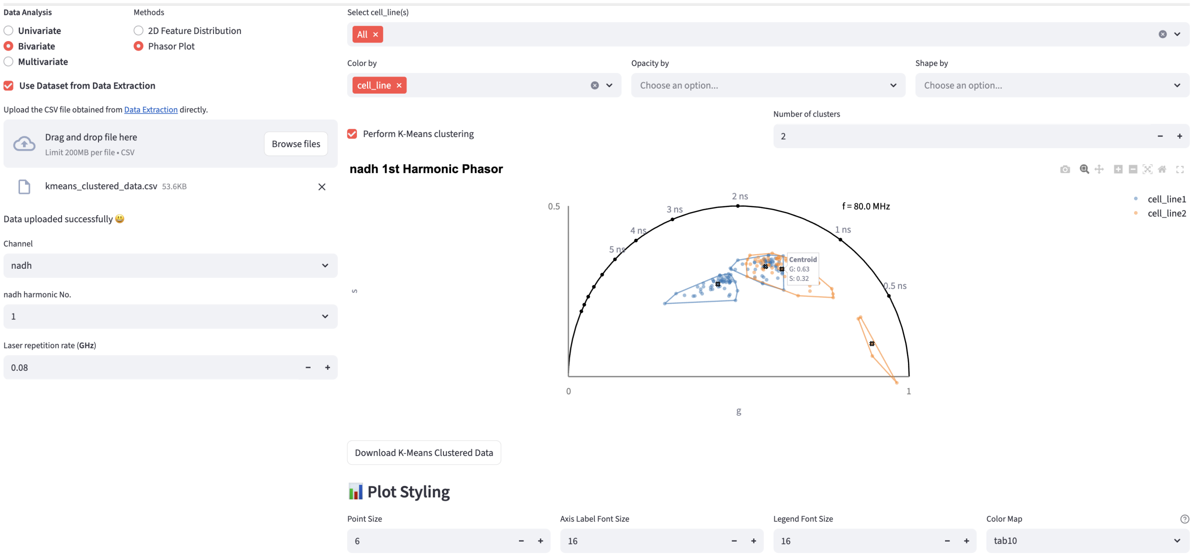

#### 17.1 Shared Interface Components

- On the top right, users can use the filters to subset the data to find the groups of interest.
- Below the filters, users can apply the visual channels widgets that allow users to `Color by`, `Opacity by`, and `Shape by` categorical features.
- On the bottom right, users can change the plot style using the plot styling widgets.

#### 17.2 Select Phasor Coordinate

Instead of using the bivariate selection widgets that select two generic numerical features, this method renders a dropdown menu to select the channel and another to select the harmonic.

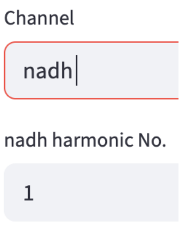

#### 17.3 K-means Clustering

Phasor coordinates of single-cell ROIs from the selected channel and harmonic are plotted in the phasor space. K-means clustering can be performed for each color group (created by `Color by`). If `Perform K-means Clustering` is checked, users can specify the number of clusters to be created based on their prior knowledge. In the future, automatic tuning for this parameter, akin to selecting the number of components of GMM, using silhouette analysis will be added.

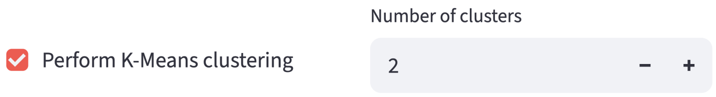

K-means clustering is performed given the number of clusters specified for each color group. Mathematically, it is an iterative algorithm that finds the best cluster assignment with regard to what cluster each point belongs to and what centroid the cluster has. The *best* is defined as the one that minimizes the sum of the squared distances between each point and the center of the cluster it belongs to.

$$\min_{\{C_k\}_{k=1}^{K}} \sum_{k=1}^{K} \sum_{x_i \in C_k} \|x_i - \mu(C_k)\|_2^2, \qquad \mu(C_k) = \frac{1}{|C_k|} \sum_{x_i \in C_k} x_i.$$

First, FLIM Playground standardizes the phasor coordinates because $g$ (real part) and $s$ (imaginary) are not necessarily on the same scale.

Then, it performs k-means clustering using the `sklearn.cluster.KMeans` function. All the other hyperparameters control the optimization process (e.g. `init` controls the initialization of the centroids), and they are set to their default values. `42` is used as the random seed to ensure reproducibility.

To visualize the result of clustering, FLIM Playground draws a polygon (convex hull) that contains all the points of each cluster, the color of which is determined by the color group the cluster belongs to. It uses the `scipy.spatial.ConvexHull` function to calculate the convex hull. The center of each cluster is marked as a black bordered cross and users can hover over it to see its coordinates.

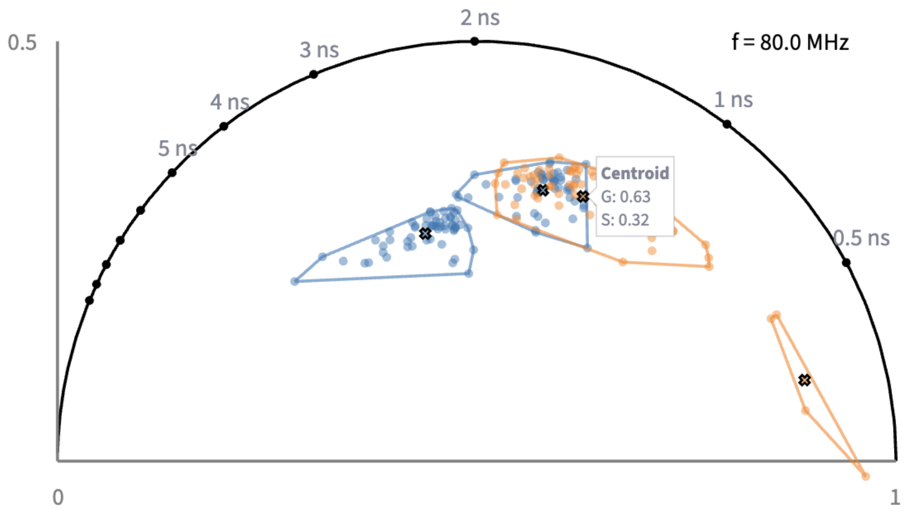

##### 17.3.1 Export Clustered Dataset

Users can download the clustered dataset by clicking:

Download K-Means Clustered Data

The downloaded dataset keeps all the features plus the new categorical feature column (`k_means_cluster`), which is recognized by any method in Data Analysis as a categorical feature like others. The rows that pass the filters are included.

### Part V

### Multivariate Analysis

### 18 Dimension Reduction

To represent and visualize high-dimensional data in a meaningful way and help users interpret the data, dimension reduction methods project the data into a lower-dimensional space. Three popular dimension reduction methods are often used: UMAP, PCA, and t-SNE.

Principal Component Analysis (PCA) is a linear dimension reduction method that projects the data onto the directions of the largest variance. It is fast, deterministic, and easier to interpret (PCA loadings are provided), but it is limited by the linear nature of the projection. For a detailed discussion of PCA, please refer to this post.

Uniform Manifold Approximation and Projection (UMAP) and t-Distributed Stochastic Neighbor Embedding (t-SNE) are non-linear dimension reduction methods that can capture more complex patterns in the data. They are more flexible and can handle non-linear relationships between features, but they are more computationally expensive, less interpretable, not deterministic (FLIM Playground uses the random seed 42 to ensure reproducibility), and dependent on the hyperparameters. Here is a post that explains how UMAP and t-SNE work at a high level and the meaning of some of the hyperparameters.

The numerical features are standardized before the dimension reduction.

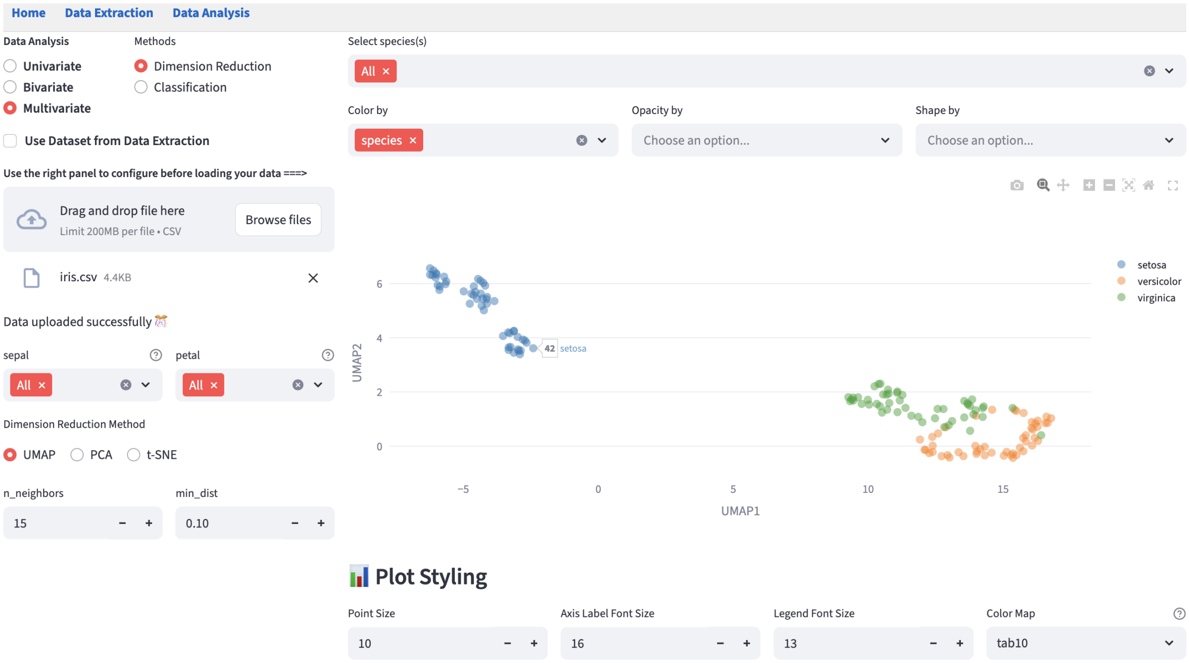

#### 18.1 Shared Interface Components

- On the left, users can use the selection widgets to select multiple numerical features from each feature group.

- On the top right, users can use the filters to subset the data to find the groups of interest.
- Below the filters, users can apply the visual channels widgets that allow users to `Color by`, `Opacity by`, and `Shape by` categorical features.
- On the bottom right, users can change the plot style using the plot styling widgets.

#### 18.2 Hyperparameter Widget

Hyperparameters really matter, so we provide a set of interactive widgets to help users change the hyperparameters and see their effects in real time.

##### 18.2.1 UMAP

FLIM Playground uses the `umap-learn` implementation of UMAP. It chooses to expose `n_neighbors` and `min_dist` for users to tweak. Other hyperparameters are set to default values.

##### 18.2.2 t-SNE

FLIM Playground uses the `sklearn.manifold.TSNE` implementation of t-SNE. It chooses to expose `perplexity` and `early_exaggeration` for users to tweak. Other hyperparameters are set to default values.

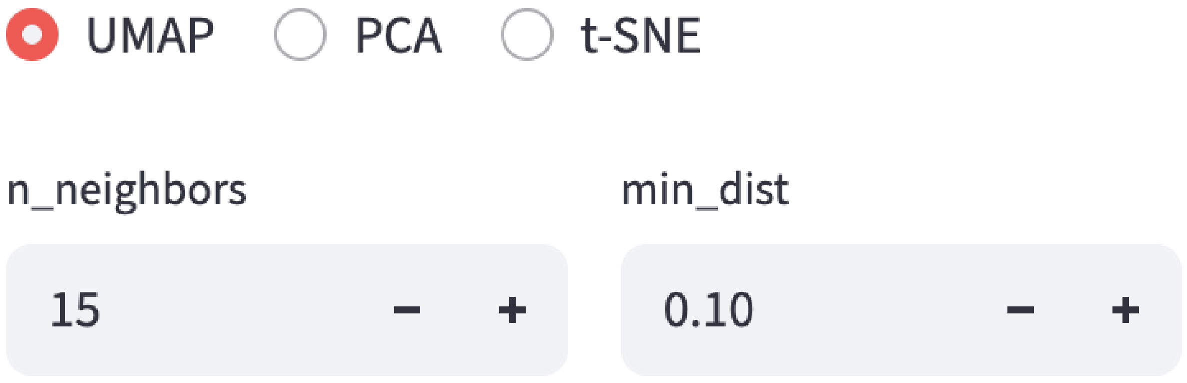

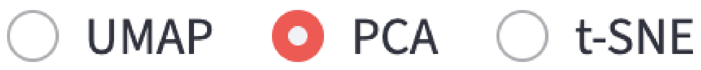
UMAP
PCA
t-SNE

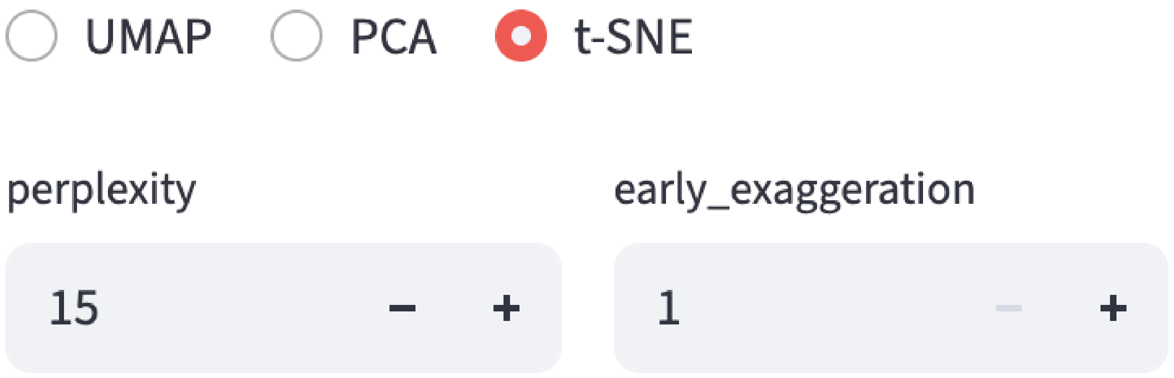
UMAP
PCA
t-SNE
perplexity
15
early_exaggeration
1

### 19 Classification

A machine learning classifier learns patterns from categorized data and then predicts the category of new, unseen data. This module provides a set of machine learning classifiers and interactive widgets to help users perform classification.

All the classifiers are implemented using `scikit-learn` and the performance metrics are calculated using `sklearn.metrics`. A fixed random seed (42) is used for all the classifiers to ensure reproducibility.

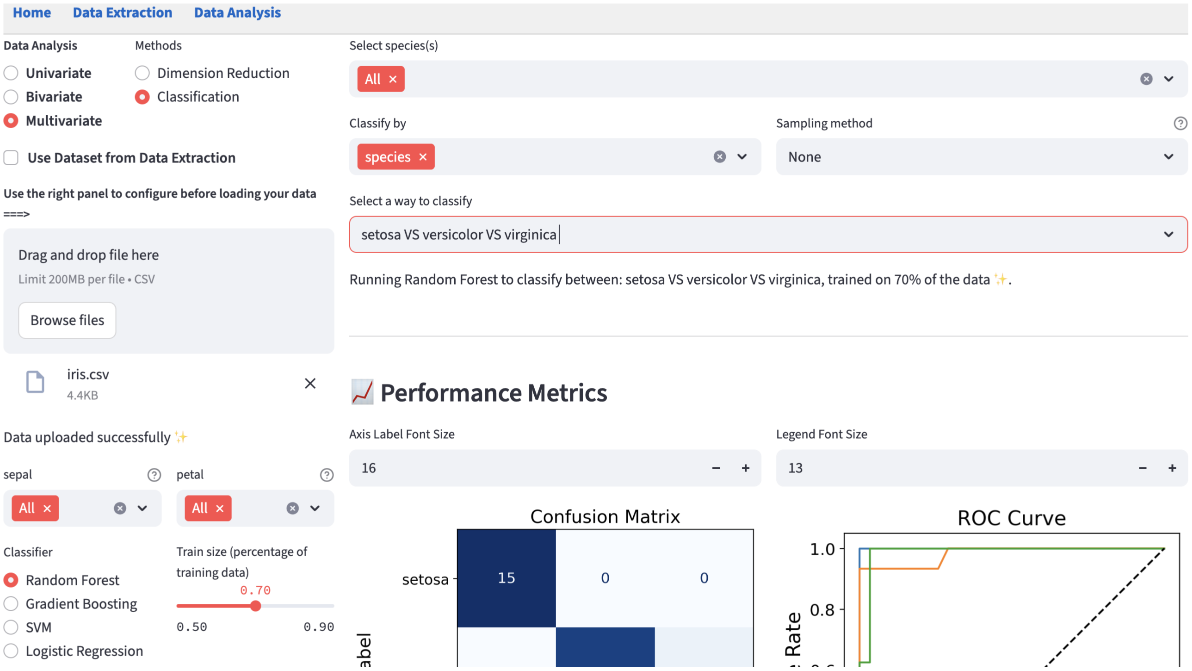

The performance metric plots are cut off in the screenshot. See the performance metrics section for the complete view.

#### 19.1 Shared Interface Components

- On the left, users can use the selection widgets to select numerical features for the classifier. Multiple features can be selected from each feature group.

- On the top right, users can use the filters to subset the data to find the groups of interest.
- Above the performance metrics plots, users can change the plot style using the plot styling widgets.

#### 19.2 Method-specific Components

- Choose the classifier and train-test split: the slider controls the proportion of the data used to train the classifier and the rest is used as unseen data to test its generalizability.

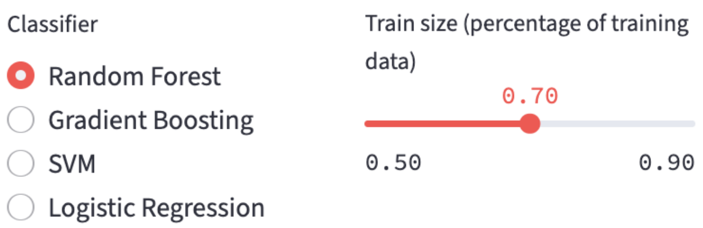

- `Classify by`: The `Color by` from the visual channels widgets is renamed to `Classify by` and it is used to form classes based on (a combination of) selected categorical features. Other visual channels are not supported in this method.

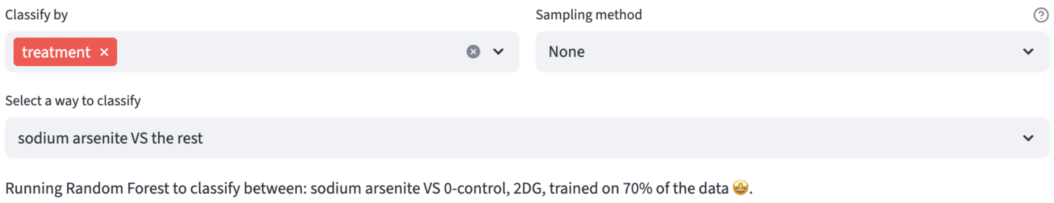

- Choose the classes to be classified: classes are formed based on the combinations of categories of the selected categorical features. In general, it supports binary classification and in general n-way classification where n is the number of classes. If one categorical feature is selected and after filtering has 3 categories, then all possible ways to classify are produced including the three-way classification. Additionally, the `class x vs the rest` option is provided for each class.

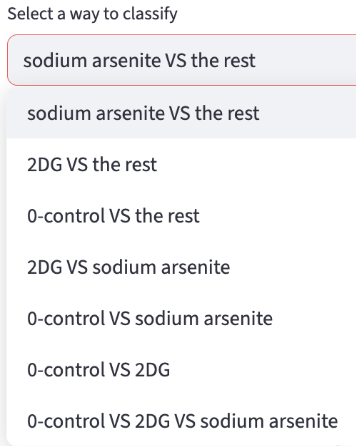

- Sampling methods:
  - `None`: stratified random sampling of the data on classification classes, regardless of the class size. It is implemented using `train_test_split` from `sklearn.model_selection`.
  - `Undersampling`: All classes are downsampled to the size of the smallest class. It is implemented using `RandomUnderSampler` from `imbalanced-learn`.
  - `Oversampling`: All classes are upsampled to the size of the largest class by randomly duplicating samples from minority classes. It is implemented using `RandomOverSampler` from `imbalanced-learn`.

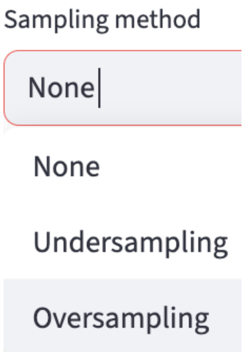

> **Note**
>
> FLIM Playground applies the z-score normalization to the features before training if the chosen classifier is `SVM` or `Logistic Regression`, because they are sensitive to feature magnitudes, while `Random Forest` and `Gradient Boosting` are not.

##### 19.2.1 Performance Metrics

To evaluate the performance of the classifiers, performance metrics visualizations are provided.

###### 19.2.1.1 Confusion Matrix

FLIM Playground evaluates the trained classifier on the held-out test set and summarizes its predictions with `sklearn.metrics.confusion_matrix`. In the binary case, it is represented as:

| | Predicted Positive | Predicted Negative |
|---|---|---|
| **Actual Positive** | **True Positive (TP)** – correct hits | **False Negative (FN)** – missed positives |
| **Actual Negative** | **False Positive (FP)** – false alarms | **True Negative (TN)** – correct rejections |

The accuracy is calculated as the ratio of the sum of the true positives and true negatives to the sum of all the elements in the confusion matrix. Other metrics are calculated for each class based on the confusion matrix:

- Accuracy: $\frac{TP+TN}{TP+FP+TN+FN}$
- Precision: $\frac{TP}{TP+FP}$
- Recall (Sensitivity): $\frac{TP}{TP+FN}$
- Specificity: $\frac{TN}{TN+FP}$
- Youden's Index: Sensitivity $+$ Specificity $- 1$
- F1 Score: $\frac{2\times\text{Precision}\times\text{Recall}}{\text{Precision}+\text{Recall}}$

📈 **Performance Metrics**

**Overall Metrics**

| Metric | Value |
|---|---|
| Accuracy | 0.9742 |
| N | 2520 |

**Per-Class Metrics**

| Class | N | Precision | Recall | Specificity | Youden's J | F1 Score |
|---|---|---|---|---|---|---|
| Cyanide | 1058 | 0.9636 | 0.9754 | 0.9733 | 0.9487 | 0.9695 |
| the rest | 1462 | 0.9821 | 0.9733 | 0.9754 | 0.9487 | 0.9777 |

Axis Label Font Size

18 – +

Legend Font Size

16 – +

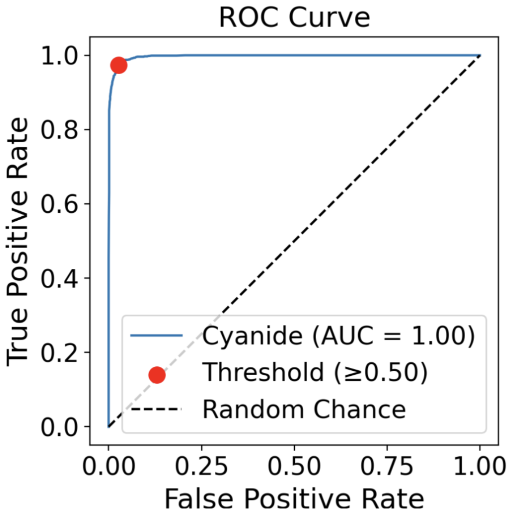

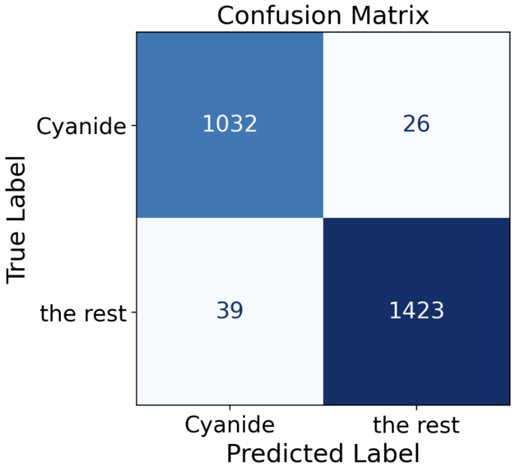

##### 19.2.1.2 ROC Curve

Another way to look at the performance of the classifier is to plot the Receiver Operating Characteristic (ROC) curve and its Area Under the Curve (AUC).

A classifier usually outputs a score for each sample. To turn scores into hard labels, a decision threshold must be chosen. Confusion matrix looks at the performance of the classifier at a single threshold (in the binary case, the threshold is 0.5), while ROC curve looks at the performance of the classifier at all thresholds (from 0 to 1).

At the origin (0, 0), the threshold is 1, so the classifier predicts all samples as negative, making the false positive rate $\mathrm{FPR} = \frac{\mathrm{FP}}{\mathrm{FP+TN}} = 0$ and the true positive rate $\mathrm{TPR} = \frac{\mathrm{TP}}{\mathrm{TP+FN}} = 0$, i.e. the origin. At the point (1, 1), the threshold is 0, so the classifier predicts all samples as positive, making the false positive rate 1 and the true positive rate 1, i.e. the point (1, 1).

AUC summarizes the ROC curve by integrating it from 0 to 1, so it is threshold-free and has a nice interpretation: it is the probability that the classifier assigns a higher score to a randomly chosen positive sample $X^+$ than to a randomly chosen negative sample $X^-$.

To generalize to the $K$-class cases ($K > 2$), FLIM Playground chooses the One-vs-Rest (OvR) strategy. For each class $i$, it trains a binary classifier to predict the class $i$ against all other classes. The ROC curve is then computed and plotted for each class.

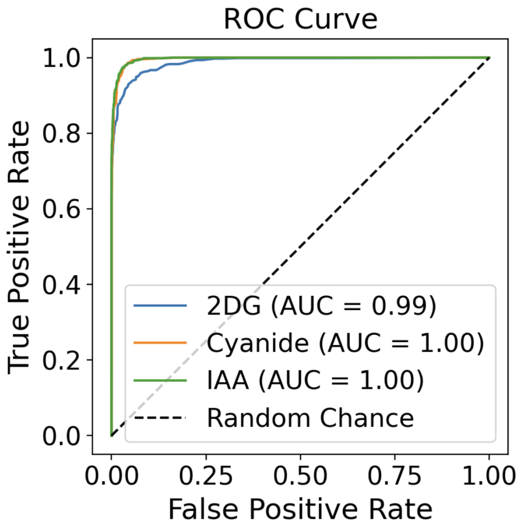

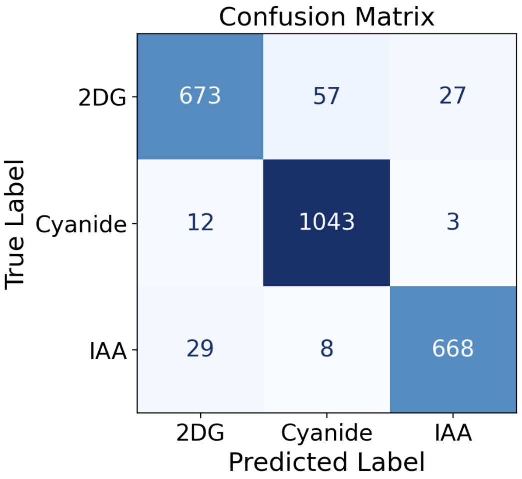

##### 19.2.1.3 Feature Importance

`Random Forest` and `Gradient Boosting` classifiers provide feature importance scores.

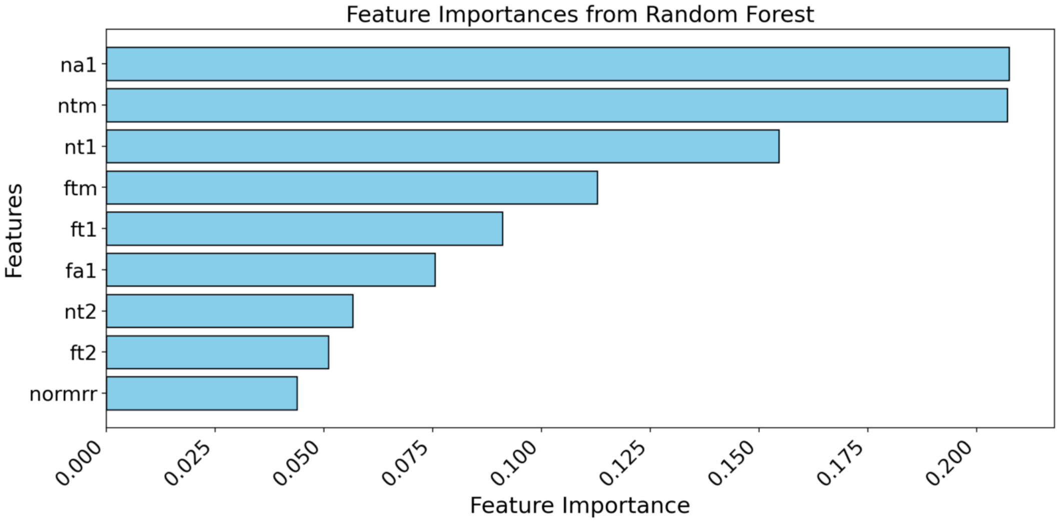

1. Datta, R., Heaster, T. M., Sharick, J. T., Gillette, A. A. & Skala, M. C. Fluorescence lifetime imaging microscopy: fundamentals and advances in instrumentation, analysis, and applications. *Journal of Biomedical Optics* **25**, 071203 (2020).
2. Stringer, C., Wang, T., Michaelos, M. & Pachitariu, M. Cellpose: A generalist algorithm for cellular segmentation. *Nature methods* **18**, 100–106 (2021).
3. Gohlke, C., Pannunzio, B., Schüty, B. & Blanco, R. Phasorpy/phasorpy: v0.7. Zenodo https://doi.org/10.5281/zenodo.16923774 (2025).
4. Gottlieb, D., Asadipour, B., Kostina, P., Ung, T. P. L. & Stringari, C. FLUTE: A python GUI for interactive phasor analysis of FLIM data. *Biological Imaging* **3**, e21 (2023).
5. Kapsiani, S. *et al.* FLIMPA: A versatile software for fluorescence lifetime imaging microscopy phasor analysis. *Analytical Chemistry* (2025).
6. Becker, W. *The Bh TCSPC Handbook*. (Becker & Hickl GmbH, 2021).
7. Stirling, D. R. *et al.* CellProfiler 4: Improvements in speed, utility and usability. *BMC Bioinformatics* **22**, 433 (2021).
8. Stirling, D. R., Carpenter, A. E. & Cimini, B. A. CellProfiler analyst 3.0: Accessible data exploration and machine learning for image analysis. *Bioinformatics* **37**, 3992–3994 (2021).
9. Samimi, K. *et al.* Segmentation-guided photon pooling enables robust single cell analysis and fast fluorescence lifetime imaging microscopy. *bioRxiv* https://doi.org/10.1101/2025.09.30.679660 (2025) doi:10.1101/2025.09.30.679660.
10. Samimi, K. *et al.* Autofluorescence lifetime flow cytometry with time-correlated single photon counting. *Cytometry Part A* **105**, 607–620 (2024).
11. Warren, S. C. *et al.* Rapid global fitting of large fluorescence lifetime imaging microscopy datasets. *PLoS ONE* **8**, e70687 (2013).
12. Cleveland, W. S. & McGill, R. Graphical perception: Theory, experimentation, and application to the development of graphical methods. *Journal of the American Statistical Association* **79**, 531–554 (1984).
13. Virtanen, P. *et al.* SciPy 1.0: Fundamental Algorithms for Scientific Computing in Python. *Nature Methods* **17**, 261–272 (2020).
14. Shannon, C. E. A mathematical theory of communication. *The Bell System Technical Journal* **27**, 379–423 (1948).
